# Magnesium Transporter MgtA revealed as a Dimeric P-type ATPase

**DOI:** 10.1101/2024.02.28.582502

**Authors:** Rilee Zeinert, Fei Zhou, Pedro Franco, Jonathan Zöller, Henry J. Lessen, L. Aravind, Julian D. Langer, Alexander J. Sodt, Gisela Storz, Doreen Matthies

## Abstract

Magnesium (Mg^2+^) uptake systems are present in all domains of life given the vital role of this ion. Bacteria acquire Mg^2+^ via conserved Mg^2+^ channels and transporters. The transporters are required for growth when Mg^2+^ is limiting or during bacterial pathogenesis, but, despite their significance, there are no known structures for these transporters. Here we report the first structure of the Mg^2+^ transporter MgtA solved by single particle cryo-electron microscopy (cryo-EM). Using mild membrane extraction, we obtained high resolution structures of both a homodimeric form (2.9 Å), the first for a P-type ATPase, and a monomeric form (3.6 Å). Each monomer unit of MgtA displays a structural architecture that is similar to other P-type ATPases with a transmembrane domain and two soluble domains. The dimer interface consists of contacts between residues in adjacent soluble nucleotide binding and phosphotransfer regions of the haloacid dehalogenase (HAD) domain. We suggest oligomerization is a conserved structural feature of the diverse family of P-type ATPase transporters. The ATP binding site and conformational dynamics upon nucleotide binding to MgtA were characterized using a combination of cryo-EM, molecular dynamics simulations, hydrogen-deuterium exchange mass spectrometry, and mutagenesis. Our structure also revealed a Mg^2+^ ion in the transmembrane segments, which, when combined with sequence conservation and mutagenesis studies, allowed us to propose a model for Mg^2+^ transport across the lipid bilayer. Finally, our work revealed the N-terminal domain structure and cytoplasmic Mg^2+^ binding sites, which have implications for related P-type ATPases defective in human disease.

## Introduction

The divalent cation magnesium (Mg^2+^) is an essential metal supporting core biological processes including replication, transcription, translation, energy production, protein function and stability, through its interactions with most polyphosphate compounds such as ATP and its roles in enzyme catalysis. In bacteria, Mg^2+^ translocation across the plasma membrane is carried out by the CorA and MgtE families of channel proteins and the CorB/C and MgtA/B families of transporters (reviewed in ^1–3^).

Bacterial CorA and MgtE are constitutively expressed and are required for maintenance of intracellular levels of Mg^2+1^ as are their eukaryotic homologs, members of the MRS2 and SLC41 families, respectively. In bacteria, the loss of CorA results in a reduction in pathogenicity ^4–6^ and in eukaryotes, the loss of Mrs2 impairs mitochondrial function and results in cell death ^7–9^. The role of bacterial MgtE is less well characterized, however, mutations in human SLC41A1 have been linked to cancer and neurodegenerative diseases ^10–12^. Given their implications for bacterial pathogenicity and human health, the structures of CorA and MgtE have been studied extensively ^13–20^. However, aside from mitochondrial MRS2 ^21,22^, there are currently no full-length structures of their eukaryotic counterparts.

When Mg^2+^ becomes less abundant, as during an infection, the homologous MgtA and MgtB transporters (Extended Data Fig. 1) are induced to ensure maintenance of adequate intracellular Mg^2+^ concentrations (reviewed in ^23^). Thus, in *Salmonella enterica*, deletion of *mgtA* and *mgtB* results in sensitivity to the antibiotic polymyxin B, increased cell lysis, decreased survival of bacteria in macrophages and reduced pathogenicity ^24–27^.

MgtA and MgtB are members of the broader family of P-type ATPases responsible for transporting numerous biologically important transition metals including Ca^2+^, Na^+^/K^+^, H^+^/K^+^ and Mg^2+^, several heavy metals such as Co^2+^ and Zn^2+^, as well as lipids (reviewed in ^28^), using ATP hydrolysis to fuel transport. They share a high degree of sequence similarity (Extended Data Fig. 2), particularly of functionally significant residues (Extended Data Fig. 3) and, based on AlphaFold ^29,30^, are predicted to be structurally similar (Extended Data Fig. 4). There are no orthologs of MgtA in vertebrates. Of the vertebrate P-type ATPases, those from the Ca^2+^ transporting clade are the closest to MgtA in sequence. These include the Ca^2+^/Mn^2+^ ATPase transporter ATP2C1A, which is implicated in the skin disease Hailey-Hailey and the extensively studied Ca^2+^ transporter SERCA, which is mutated in Brody’s myopathy (Extended Data Fig. 2 and 4). The mechanism of ion transport by P-type ATPases is described by a Post-Albers cycle where the transporter alternates between the so-called E1 and E2 states (reviewed in ^31,32^). The ATPases are thought to use a highly conserved mechanism of action to coordinate transport of their respective ions across the membrane (reviewed in ^28^). The transition between E1 and E2 states is induced by ligand binding, hydrolysis of ATP, and the transfer of the γ-phosphate of ATP to the aspartate within the conserved DKTGT consensus motif (D373 in *Escherichia coli* MgtA), which distinguishes the P-type ATPases from other ATPases (reviewed in ^33^). For MgtA, a D373N substitution abolished ATP hydrolysis *in vitro,* emphasizing its importance in the catalytic cycle ^34^. Upon metal translocation, the aspartate residue is dephosphorylated by the glutamic acid (E215 in *E. coli* MgtA) from the TGES loop in the A domain, and the protein returns to the E1 state (reviewed in ^35^). The serine in the TGES loop has been shown to be phosphorylated during transport and have a regulatory function in the potassium P-type ATPase KdpFABC ^36^ but is less well conserved. Many insights into P-type ATPase function have been gained through structural work on SERCA with over 70 monomeric structures capturing the various states of its transport cycle (reviewed in ^37^). However, despite the importance of bacterial Mg^2+^ transporters, and the possibility of interfering with the transporter to block infection, there is limited information about their mechanisms of action with only a single structure of the MgtA soluble nucleotide binding domain ^38^.

Recent work has focused on understanding the biochemical properties of MgtA. The *E. coli* MgtA protein (100 kDa, 898 amino acids) is predicted to consist of 10 transmembrane α-helices interwoven with cytosolic regions. The cytosolic regions include an N-terminal tail, a DSβH (double-stranded β-helix fold) domain termed the A (actuator) domain occurring between TM2 and TM3, and a catalytic haloacid dehalogenase domain (HAD) between TM4 and TM5. The HAD is comprised of the P (for phosphorylation) subdomain, as it bears the reversibly phosphorylatable aspartate, and the cap subdomain, termed N (for nucleotide), as it binds the sugar and base moieties of the nucleotide and excludes water during phosphotransfer (Fig. 1 and Extended Data Fig. 1) ^39^. Using *in vitro* ATP hydrolysis as a proxy for MgtA transport activity, recent work showed MgtA is regulated by both Mg^2+^ levels and the presence of the anionic lipid cardiolipin ^34^. MgtA colocalizes with cardiolipin *in vivo* suggesting lipid composition is important for MgtA localization and activity ^34^. In both *E. coli* and *S. enterica*, the stability of the transporter is regulated by small proteins of less than 50 amino acids ^27,40^. The structural basis for how Mg^2+^ concentration and lipid composition modulate MgtA activity and how small proteins regulate the degradation of the transporter is unclear.

**Fig 1.**
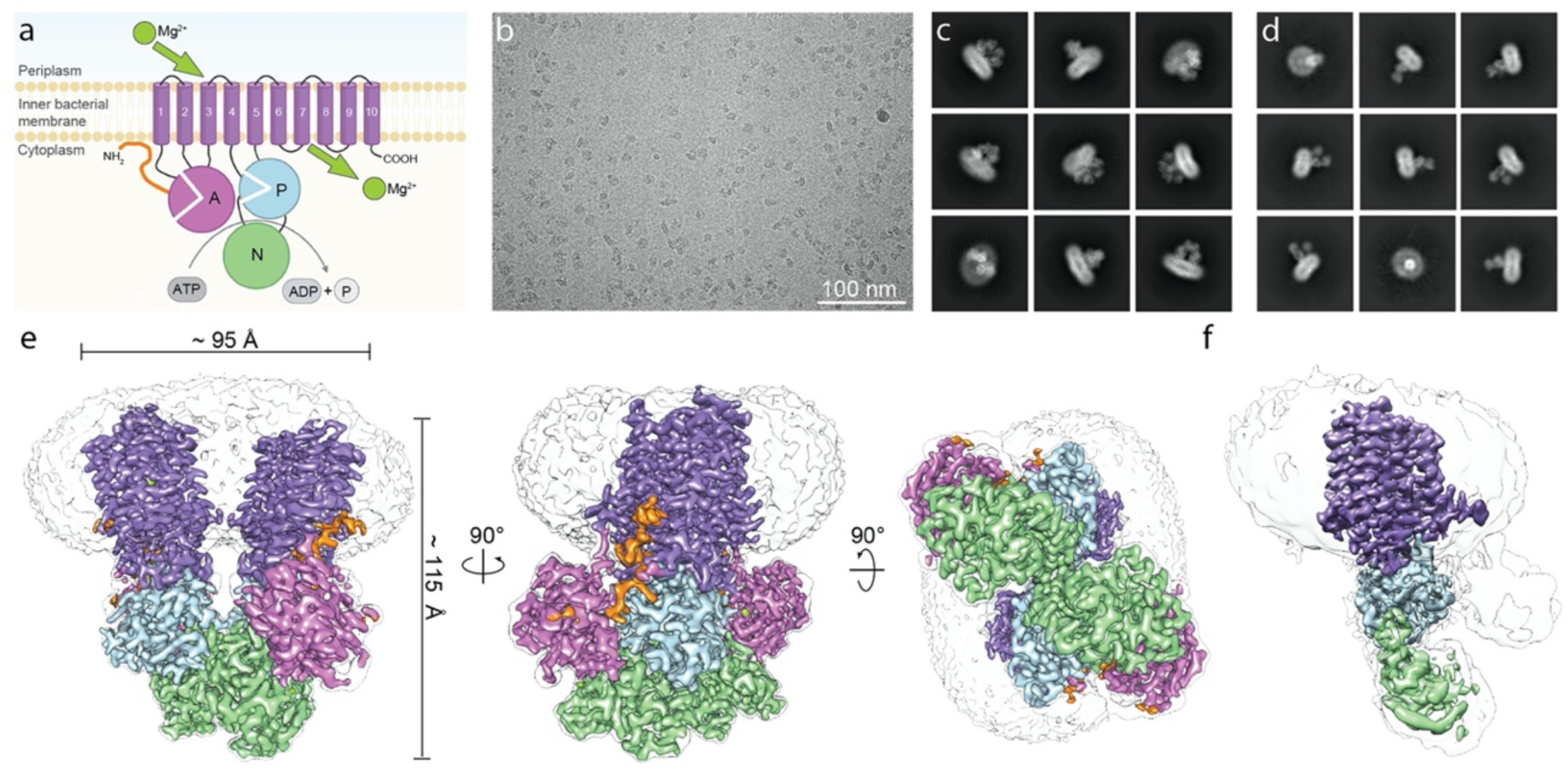
Cryo-EM of magnesium transporter MgtA reveals a high-resolution dimer and a monomeric structure. **a**, Schematic representation of MgtA/B based on P-type ATPase structural homology showing ten conserved transmembrane helices colored in purple (1-10), the actuator (A) domain in orchid, the phosphorylation (P) subdomain in light blue, the nucleotide (N) binding or CAP subdomain in light green, and the predicted unstructured N-terminal tail in orange. The A domain is split into two regions a/b. The b segment of the A domain is comprised of a <u>D</u>ouble <u>S</u>tranded <u>beta</u>-<u>H</u>elix fold (DSβH). The soluble P subdomain is a noncontiguous segment comprised of two regions a/b that house the key catalytic residues required for phosphorylation. The N subdomain binds the nucleotide and aids in catalysis. The P and N subdomains together comprise the <u>H</u>alo<u>a</u>cid <u>d</u>ehalogenase (HAD) domain. **b**, Representative micrograph of purified *Escherichia coli* MgtA. **c**-**d**, Representative final 2D class averages for the ∼200 kDa dimer (**c**) and the ∼100 kDa monomer (**d**), respectively, with a box size of 384 pixels (approximately 319 Å). **e**-**f**, Cryo-EM reconstruction of the MgtA dimer (**e**, also see Extended Data Fig. 9a and Extended Data Movie 1) and of the monomer (**f**, also see Extended Data Fig. 9b and Extended Data Movie 2). Cryo-EM maps in **e**-**f** are colored as in **a**. A transparent gray map at a much lower threshold indicates the detergent micelle and the density for the more flexible loops and the A domain in the monomer map.

Here we used single-particle cryo-electron microscopy (cryo-EM) to determine the first structure of the bacterial protein MgtA at an overall resolution of 2.9 Å, illuminating several aspects of the transporter. Surprisingly, we found that MgtA exists as both a dimer and a monomer when purified using mild detergents. A 3.6 Å structure of the monomer showed significant rearrangements of the transmembrane α-helices as well as an overall open state of the cytosolic domains when compared to the compact dimer structure. The N-terminus, found in MgtA and the H^+^/K^+^ and Na^+^/K^+^ ATPase transporters, previously predicted to be disordered, was captured in the dimer structure. MD simulations in a lipid environment together with sequence conservation and a co-purification assay support the conclusion that the dimer is a stable structure that may be a broadly conserved property of P-type ATPase transporters. Cryo-EM structures of the MgtA dimer bound to the ligands ATP, ATPγS, and ADP, at resolutions of 3.7, 3.9, and 3.8 Å, respectively, identified residues that facilitate ATP binding. The cryo-EM maps also showed a strong density for a Mg^2+^ ion in the middle of TM α-helices 5, 7 and 8, which, based on sequence homology predictions, is predicted to be a novel Mg^2+^ ion binding site. Two additional Mg^2+^ ion binding sites were resolved between the cytosolic domains. Conservation analysis combined with mutagenesis studies revealed residues required for Mg^2+^ ion selectivity, transport and possibly regulation. Based on our collective data, we propose a model for Mg^2+^ trafficking from the periplasm through the TM α-helices to the bacterial cytosol and regulatory roles for other features of MgtA. Given the functional conservation found within the P-type ATPase family, these features likely are also relevant to other transporters in this family.

## Results

### Structural analysis reveals dimeric and monomeric forms of MgtA

*E. coli* MgtA was previously shown to bind and be stabilized by the 31-amino acid MgtS protein ^40^. We set out to purify the MgtA-MgtS complex from cells co-overexpressing MgtA-MgtS. Detergent solubilization of *E. coli* membranes containing both overexpressed proteins revealed that weaker detergents such as glycol-diosgenin (GDN) maintained MgtA-MgtS interactions while stronger and more commonly-used detergents such as n-dodecyl-ß-D-maltoside (DDM) and lauryl maltose neopentyl glycol (LMNG) disrupted the hetero-complex (Extended Data Fig. 5). Intriguingly, LMNG and DDM-solubilized MgtA migrated at ∼200 kDa by Native-PAGE, while GDN-solubilized MgtA migrated at ∼400 kDa suggesting MgtA may exist in two distinct oligomeric states (Extended Data Fig. 5). For structural studies we proceeded with MgtA-MgtS purified with GDN in the presence of 5 mM MgCl_2_ given that it preserved native protein-protein interactions (Extended Data Fig. 6).

Negative-staining EM initially showed that the purified sample was homogenous, containing individual particles of roughly the expected size of ∼8-11 nm (Extended Data Fig. 7). Cryo-EM images of purified MgtA (Fig. 1b) revealed particles of sizes between ∼7-11 nm including typical membrane protein side views. 2D class averages showed particles with larger (Fig. 1c) and smaller (Fig. 1d) detergent micelles, indicating the location of the transmembrane region of the protein and suggesting the protein existed in two states. During data processing, it became clear that the larger particles represented a dimeric state of the magnesium transporter MgtA, while the smaller particles corresponded to a monomeric state, consistent with the two forms of the transporter detected by Native-PAGE (Extended Data Fig. 5 and 6) as well as by negative-staining EM (Extended Data Fig. 7). The characteristic A and HAD (containing N and P subdomains) soluble domains were visible for some side view 2D class averages for the smaller particles (Fig. 1d). The final dimer maps obtained here had an overall resolution of 2.9 Å (C2) and 3.0 Å (C1) (Fig. 1e, Extended Data Fig. 8-10, Extended Data Movie 1) with local resolutions ranging from ∼2.2 to ∼3.5 Å (Extended Data Fig. 9). The final monomer map had an overall resolution of 3.65 Å, with local resolutions ranging from ∼2.8 to ∼7 Å. The lowest resolution corresponded to the A domain, indicating a high degree of flexibility of that region (Fig. 1f and Extended Data Fig. 8 and 9, Extended Data Movie 2). Interestingly, none of the maps revealed densities for the small protein MgtS. We thus focused on features of MgtA and P-type ATPases more broadly.

We observed extra densities corresponding to detergent micelles in the raw EM images and 2D class averages (Fig. 1b, c and d and Extended Data Fig. 7) and the cryo-EM maps (Fig. 1e-f and Extended Data Fig. 8 and 11a). Upon increasing the threshold of the dimeric EM density map, elongated densities, corresponding to detergents or co-purified lipids were observed directly adjacent to the TM α-helices indicating the location of the inner and outer lipid membrane leaflet (Extended Data Fig. 11a). However, as is quite common in membrane protein structures, the local resolution was insufficient to assign a molecular identity. The hydrophobic regions of the TM segments and the border of the inner lipid membrane leaflet could be discerned from the electrostatic potential surface representation of the final structural model of MgtA as well as by the hydrophobic region of the bilayer in the MD simulation (Extended Data Fig. 11b). MD simulations in conditions that mimic the native lipid environment also revealed several arginine and lysine residues that were in prolonged contact with the headgroups of anionic lipids (Extended Data Fig. 11c-d). Interestingly, cardiolipin (highlighted in green), shown to regulate MgtA and colocalize with the transporter *in vivo* ^34^, was found to migrate toward the transporter during the simulation (Extended Data Fig. 11e).

To complement our structural analysis, we treated GDN-solubilized MgtA with a zero length crosslinker (Extended Data Fig. 12a) and identified cross-linked residues by mass spectrometry (Extended Data Fig. 12b). Based on the dimeric structure, the distances between the crosslinked residues ranged from 3.5 to 45.5 Å. The crosslinks of <10 Å were consistent with nearby residues in our final dimeric reconstruction (Extended Data Fig. 12c-d). Most of the long range crosslinks (>20 Å) were between the end of TM α-helix 2 (K144) and the A domain or P subdomains and could indicate that GDN-solubilized MgtA has multiple conformational states, especially since the A domains of P-type ATPases undergo large structural rearrangements during the transport cycle. However, it is possible that some of the crosslinks occurred between neighboring molecules. Taken together, these data support our final reconstruction of MgtA and suggest the existence of additional structural states.

### The MgtA monomer has a more open conformation than the dimer

To identify structural changes, we superimposed the monomer MgtA onto the dimer structure and visualized changes by RMSD and morphing (Extended Data Fig. 13 and Extended Data Movie 3). The A domain displayed the greatest degree of difference, but several other significant differences were also observed. First, the TM1 and TM2 α-helices moved by ∼1.5 helix turns towards the cytoplasm (Extended Data Fig. 13e), possibly because they are pulled by the mechanical forces of the A domain rotation. Second, TM4, which harbors E331, a well conserved residue involved in coordinating ions in other P-type ATPases, appears to unwind allowing E331 to rotate (Extended Data Fig. 13f). The rotation of E331 may make it more available for ion binding/unbinding in the monomer. Third, the A domain and P and N subdomains all move further away from each other, adopting a more open structure (Extended Data Fig. 13c, g-i). The extended configuration of the cytosolic domains in the open Mg^2+^-bound monomer structure resembles the SERCA Ca^2+^-bound state (PDB 1SU4 ^41^).

### Structural architecture of the N-terminus of MgtA suggests a regulatory role

The overall subunit architecture is highly similar among P-types ATPases, but MgtA (residues 1-36) and both the Na^+^/K^+^ and H^+^/K^+^ ATPases have poorly conserved N-terminal tails (Extended Data Fig. 14a). For the Na^+^/K^+^ and H^+^/K^+^ ATPases, the N-termini have been proposed to be regulatory elements though the details remain obscure ^42^. Previous *in vitro* studies showed that, in isolation, the N-terminus of MgtA is intrinsically disordered and interacts with anionic lipid vesicles suggesting this domain might act as a membrane tether ^43^. AlphaFold predictions of MgtA and other P-type ATPases show that the structural prediction of this segment is uncertain, consistent with the N-terminus being disordered or requiring other protein contacts for stabilization (Extended Data Fig. 4). However, we observed well-resolved density of the N-terminus in our MgtA dimer map (Fig. 1e and Extended Data Fig. 10). The N-terminus covers a negatively charged surface spanning the TM2 and TM4 α-helices and interacts with cytosolic A and P regions (Extended Data Fig. 14b, c). The resulting structural model is dramatically different from the AlphaFold prediction (Extended Data Fig. 14d). Attempts to truncate or mutate the N-terminus resulted in decreased stability of the transporter preventing functional analysis (Extended Data Fig. 14e). This suggests that several electrostatic interactions might drive or stabilize the N-terminus in the conformation we observe and that this element is required for the stability of the transporter. Salt bridges are present between the N-terminus (R11) and TM2 (E139) and TM4 (E331), and between N-terminus (R16) and TM2 (D147) (Extended Data Fig. 14f). We also observed several interdomain contacts; between the N-terminus (D21) and the base of TM2 (K150) and the A domain (Q84), as well as between the N-terminus (R20) and the P subdomain (D670) (Extended Data Fig. 14f). Interestingly, the N-terminus appears to snorkel under the A domain forming a knot-like structure. Given the unique configuration and interdomain contacts, we hypothesize that the N-terminus may help stabilize the structural state and have a regulatory function since it interacts with residues such as E331, which is thought to be involved in ion translocation. Simulations show that the N-terminus adopts a stable, short amphipatic α-helix at the leaflet surface, consistent with the hydrophobicity of I5 and L9 and charge of K3 and R8, interacting with nearby lipid phosphate moieties (Extended Data Movie 4). This region is poorly conserved; nevertheless, it is consistently present and tends to be amphipathic across the MgtA-like clade of Mg^2+^ transporters. We suggest that while there is no constraint for specific interactions, the N-terminus likely plays a regulatory role as a flexible element that dynamically associates with the above-mentioned negatively charged surface.

### The MgtA dimer is stable and likely conserved in closely related P-type ATPases

The observation of an MgtA homodimer by cryo-EM suggests other members of the P-type ATPases may also form stable dimers. Early studies of the SERCA Ca^2+^ transporter, one of the closest mammalian homologs of MgtA, showed that SERCA can oligomerize and is influenced by both detergents and lipid membrane composition ^44–51^. Furthermore, recent work on the SERCA pump suggests that dimerization takes place *in vivo* ^52^ and is required for function ^53,54^. The plasma membrane calcium ATPases (PMCA) also have been reported to be regulated by dimerization (reviewed in ^55^). However, aside from the fungal plasma membrane proton transporter Pma1, which was solved as a hexamer ^56^, no homo-oligomeric P-type ATPase structures have been reported.

The dimer interface of MgtA is comprised of salt bridges (K382/E582, K548/E549), hydrogen bonding between polar residues (Q380/Q380 and N387/Q508) and hydrophobic contacts (V384, L385, L510, L544, P546, P547) of the N and P subdomains of neighboring MgtA subunits (Fig. 2a, Extended Data Fig. 1 and Extended Data Table 1). The TM domains do not appear to be involved in dimerization since the closest distance between the two subunits in the TM region is ∼8 Å (between I839 of TM9 of each monomer). While MD simulations cannot easily assess the absolute stability of an interface, they can indicate the dynamic nature of interactions, likely correlating with stability. Interaction partners Q380-Q380, K548-E549, and K382-E582 were all closely positioned and persistent throughout the simulation. Our simulations also revealed an additional salt bridge (E386-K578) that was not present in our structure. The hydrophobic interface was intact through the entire simulation (Extended Data Movies 5 and 6). The dimer interface residues are well conserved among MgtA and MgtB family members and to a lesser extent in the mammalian homolog SERCA, but not in more distantly related P-type ATPase proteins (Extended Data Fig. 2 and 3). These data suggest that dimerization is an important feature for at least a subset of these proteins.

**Fig 2.**
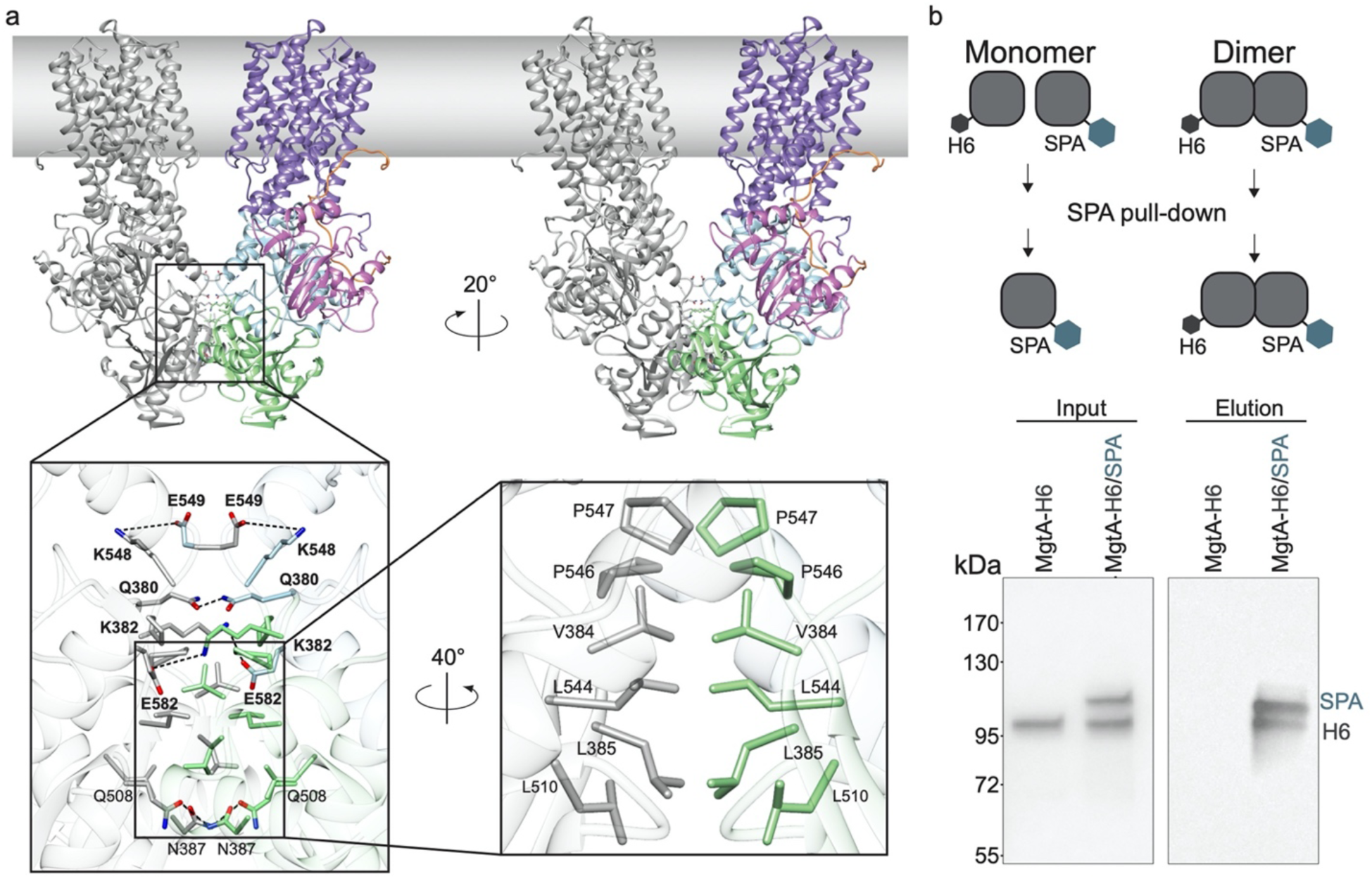
The dimer interface is formed by both hydrophobic and polar interactions. **a**, Side view of the overall dimer structure with the left monomer in gray and the right monomer colored as in Fig. 1. A close-up view of the extensive dimer interface between the two N and P subdomains with sidechain residues displaying charge interactions across the dimer interface. Rotation of the structure highlights hydrophobic interactions at the dimer interface. Molecular dynamic simulations (see Extended Data Movies 5 and 6) show consistent interactions across the dimer interface between K382-E582 (64%), Q380-Q380 (87%), and K548-E549 (88%) shown in bold. **b**, Co-purification of two differentially tagged MgtA derivatives, one tagged with His6 and the other tagged with the larger SPA tag as shown in the top schematic reveals copurification of the two proteins (right elution panel). Proteins were visualized by Western blot analysis using MgtA antibodies.

To test if oligomerization of MgtA could be observed *in vivo*, we co-expressed MgtA with a small tag (6XHis) and MgtA with a large tag (SPA comprised of a calmodulin binding domain and a FLAG epitope) (Fig. 2b). As a control for non-specific interaction with the FLAG resin we included a sample only expressing the smaller MgtA-H_6_. Consistent with stable dimer formation *in vivo*, we found that when we purified MgtA-SPA from cells expressing both proteins the elution fraction contained both MgtA-SPA and MgtA-H_6_ at a nearly equal ratio, which was not present in the MgtA-H_6_ negative control samples (Fig. 2b).

We next sought to test whether dimerization was required for MgtA function by introducing several mutations along the dimer interface (Extended Data Fig. 15a). We assessed the ability of MgtA dimer mutants to complement an *E. coli* strain, which is deficient in all endogenous Mg^2+^ uptake systems and requires supplementation with MgSO_4_ or an active Mg^2+^ transporter for growth ^57^. Under low magnesium conditions (0 and 1 mM MgSO_4_), wild-type MgtA was capable of supporting growth under both non-inducing and inducing conditions (Extended Data Fig. 15b and Extended Data Table 2). However, all the dimer-interface mutants were at least partially defective for complementation despite being present at detectable levels (Extended Data Fig. 15b-c and Extended Data Table 2). Even with the high number of mutations, dimerization was still observed for the various mutants in co-IP assays (Extended Data Fig. 15d). The reduced complementation could be due to slightly lower mutant protein levels or native dimeric contacts are required for maximal activity. Together the data support the conclusion that MgtA adopts a stable dimeric structure.

### The MgtA dimer can bind ATP

We next asked whether the dimer would bind ATP and whether nucleotide binding would lead to conformational changes compared to the nucleotide-free structure. MgtA fractions containing predominantly dimer (Extended Data Fig. 6b) were mixed with 5 mM ATP, non-hydrolysable ATPγS, or ADP prior to grid preparation. The cryo-EM maps for the samples containing Mg^2+^ and the nucleotides all show extra density at the expected and conserved nucleotide binding sites (Extended Data Fig. 16 and 17). While our maps were only of intermediate resolution (3.72 Å, 3.87 Å and 3.75 Å for the ATP-, ATPγS-, and ADP-bound forms, respectively) probably due to the lower protein concentration, thicker ice and lower particle numbers, we did not detect large structural rearrangements when comparing our nucleotide free (apo) map to our nucleotide bound maps (Extended Data Fig. 16c).

The ATP binding site, resolved locally at 2.9-3.2 Å (Fig. 3a-c and Extended Data Fig. 16 and 17a-b, and Extended Data Movie 7) is consistent with other P-type ATPase nucleotide binding pockets (Extended Data Fig. 18). Unique to the core MgtA-MgtB clade, including EcMgtA, is a second phenylalanine, F447, near the F445 residue that is more broadly conserved in the nucleotide binding subdomains of P-type ATPases. F447 makes additional contacts with the adenine ring of the ATP (Fig. 3b), while in most other structures the adenine moiety interacts only with F445 (Extended Data Fig. 18). MD simulations sampled a range of Mg-ATP conformations in our MgtA-ATP structure (Extended Data Fig. 19). In both ATP binding pockets of the simulated dimer, the adenine moiety interacts with F445 and N415 before a major conformational change moves the nucleotide toward R616 after 1.24 and 1.9 microseconds, while the phosphate tail stayed bound.

**Fig 3.**
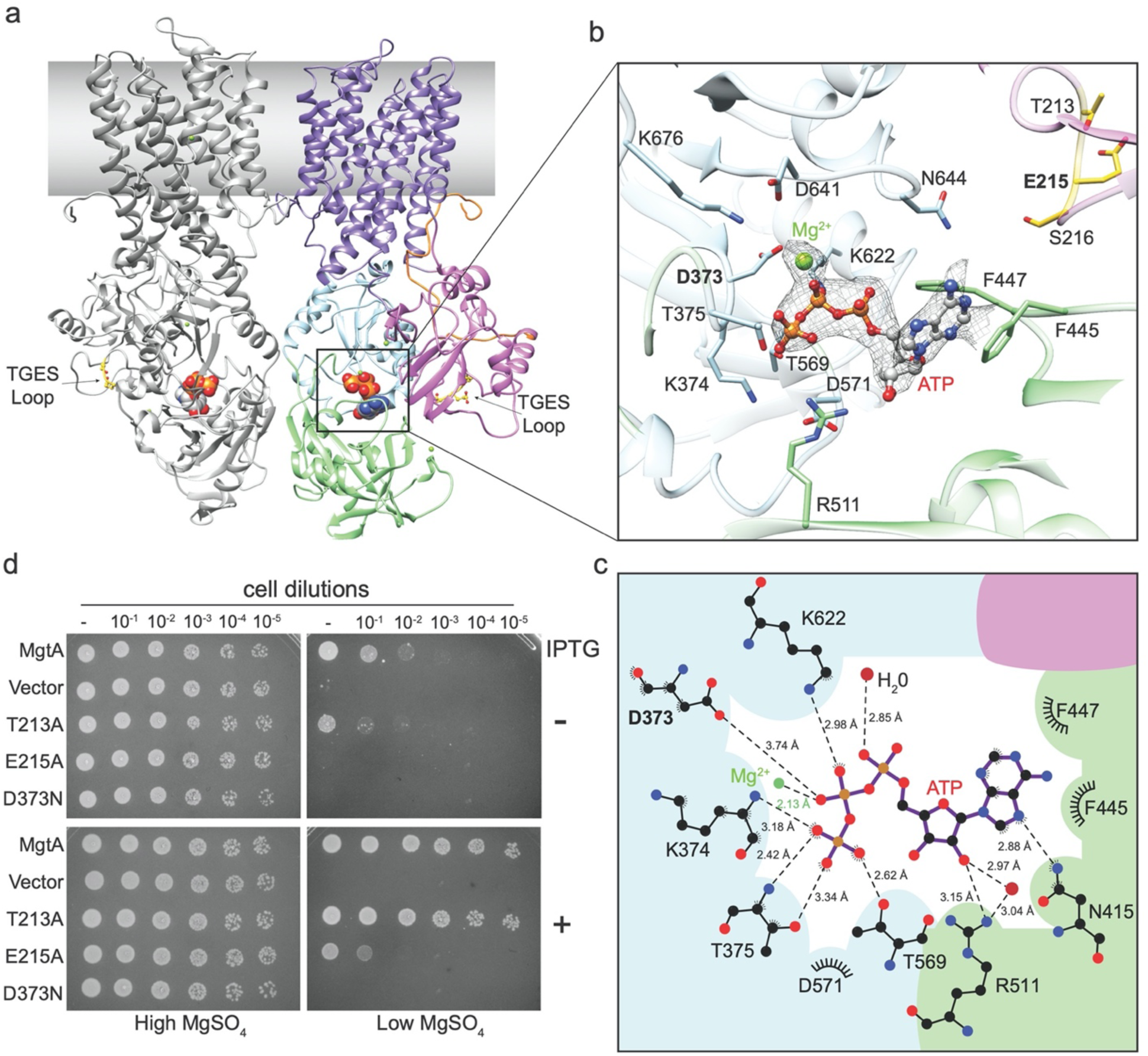
The nucleotide binding pocket of MgtA is accessible in the dimeric state. **a**, Side view of the MgtA dimer with the left monomer in gray and the right monomer colored as in Fig. 1, highlighting the ATP molecule represented in spheres located in between the soluble A domain and P and N subdomains and the dephosphorylation TGES loop in yellow. **b**, A close-up view of the ATP binding site highlighting residues in close proximity to the Mg-ATP molecule which is shown in a ball and stick representation and the cryo-EM map density in gray mesh. Residues from the TGES loop involved in dephosphorylation and located in the A domain are colored yellow. **c**, Distances from ATP to amino acids, water molecules and Mg^2+^ ion in the nucleotide binding pocket of MgtA from *E. coli*, determined using LIGPLOT+ ^88^. **d**, MgtA_D373N_ is unable to completement a Mg^2+^-auxotrophic *E. coli* strain indicating this mutant transporter does not translocate Mg^2+^ ions. D373 is the residue that is being phosphorylated upon ATP hydrolysis. MgtA_E215A_ is only partially able to complement upon overexpression. E215 is part of the TGES loop involved in dephosphorylation. Overnight cultures were serial diluted and spotted onto LB agar plates supplemented with high (100 mM) or low (1 mM) MgSO_4_ with (+) and without (−) 0.1 mM IPTG for induction and grown at 37°C (also see Extended Data Fig. 21).

MgtA with ATPγS, though resolved at slightly lower resolution, was structurally similar to MgtA with ATP (Extended Data Fig. 16 and 17). Some extra density in the nucleotide binding pocket is observed in the final map of the MgtA dimer in the presense of ADP, but the density was much weaker and not sufficiently clear to model ADP into the pocket. This indicates that some ADP can enter the nucleotide binding pocket but cannot be bound as tightly as ATP or ATPγS. Collectively, these data show that the dimer can bind the nucleotides similar to other P-type ATPases.

To complement our single particle studies, we carried out Hydrogen Deuterium Exchange Mass Spectrometry (HDX-MS), which can report on regions able to exchange hydrogen with deuterium, to assess structural changes upon ATP binding. To achieve the high protein concentration required for HDX-MS, we solubilized the protein with DDM, which gives 10-fold higher yields but results in predominantly monomeric MgtA (Extended Data Fig. 5). MgtA was incubated without (control) or with (binding) ATP or ATPγS. Uptake differences for binding versus control were then plotted onto our dimer and monomer structures for visualization (Extended Data Fig. 20). For both ATP and ATPγS, the most dramatic differences in uptake are in the N and P subdomains, which are required to directly coordinate ATP and exclude water from the hydrolysis reaction. Decreased uptake is observed in several peptide stretches containing residues required for ATP binding. Outside of the segments directly involved in ATP binding, we observed decreased uptake for several TM helices that contain conserved ion binding site residues required for translocation in related P-type ATPases, possibly reflecting interactions with Mg^2+^ ions during the translocation. We observed fewer changes in the TM segments when ATPγS was used, suggesting that the TM α-helices undergo less binding or indicate that the transport is stalled when unable to hydrolyze ATP. We suggest that the reduction in uptake in the cytosolic domains is the result of a transition to a more compact structure upon ATP binding. This compaction is similar to what is observed for the dimer structure compared to the monomer structure (Fig. 2, Extended Data Fig. 13, Extended Data Movie 3) and is also seen for SERCA ^58^. Alternatively, MgtA may adopt a conformation which excludes water from these sites. Taken together, the data suggest the monomer and dimer are both capable of ATP binding and the monomer, upon binding ATP, adopts a more compact overall architecture potentially similar to that of the dimer state.

To assess the *in vivo* function of the DxT motif and the TGES dephosphorylation loop, we introduced a mutation at D373 (MgtA_D373N_), which was previously shown to abolish ATP hydrolysis *in vitro* ^34^ and alanine substitutions at T213 and E215 in the TGES loop (MgtA_T213A_ and MgtA_E215A_). The mutants were again tested for their ability to complement the Mg^2+^ uptake deficient strain. Under low magnesium conditions (0 and 1 mM MgSO_4_) wild-type MgtA and MgtA_T213A_ were capable of supporting growth in both non-inducing and inducing conditions (Fig. 3d and Extended Data Fig. 21). In contrast, MgtA_D373N_ and MgtA_E215A_ had significantly less growth without MgSO_4_ supplementation (Fig. 2d), despite being stably expressed (Extended Data Fig. 21b). These data show D373 within the DxT motif is essential and E215 of the dephosphorylation loop has a significant role in the MgtA transport mechanism.

### Multiple Mg^2+^ binding sites are detected in the MgtA dimer

Our cryo-EM maps for the dimeric structure of MgtA show extra densities corresponding to three Mg^2+^ ions (denoted I, II and III for referencing purposes), likely captured during the purification in the presence of 5 mM MgCl_2_ (Fig. 4a, Extended Data Fig. 10). Site I is comprised of residues D780, N709, S776, and S813 residing in transmembrane helices 5, 7, and 8 along with two molecules likely to be water (Fig. 4a). MD simulations of the MgtA dimer show the ion is stably bound at this site, with the ion tightly bound by D780 (Extended Data Movie 8). Sequence conservation analysis comparing the MgtA clade and the SERCA Ca^2+^ clade revealed that residue D780 is exclusively present in the magnesium transporter clade (Extended Data Fig. 22). These data suggest that D780 is a key nodal position that may help with ion capture or selectivity. An assessment of sequence conservation and structural similarity revealed two additional transmembrane binding sites, comprised of residues E331 and D738, that are well conserved in other P-type ATPases (Extended Data Fig. 2 and 22). Consistent with the importance of E331 and D780 residues, alanine substitutions at these positions prevented complementation of the transport deficient mutant (Fig. 4b and Extended Data Fig. 23). A MgtA_D738A_ mutant, detected at wild-type levels, could only partially complement in the absence of induction, though higher levels of the mutant protein did restore growth (Extended Data Fig. 23c). These observations suggest the mutant has slightly reduced capacity for translocation, but is not completely inactive.

**Fig 4.**
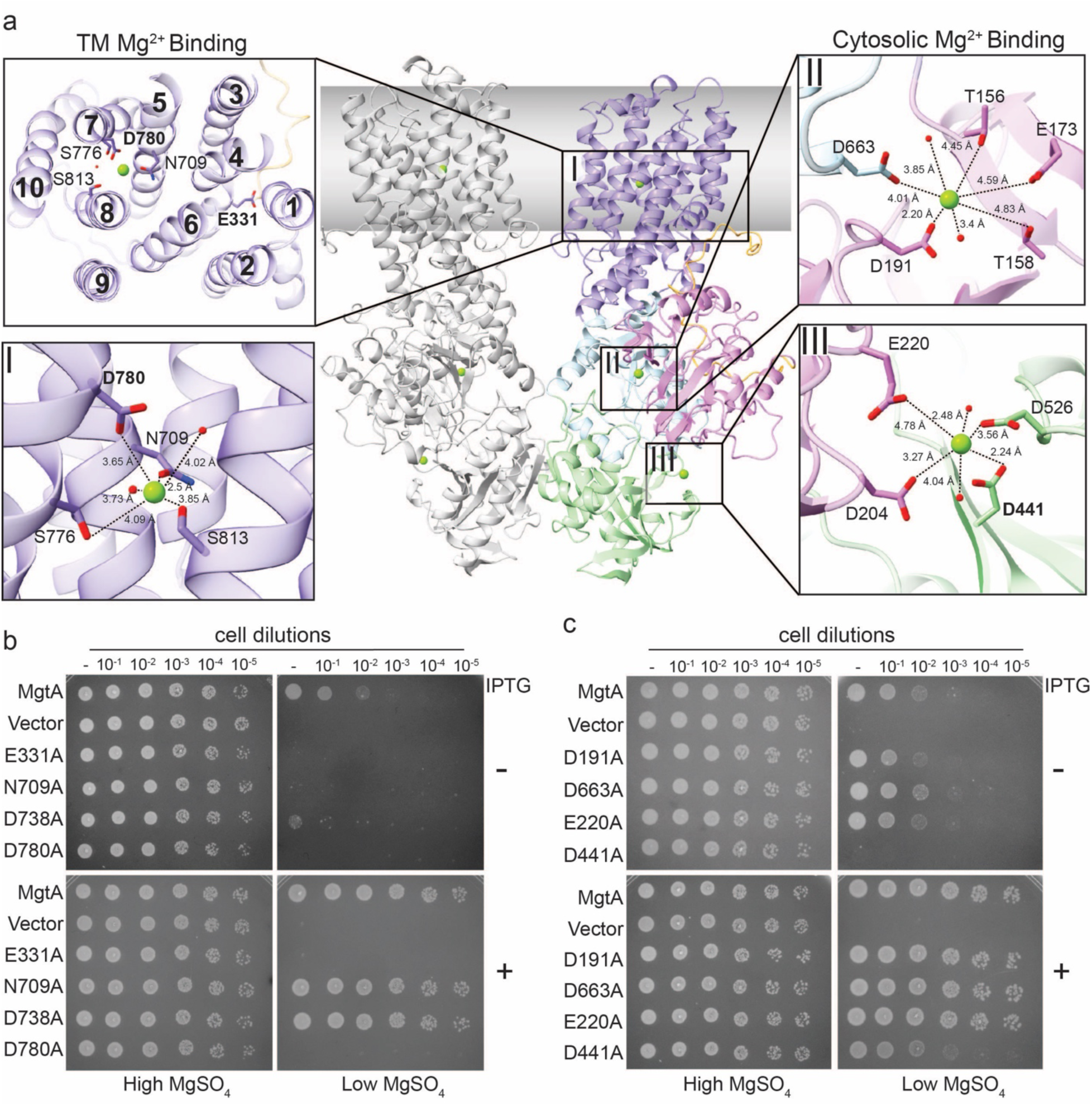
Mg^2+^ binds sites near the middle of the transmembrane domain and between the cytosolic A domain and N and P subdomains. **a**, Side view of the overall dimeric structure with the left monomer in gray and the right monomer colored as in Fig. 1 with close-up views of the resolved Mg^2+^ ions and the nearby residues and proximal resolved water molecules (small red spheres). Site I is located within the TM α-helices 5, 7, and 8. Site II is comprised of residues from the A domain and P subdomain. Site III is comprised of residues from the A domain and N subdomain. **b**, MgtA_E331A_ (located in TM4) and MgtA_D780A_ (located in TM7) are unable to complement a Mg^2+^-auxotrophic *E. coli* strain. **c**, MgtA_D441A_ (site III) is only partially able to complement the Mg^2+^-auxotrophic *E. coli* strain upon overexpression. Complementation assays in **b** and **c** were carried out as for Fig. 3d (also see Extended Data Fig. 23).

We next looked for the presence of a solvated path that could support the hydration of the Mg^2+^ ion during transport by tracing resolved waters in our structure and present in our simulations (Extended Data Fig. 24). A snapshot of the TM region after one microsecond of simulation time shows that this timeframe is sufficient to hydrate the TM interior. A solvated path is created near the periplasmic side consisting of residues H729 and Q733 in TM6 and W806 in TM8 and continues to interior residues S705 and N709 in TM5 and D780 in TM8 (site I), as well as closer to the cytoplasmic side near residues T702 and N706 in TM5 and Q741 in TM6, including a large solvated chamber bounded by residues N734 and D738 in TM6 and N99, N102, T106 in TM1, S131 in TM2, and E331 in TM4 (Extended Data Fig 24).

Soluble domain sites II and III are comprised of residues in the cytosolic A domain and P and N subdomains of the protein. Site II is formed between P subdomain residue D663, and A domain residues D191, T156, T158, and E173 with densities for nearby water molecules (Fig. 4a, Extended Data Fig. 10). Site III is made of N subdomain residues D441 and D526, and A domain residues D204 and E220, and neighboring water molecules (Fig. 4a, Extended Data Fig. 10). To assess the functional importance of the closest residues, we introduced alanine mutations at site II (MgtAD191A and MgtAD663A) and site III (MgtAE220A and MgtAD441A) and assessed their ability to support growth of the Mg^2+^ uptake deficient strain. Site II mutants, MgtAD191A and MgtAD663A, and site III mutant, MgtA_E220A_, were all capable of supporting growth under low Mg^2+^ concentrations (Fig. 4c and Extended Data Fig. 23). Intriguingly, mutant MgtAD441A displayed a significant reduction in growth under low Mg^2+^ with and without inducer (Fig. 4c and Extended Data Fig. 23). Given the residues that comprise site III are located near F445 and F447 that directly interact with ATP and residues D204 and E220 that lie on either side of the TGES loop it is possible this site impacts the ATP hydrolysis cycle.

## Discussion

Here we present the first high-resolution structure of the P-type ATPase Mg^2+^ transporter MgtA, in a dimeric state bound to Mg^2+^. Additional structures of dimeric MgtA bound to ATP, ATPγS, and ADP resolved several residues involved in nucleotide binding, but unlike what has been observed for all monomeric P-type ATPase structures, the cytosolic domains did not show large structural changes upon nucleotide addition. This work also provides the first structure of the N-terminal domain, not well predicted by AlphaFold, found in some P-type ATPase proteins. Additionally, we documented a well-defined Mg^2+^ density coordinated by residues within TM α-helices 5, 7, and 8. Mutational analysis of this site as well as others based on sequence conservation resulted in impaired growth under Mg^2+^ limiting conditions and thus revealed residues involved in ion translocation across the membrane. We also identified two cytosolic Mg^2+^ binding sites, which based on mutational analysis, might function in a regulatory manner. These data provide the first structural insights into the function of the important MgtA family of bacterial Mg^2+^ transporters and emphasize the importance of mild membrane extractions of membrane proteins. Given the conservation among the P-type ATPase family of transporters, our work also has broader implications.

### Model for path of Mg^2+^ translocation

Functional conservation in the TM segments, which are more divergent among P-type ATPases than the rest of the protein, in combination with our newly solved structure, the results of our MD simulations, and mutational analyses, allow us to propose a working model for the path Mg^2+^ travels through MgtA across the membrane (Fig. 5a). In this model Mg^2+^ ions are funneled in from the periplasmic facing residues H729 and Q733 of TM6 and W806 of TM8. The ions are specifically selected for by residues S705 and N709 of TM5, S776 and D780 of TM7, S813 and Q814 of TM8, which are under a strong evolutionary constraint in the MgtA clade but not in the SERCA clade. These residues are either proximally to or directly coordinate the Mg^2+^ in our structure; a D780A mutation abolished all transport activity. Of note, residue G773 in TM7 is proximal to residues S776 and D780 (directly coordinating the Mg^2+^ ion) and is likely critical for mobility during trafficking of the ion. A comparable residue that induces a kink has been observed in the Na^+^/K^+^ ATPase structure ^59,60^.

**Fig 5.**
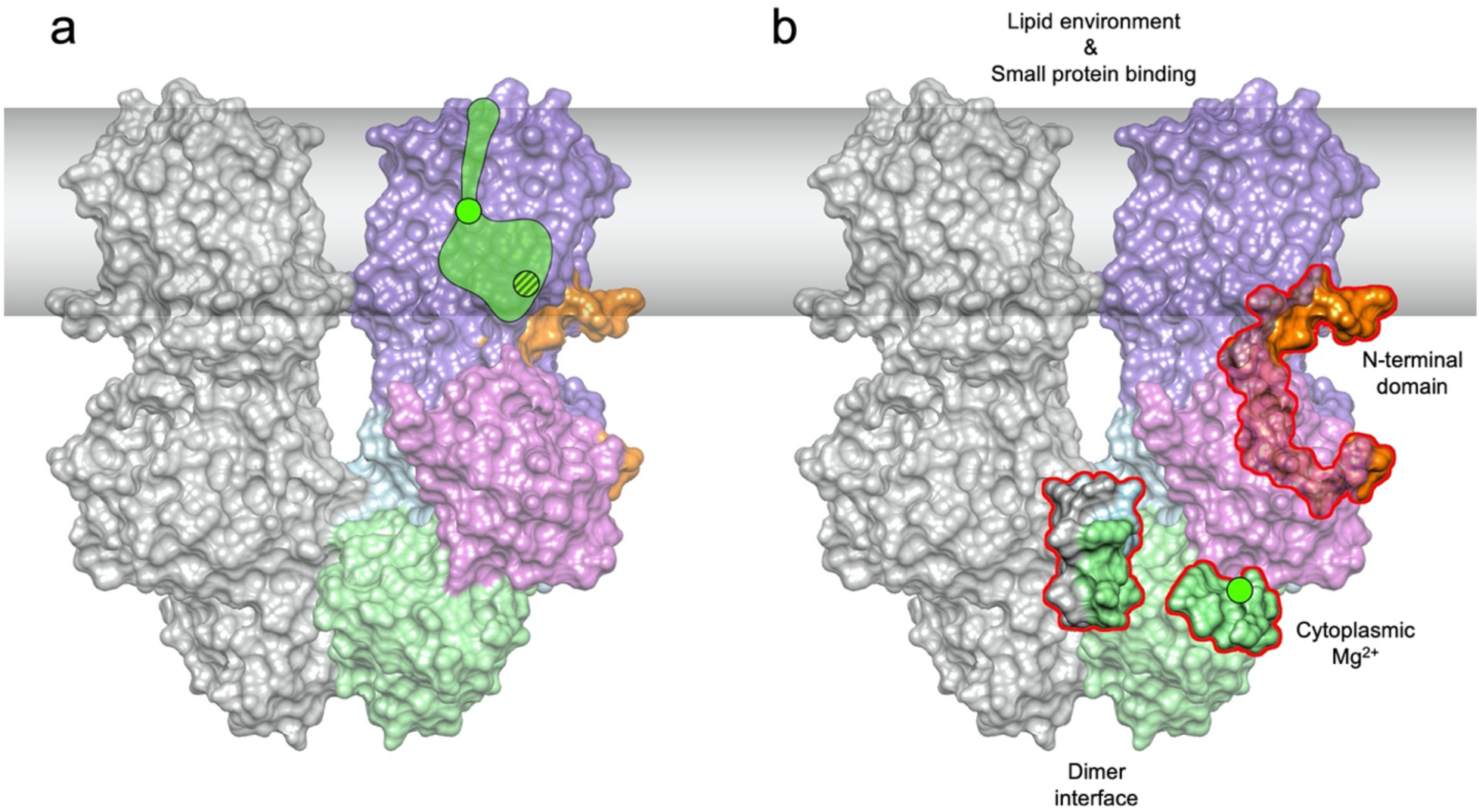
Summary of MgtA structural insights. **a**, Proposed Mg^2+^ ion path based on structural, conservation analysis, and MD simulation data. **b**, Overview of putative regulatory features of MgtA including the dimer interface, cytoplasmic Mg^2+^ binding sites, N-terminal domain, lipid environment, and small protein binding. Green circles represent Mg^2+^ ions. Color scheme is the same as in Fig. 1.

After selection of the Mg^2+^ ion, a constellation of interior residues E331 of TM4, N706 of TM5, and N734 and D738 of TM6 likely maintains the aqueous environment where the acidic/amido residues provide further selectivity for cations by directly coordinating them. We do not detect any additional strong density for other Mg^2+^ ions in this region. However, in the SERCA1a structure, two Ca^2+^ ions were observed in the vicinity of these positions ^41^. Given that mutation of E331 to an alanine in TM4 inactivates the transporter and mutation of D738 modestly impairs activity, we suggest that these sites are involved in Mg^2+^ ion translocation. Cytoplasmic facing residues T702 and N706 of TM5 and Q741 of TM6 may also be involved in the ion transfer towards E331. From residue E331, Mg^2+^ is very close to the cytoplasmic side. We speculate that the large movement of TM1-2 towards the cytoplasmic side as well as the unwinding of TM4 observed in the monomeric structure are involved in the transport mechanism.

### Mechanisms of post-translational regulation of MgtA

As for other P-type ATPases, the levels and activity of MgtA are highly regulated. Our ability to resolve dimeric and monomeric forms of MgtA allows us to make comparisons between the structures and propose models regarding regulation (Fig. 5b). The first feature of interest is the N-terminal tail, which was uniquely resolved in our dimer structure. Interestingly, we observe that the ion binding residue E331 contained within TM4, is rotated outward towards the N-terminal tail in the dimeric structure, while, in the monomeric structure, TM4 unwinds and residue E331 appears to rotate towards the other TM α-helices presumably making it more accessible to ion binding. The N-terminus forms an unsual knot like structure by snorkeling under the A domain making many contacts to the rest of the protein. The A domain appears to be highly flexible in the monomeric form and we were unable to resolve the N-terminus in this structure. These two observations suggest the N-terminus regulates the translocation of ions across the membrane by preventing E331 from interacting with Mg^2+^ and restricts the large scale rearrangements of the A domain observed in the monomeric structure. These suggestions likely are relevant to Na^+^/K^+^-ATPases and H^+^/K^+^-ATPases, which also possess extended N-termini.

We hypothesize that MgtA dimerization provides an additional opportunity for regulation. The interactions between adjacent subunits is formed via many conserved residues residing in the N and P subdomains and could have a number of consequences. Dimerization could allow for functional coupling of the two monomer subunits in a concerted effort to reduce the overall activation energy required for the rate-limiting steps in the highly dynamic catalytic cycle of ion transport. This model is plausible given that the ATP binding pocket of the dimer is accessible to nucleotides and it adopts a similar conformation as other P-type ATPases. Furthermore, enzyme coupling has previously been shown in other family members of the P-type ATPases such as the Na^+^/K^+^-ATPase and the SERCA Ca^2+^-ATPase ^53,61^. It is structurally unclear how coupling might be coordinated. Alternatively, our dimeric structure might only represent one state of the transport cycle or oligomerization modulates the stability of the protein.

We also suggest that the two Mg^2+^ ion densities detected between the A domain and N or P subdomains in our dimeric MgtA structure have regulatory roles. Mutational disruption of one of the two cytoplasmic Mg^2+^ binding sites reduced transport activity. The functional significance of the remaining binding site will require additional probing. The strong reduction in transport activity of MgtA_D441A_ and the proximity of D441 to F445, which forms interactions with ATP, suggests this site is important for enzymatic activity. In the transport cycle of other P-type ATPase transporters such as SERCA, the A domain is very dynamic during transport. The two cytosolic Mg^2+^ binding sites in our structure are formed through contact with the A domain. Thus, we speculate that they may act to stabilize the closed conformation typically induced by ATP binding, possibly allowing for increased transport activity. Alternatively, the cytoplasmic sites could serve as sensors of intracellular levels of Mg^2+^ given that high levels of Mg^2+^ were reported to inhibit MgtA ATP hydrolysis *in vitro* ^34^. It is interesting to note the bacterial magnesium channel CorA has a negative feedback mechanism to sense when intracellular levels of Mg^2+^ are high (Pfoh *et al.*, 2012, Dalmas *et al.*, 2014).

One additional level of MgtA regulation already proposed by other studies is the lipid environment ^34^. Given that P-type ATPases have been shown to be sensitive to detergents used for solubilization ^47^ and that they can respond to lipids such as cardiolipin, these transporters will need to be characterized in their native lipid environments to further elucidate the impacts of oligomerization and lipid composition on the catalytic cycle.

Small proteins of <50 amino acid in length, have been shown to control MgtA stability by regulating FtsH-mediated degradation in both *E. coli* and *S. enterica* ^27,40^. Intriguingly, FtsH substrate recognition and degradation is sensitive to the oligomeric state of some substrates ^62^. Whether MgtA oligomerization is controlled by small proteins is currently unknown. AlphaFold multimer analysis of the various small proteins bound to the transporter suggests that the small proteins share a common binding site on MgtA (Extended Data Fig. 25). Intriguingly, the ability of small proteins to post-translationally regulate P-type ATPases appears to be wide-spread for this family. The SERCA Ca^2+^ transporter is regulated by multiple small proteins that bind to a common interface and are critical for its regulation including Sarcolipin (SLN), Myoregulin (MLN), Phospholamban (PLN) and DWORF (Extended Data Fig. 25) ^63–66^. For the bacterial potassium-importing KdpFABC membrane complex structure, the small protein KdpF has been reported to stabilize the complex ^67–69^. Although the ß-subunit of the Na^+^/K^+^-pump is 303 aa, it possesses a single pass TM segment that interacts with the larger α-subunit ^59,60^. The small protein binding sites determined for these examples differ from the site predicted for MgtA.

The MgtA structures presented here raise several questions for the P-type ATPase field. These questions include whether the N-termini of these proteins inhibit or facilitate transport, how dimerization impacts the transport mechanism and how many P-type ATPases adopt dimeric structures, how ion binding to cytosolic domains affects transport and whether the cytosolic domains of other P-type ATPases are bound by intracellular ions, and finally how small proteins affect the structures and stabilities of P-type ATPases and whether a larger number of these transporters will be found to have associated small proteins. Future biochemical and structural studies of the MgtA transporters, together with other work in the field, will provide further mechanistic understanding of these features. Given the importance of Mg^2+^ acquisition by MgtA/B proteins during bacterial infection our structural insights also will facilitate the rational design of drugs aimed at inhibiting this specific transporter. Similarly, given the disease relevance of human P-type ATPase, the existence of additional regulatory sites could present novel therapeutic opportunities.

## Abbreviations

DDM: n-dodecyl-β-d-maltopyranoside
EM: electron microscopy
EDC: 1-ethyl-3-(3-dimethylaminopropyl)carbodiimide hydrochloride
GDN: glyco-diosgenin
HDX-MS: hydrogen deuterium exchange mass spectrometry
LMNG: lauryl maltose neopentyl glycol
MD: molecular dynamics
ORF: open reading frame
RMSD: root-mean-square deviation
SEC: size exclusion chromatography
SERCA: SarcoEndoplasmic Reticulum Ca^2+^-ATPase
TM: transmembrane

## Acknowledgements

We are grateful to Anirban Banerjee for the use of his lab’s AKTA and comments on the manuscript; Jennifer Petersen and Joshua Zimmerberg for access to the T20 electron microscope; the NICE facility with Allison Zeher, Zabrina Lang, and Rick K. Huang for support on the G1 Krios electron microscope; Elizabeth Viverette, Jonathan Bouvette, and Mario Borgnia for support on the G4 Krios electron microscope; and Stéphane Mahé and Joe Cometa for technical support on the T20 and Krios. We also thank Prof. Dr. Koichi Ito for providing us the Mg^2+^-auxotrophic *E. coli* strain (BW25113 Δ*mgtA*, Δ*corA*, Δ*yhiD* DE3). This research was supported by the Intramural Research Programs of the *Eunice Kennedy Shriver* National Institute of Child Health and Human Development (NICHD) (ZIA HD008955 (AS), ZIA HD008855 (GS), and ZIA HD008998 (DM)) and the National Library of Medicine (LM594244 (LA)) as well as a NIGMS PRAT fellowship (1FI2GM146628-01) and NICHD Early Career Awards to RZ and funding from the German Research Foundation (DFG 3542/1-1 (JDL)). This work utilized the computational resources of the NIH HPC Biowulf cluster (http://hpc.nih.gov). Fig. 1a was created with BioRender (<u>BioRender.com</u>).

## Data availability

The data that support this study are available from the corresponding authors upon request. Cryo-EM maps have been deposited in the Electron Microscopy Data Bank (EMDB) under accession codes EMD-42794 (apo-dimer, C2), EMD-42795 (apo-dimer, C1), EMD-42796 (apo-monomer, C1), EMD-42797 (ATP-dimer, C2), EMD-42798 (ATPγS-dimer, C2), EMD-42799 (ADP-dimer, C2). Raw movies will be uploaded to the Electron Microscopy Public Image Archive (EMPIAR). The atomic coordinates have been deposited in the Protein Data Bank (PDB) under accession codes 8UY7 (apo-dimer, C2), 8UY8 (apo-dimer, C1), 8UY9 (apo-monomer, C1), 8UYA (ATP-dimer, C2), 8UYB (ATP□S-dimer, C2), 8UYC (ADP-dimer, C2); Trajectories and AMBER input files for simulations with and without ATP are publicly available at Zenodo (http://doi.org/10.5281/zenodo.10017395). The mass spectrometry proteomics data have been deposited to the ProteomeXchange Consortium via the PRIDE partner repository ^70^.

Anonymous reviewer access is available upon request. The source data for Fig. 2b, and Extended Data figures 5, 6, and 15d are provided in the Source Data file with this paper.

## Author Contributions

RZ: expressed and purified protein, SDS-PAGE and Western blots, mutagenisis, *in vivo* assays, made negative staining and cryo-EM grids, analyzed data, generated figures, wrote manuscript.

FZ: performed NATIVE-PAGE, made negative staining and cryo-EM grids, screened negative staining and cryo-EM grids, helped collect negative staining and cryo-EM data.

PF: performed mass spectrometry analysis of crosslinked samples, data evaluation and generated figures.

JZ: performed HDX-MS experiments and HDX-MS data evaluation and generated figures.

HL: performed Molecular Dynamics simulations.

LA: carried out computation sequence/structural analysis, identified sequence motifs and generated large sequence alignment.

JDL: analyzed crosslinking and HDX-MS data.

AS: performed Molecular Dynamics simulations, analyzed data, generated figures, wrote manuscript, designed the project.

GS: analyzed data, generated figures, wrote manuscript, initiated and designed the project.

DM: collected and processed negative staining data, processed cryo-EM data, produced structural models, analyzed data, generated figures, wrote manuscript, designed the project.

All authors contributed to the manuscript.

## Methods

### Plasmid construction

All primers, plasmids, details of plasmid construction, and strains used in protein expression, complementation studies, and co-IP studies can be found in supplementary data 2. The pGEX-MgtA, pGEX-empty, pGEX-MgtA D373N, pGEX-MgtA E331A, pGEX-MgtA D738A, pGEX-MgtA D780A, pGEX-MgtA dimer (12mer) plasmids were *de novo* gene synthesized by GenScript. Plasmids created in this study were assembled using the Gibson Assembly Cloning Kit (New England Biolabs). All plasmid sequences were confirmed by whole plasmid sequences services provided my Plasmidsaurus SNPsaurus LLC.

### Protein expression and membrane preparation

*E. coli* C43 (DE3) cells were transformed with the plasmid pETDuet-1-MgtAS, allowed to recover for 1 h at 37°C with shaking at 250 rpm. Following recovery, the cells were grown overnight at 37°C with shaking in LB media containing 100 *µ*g/mL ampicillin (amp). The saturated cultures were diluted 1:100 into LB media with 100 *µ*g/mL amp and incubated at 37°C to OD_600_ ∼ 0.2. The 2 L cultures were cooled to 18°C prior to induction with 0.5 mM Isopropyl-ß-D-1-thiogalactopyranoside (IPTG) and then grown overnight at 18°C shaking at 200 rpm. Cells were collected after 16 h by centrifugation at 5,000 x g for 10 min at 4°C. Cell pellets were resuspended in 50 mM Tris/HCl pH 7.5, 50 mM K_2_SO_4_, 5 mM MgCl_2_, 1 mM DTT (wash buffer) and Roche cOmplete^™^ Protease Inhibitor Cocktail. The cell resuspension was passed 3 times using a microfluidizer processor (Microfluidics) at 20,000 psi. Crude cell lysate was cleared by centrifugation at 15,000 x g for 30 min. The cleared supernatant was centrifuged at 100,000 x g for 1 h. The membrane pellet was rinsed or washed by homogenization in 50 mL with wash buffer to remove residual soluble proteins. Membranes were resuspended using a Potter-Elvehjem-type tissue homogenizer at a final concentration of 10 mg/mL. All cell lysis steps were carried out at 4°C. Glycerol was added to resuspended membranes at a final concentration of 20%. Aliquots were flash frozen in liquid nitrogen and stored at −80°C until further use.

### Membrane solubilization optimization

Membranes (10 mg/mL) were solubilized with 1% glyco-diosgenin (GDN, Anatrace), a synthetic drop-in substitute for Digitonin, 1% n-Dodecyl-β-Maltoside (DDM, Sigma), or 1% Lauryl Maltose Neopentyl Glycol (LMNG, Anatrace) for 16 h rotating at 4°C. Insoluble membranes were pelleted by centrifugation at 100,000 x g for 1 h at 4°C. The supernatant containing detergent solubilized membrane proteins were then normalized for protein concentration, analyzed by Blue Native-PAGE using 4-16% BisTris polyacylamide gels. For Blue Native-PAGE samples were run 1 h at 150 V, followed by an additional 1 h at 250 V on ice. To allow for efficient transfer to PVDF membranes, the cathode buffer was diluted 1/10 (light blue) when the protein had migrated 1/3 of the gel. The gels were then transferred to PVDF membranes for 1 h at 100 V. Following transfer, the blots were blocked with Tris-buffered saline with 0.1% Tween-20 detergent (TBST) + 5% milk and probed using TBST + 5% milk with anti-FLAG (ANTI-FLAG(R) M2-Peroxidase (HRP) by Sigma) or anti-His_6X_ (His HRP conjugate by Qiagen) antibodies at a 1:1,000 dilution. Blots were washed 3 times with TBST for 5 min and imaged.

### Protein purification

Membranes (10 mg/mL) were solubilized with 1% glyco-diosgenin (GDN) for 16 h rotating at 4°C. Insoluble membranes were pelleted by centrifugation at 100,000 x g for 1 h at 4°C. Imidazole was added to the solubilized membrane proteins at a final concentration of 15 mM and the sample was batch bound with Ni-NTA resin for 3 h at 4°C. Resin was washed with 30 column volumes of Buffer A (50 mM Tris/HCl pH 7.5, 50 mM K_2_SO_4_, 5 mM MgCl_2_, 15 mM imidazole, 0.007% GDN, 1 mM DTT). MgtA was eluted from Ni-NTA resin with Buffer B (50 mM Tris/HCl pH 7.5, 50 mM K_2_SO_4_, 5 mM MgCl_2_, 200 mM imidazole, 0.007% GDN, 1 mM DTT). Eluted MgtA was spin concentrated to ∼15 mg/mL prior to size exclusion chromatography (SEC) using 100 kDa Amicon centrifuge tubes. Concentrated MgtA was spun at 15,000 x g for 30 min at 4°C prior to SEC using a Superose 6 Increase 10/300 GL with Buffer C (50 mM Tris/HCl pH 7.5, 50 mM K_2_SO_4_, 5 mM MgCl_2_, 0.007% GDN, 2 mM DTT). Fractions were analyzed by SDS-PAGE and Blue Native-PAGE using 4-16% BisTris polyacrylamide gels. For SDS-PAGE samples were run at 200 V on ice until the dye front reached the bottom of the gel. For Blue Native-PAGE samples were run 1 h at 150 V, followed by an additional 1 h at 250 V on ice using cold buffer. The desired fractions were pooled, and spin concentrated using 100 kDa Amicon centrifuge tubes to ∼2-3 mg/mL prior to cryo grid preparations.

### Crosslinking and mass spectrometry

MgtAS was purified as described above with slight modifications. To remove amines which interfere with crosslinking, HEPES was used in place of Tris. Purified MgtAS at a concentration of 1 mg/mL was crosslinked with 0.5% 1-ethyl-3-(3-dimethylaminopropyl)carbodiimide hydrochloride (EDC) in 50 mM HEPES pH 7.5, 50 mM K_2_SO_4_, 5 mM MgCl_2_, 0.007% GDN, 2 mM DTT at room temperature for 1 h. The protein samples were then quenched with equal volumes of 2x Laemmli loading buffer and heat-denatured at 95°C for 5 min. Cross-linked MgtA was resolved using SDS-PAGE followed by Coomassie Blue G-250 staining. The ∼100 kDa band for MgtA was cut and stored at −20°C.

Gel-bands were digested with the In-Gel Tryptic Digestion Kit (Thermo Fisher Scientific, Dreieich, Germany) with minor adaptations. The excisions were destained, reduced and alkylated according to the manufacturers protocol and digested overnight with trypsin (SERVA, Heidelberg, Germany) followed by 4 h of digestions with chymotrypsin (Sigma-Aldrich, St. Louis, USA) at 37°C. The digested peptides were transferred to a clean tube and solvents were evaporated in a speed vac (Eppendorf). Dried peptides were reconstituted in 5% acetonitrile (ACN) with 0.1% formic acid (FA). Peptides were loaded onto an Acclaim PepMap C_18_ capillary trapping column (particle size 3 µm, L = 20 mm) and separated on a ReproSil C_18_-PepSep analytical column (particle size = 1,9 *µ*m, ID = 75 *µ*m, L = 50 cm, Bruker Coorporation, Billerica, USA) using a nano-HPLC (Dionex U3000 RSLCnano) at a temperature of 55°C. Trapping was carried out for 6 min with a flow rate of 6 μL/min using a loading buffer composed of 0.05% trifluoroacetic acid in H_2_O. Peptides were separated by a gradient of water (buffer A: 100% H_2_O and 0.1% FA) and acetonitrile (buffer B: 80% ACN, 20% H_2_O, and 0.1% FA) with a constant flow rate of 250 nL/min. The gradient went from 4% to 48% buffer B in 45 min. All solvents were LC-MS grade and purchased from Riedel-de Häen/Honeywell (Seelze, Germany). Eluting peptides were analyzed in data-dependent acquisition mode on an Orbitrap Eclipse mass spectrometer (Thermo Fisher Scientific, Dreieich, Germany) coupled to the nano-HPLC by a Nano Flex ESI source. MS^1^ survey scans were acquired over a scan range of 350–1400 mass-to-charge ratio (m/z) in the Orbitrap detector (resolution = 120k, automatic gain control (AGC) = 2e^5^, and maximum injection time: 50 ms). Sequence information was acquired by a “ddMS^2^ OT HCD” MS^2^ method with a fixed cycle time of 5 s for MS/MS scans. MS^2^ scans were generated from the most abundant precursors with a minimum intensity of 5e^3^ and charge states from two to eight. Selected precursors were isolated in the quadrupole using a 1.4 Da window and fragmented using higher-energy C-trap dissociation (HCD) at 30% normalized collision energy. For Orbitrap MS^2^, an AGC of 5e^4^ and a maximum injection time of 54 ms were used (resolution = 30k). Dynamic exclusion was set to 30 s with a mass tolerance of 10 parts per million (ppm). Each sample was measured in duplicate LC-MS/MS runs. MS raw data were processed using the MaxQuant software (v2.1.0.0) with customized parameters for the Andromeda search engine. Spectra were matched to a FASTA file containing the MgtA and MgtS sequences downloaded from UniProtKB (April 2021), a contaminant and decoy database, with a minimum tryptic peptide length of six amino acids and a maximum of five missed cleavage sites. Precursor mass tolerance was set to 4.5 ppm and fragment ion tolerance to 20 ppm, with a static modification (carboxyamidomethylation) for cysteine residues. Acetylation on the protein N-terminus and oxidation of methionine residues were included as variable modifications. A false discovery rate (FDR) below 1% was applied at crosslink, peptide, and modification levels. Identified crosslinked peptides were manually curated and high-score crosslinks were mapped on a monomeric subunit of our dimeric MgtA cryo-EM structure (PDB: 8UY7) using UCSF Chimera ^71^, in which the distances between the side-chain of the residues was calculated. All proteomics data (including acquisition and data analysis parameters) associated with this manuscript will be deposited at the ProteomeXchange Consortium (http://proteomecentral.proteomexchange.org).

### Hydrogen deuterium exchange and mass spectrometry

MgtAS was purified as previously described with slight modifications. For solubilization and purification the protein was initially solubilized from membranes using 1% DDM and the subsequent buffers were supplemented with 0.05% DDM. The final SEC buffer was adjusted to contain 5% glycerol as a cryo protectant and aliquots were flash frozen in liquid nitrogen prior to storage at −80°C. Frozen samples were freshly thawed before every HDX-MS experiment. First, equilibration (E) -, Labelling (L) - and quench (Q) - buffer were prepared with H_2_O or D_2_O for the labelling buffer (E/L – buffer: 50 mM Tris/HCl, 50 mM K_2_SO_4_, 5 mM MgSO_4_, 2 mM DTT, 0.05% DDM, pH/pD 7.5; Q - buffer: 150 mM KPO_i_, 0.05% DDM at pH 2.20).

For ATP binding, L-buffer was split into two, one for binding condition where we added 5 mM ATP (disodium salt, Sigma Product number: A3377) and the other L - buffer was not altered for control condition. Next, the sample was diluted from 17 µM to 15 µM with E-buffer containing no ATP and equilibrated at 0°C. E/L-buffers were equilibrated at 20°C and Q – buffer at 0°C before the experiment. For ATPγS binding we used the same buffers as described in the ATP binding experiment, except in the binding condition, we added 5 mM ATPγS (tetralithium salt, Sigma Product number: 119120-25MG) to the E- and L-buffers. Both samples (control and binding) were diluted with their respective E-buffers (without and with ATPγS) to a final concentration of 15 µM. Samples and buffers were equilibrated as described in the ATP binding experiment.

The HDX mass spectrometry experiments were carried out as described previously in ^72^. In brief, the experiment was carried out using a fully automated HDX-2 system supplied by Waters (Milford, USA). Protein samples were diluted in E-buffer for references measurements or L-buffer to start labelling for 2, 20, 60 and 120 min for the ATPγS experiment. In the ATP binding experiment the protein was labelled for 2 and 120 min. All time points were measured in quadruplicates. Next, the samples were quenched subsequently with ice-cold Q-buffer and immediately injected to the LC for online digestion with a Pepsin/Nepenthensin-2 column and C18 reverse phase peptide separation. Eluting peptides were measured on a Synapt-G2-Si mass spectrometer operated in HDMS^E^ mode including ion mobility separation for 3D peptide identification (RT, m/z, drift time).

Non-deuterated peptides were identified using ProteinLynx Global Server 3.0.3. (PLGS, Waters) for each condition (control and binding). Only peptides with a high confidence score (over 6) identified in at least three out of four replicates were retained for further HDX evaluation. Peak picking of all corresponding peaks was performed with HDExaminer software package by Sierra Analytics. All mass spectra of every peptide, time points and replicates were manually analyzed and curated for correct peak identification. Next, all peptides which displayed statistically significance deuterium uptake differences, based on a Student’s t-distribution with a 95% confidence interval (p ≤ 0.05), in at least three out of four time points were used for further evaluation and visualization. For the two time points labeling experiments only peptides which display statistically significance deuterium uptake difference in all labeling timepoints were used for downstream evaluation. Created deuterium uptake difference maps were plotted on our cryo-EM structures of MgtA (PDB: 8UY7 (apo-dimer, C2) and 8UY9 (apo-monomer)) using UCSF Chimera ^71^.

### Complementation assays

Function of MgtA and mutant alleles were assessed using a strain of *E. coli* lacking Mg^2+^ importers (BW25113 *ΔmgtA ΔcorA ΔyhiD DE*) ^57^. Briefly, plasmids containing wt MgtA or mutants were transformed into the Mg^2+^-auxotrophic strain with selection on plates supplemented with the appropriate antibiotics (50 *µ*g/ml kanamycin or 100 *µ*g/ml ampicillin) and 100 mM MgSO_4_ followed by growth at 37°C overnight. Transformants were grown in LB supplemented with appropriate antibiotics and 100 mM MgSO_4_ at 37°C and 250 rpm shaking overnight. The overnight cultures were normalized to 0.1 OD_600_ and 10-fold serially diluted before plating 3 µl of cells onto LB + agar (1.5% w/v) supplemented with antibiotics and either 100 mM, 1 mM, or no additional MgSO_4_. Cell samples were taken from overnight cultures, normalized by OD_600,_ and immediately resuspended in 1x Laemmli loading buffer, and stored at - 20°C prior to Western blot analysis. For plated growth with induction, IPTG was added at 0.1 mM. Plates were incubated at 37°C overnight prior to being imaged with a ChemiDoc MP Imaging System (Biorad).

### Immunoblot analysis

Frozen cell lysates were boiled at 95°C for 10 min followed by centrifugation at 15,000 rpm for 10 min to clear cellular debris. The resulting supernatant was run on precast 4-15% Tris Glycine gels (Biorad) at 200V for 1h. Proteins were transferred to nitrocellulose membranes at 100V for 1h. Following transfer, the blots were blocked with Tris-buffered saline with 0.1% Tween-20 detergent (TBST) + 5% milk and probed overnight at 4°C using TBST + 5% milk with anti-MgtA (1:2,500; generated in this study). After overnight incubation, blots were washed 3 times with TBST for 5 min and probed with goat anti-rabbit HRP antibody (1:10,000) for 1 h at room temperature. Blots were washed 3 times with TBST for 5 min and visualized using a ChemiDoc MP Imaging System (Biorad).

### *In vivo* dimerization assay

BW25113 *ΔmgtA ΔcorA ΔyhiD DE* cells were transformed with the plasmids pBAD24-MgtA-H6 and pACYC-MgtA-SPA or pBAD-MgtA-H6 and empty pACYC vector, allowed to recover for 1 h at 37°C with shaking at 250 rpm. Following transformation cells were plated onto LB + agar plates containing 100 *µ*g/mL ampicillin and 50 *µ*g/mL kanamycin, and 100 mM MgSO_4_ and grown at 37°C overnight. Large culture overnights were inoculated with single colonies and grown overnight at 37°C with shaking in LB media containing 100 *µ*g/mL ampicillin and 50 *µ*g/mL kanamycin. Cells were collected by centrifugation at 5,000 x g for 10 min at 4°C. Cell pellets were resuspended in 50 mM Tris/HCl pH 7.5, 50 mM K_2_SO_4_, 5 mM MgCl_2_ (wash buffer) and Roche cOmplete^™^ Protease Inhibitor Cocktail. Membranes were isolated as previously described for MgtAS purification and resuspended at a final concentration of 10 mg/mL.

Membranes (10 mg/mL) were solubilized with 1% glyco-diosgenin (GDN) for 16 h rotating at 4°C. Insoluble membranes were pelleted by centrifugation at 100,000 x g for 1 h at 4°C. The sample was batch bound with FLAG resin for 2 h at 4°C. Resin was washed with 30 column volumes of Buffer A (50 mM Tris/HCl pH 7.5, 50 mM K_2_SO_4_, 5 mM MgCl_2_, 0.007% GDN). MgtA was eluted from FLAG resin with 0.2 M glycine pH 2.4 and neutralized in 1x Laemmli loading buffer. Elutions were analyzed by Western blot using polyclonal anti-MgtA as described for complementation assays.

### Negative staining electron microscopy

3 µL diluted MgtA (0.01 mg/mL) was deposited on a carbon-coated 400 mesh copper grid (CF400-CU, EMS, Hatfield, PA, USA) following glow discharging (PELCO easiGlow Glow Discharge Cleaning System, Ted Pella, CA, USA) for 1 min at 15 mA. The liquid drop was allowed to incubate on the grid for 1 min. The liquid was then wicked away with blotting paper (Whatman), and quickly washed with 3 µL of Nano-W Negative Stain (2% methylamine tungstate, Nanoprobes, Yaphank, NY, USA) followed by immediate incubation with 3 µL of Nano-W for 1 additional min. Filter paper was used to wick away excess stain, followed by gently waving the grid for 1 min to produce different thicknesses or waves of stained areas. Transmission Electron Microscopy (TEM) was performed using an FEI Tecnai T20 TEM (Thermo Fisher, Waltham, MA, USA) operated at a voltage of 200 kV with a direct electron detector K2 Summit (Gatan Inc., Pleasanton, CA, USA) using SerialEM ^73^. Data was collected at a nominal magnification of 25,000 x, a binned pixel size of 3.04 Å/px and a defocus ranging from ∼1.3 to ∼2.4 µm. At a dose rate of 5 e^−^/px/s, 10 s exposures with 0.2 s frames were collected. Each 50-frame-movie was motion corrected using SerialEM ^73^, and a total of 343 images were collected. Data was unpacked using Bsoft ^74^ and processed in CisTEM ^75^. Aligned images were imported, CTF was estimated, and particles picked using a maximum radius of 90 Å and a characteristic radius of 30 Å and a threshold of 5. 76,993 particles were extracted using a box size of 84 px (∼255 Å) and multiple rounds of 2D classification were performed.

### Cryo-EM grid preparation

For apo-MgtA grids, Quantifoil R1.2/1.3 400 mesh copper grids (EMS) were glow discharged in a PELCO easiGlow Glow Discharge Cleaning System (Ted Pella) at 15 mA for 1 min. MgtAS sample (3 µL) at a concentration of 3.2 mg/mL (Extended Data Fig. 6a) was applied to the glow discharged grids. After blotting for 4-5 s in a Leica EM GP2 plunge freezer (Leica Microsystems, Wetzlar, Germany) with the chamber set to 5°C and 95% humidity, the grids were immediately plunge frozen into liquid ethane and stored in liquid nitrogen. Grids were initially screened on an FEI Tecnai T20 transmission electron microscope operated at a voltage of 200 kV with a direct electron detector K2 Summit (Gatan Inc.) using SerialEM ^73^.

For nucleotide-MgtA grids, Quantifoil R1.2/1.3 400 mesh copper grids were prepared as described above. Prior to grid preparations, MgtAS consisting of a predominantly dimer species was isolated using SEC and spin concentrated to a final concentration of 2.3 mg/mL (Extended Data Fig. 6b). After spin concentration, 10 µl of MgtAS was incubated with 0.5 µL of 100 mM ATP (Sigma, Catalog # A2383) (for 2 min), ATPγS (Sigma, Catalog # 119120) (for 15 min), or ADP (Sigma, Catalog # A2754) (for 15 min) and 3 µL were applied to the glow discharged grids. After blotting for 6 s using a Leica EM GP2 plunge freezer with the chamber kept at 4°C and 95% humidity, the grids were immediately plunge frozen into liquid ethane and stored in liquid nitrogen.

### Cryo-EM data collection

After the grid making conditions were optimized, high resolution movies were collected on a G1 or G4 Titan Krios electron microscope (Thermo Fisher) operated at 300 kV with a K3 direct electron detector in CDS mode and energy filter using a 20 eV slit (Gatan Inc.). The image acquisition was operated with SerialEM ^73^ at a nominal magnification of 105,000 x, a physical pixel size of 0.83 (G1 Krios) or 0.85 (G4 Krios) Å/pixel in super-resolution mode (0.415 or 0.425 Å/px) and a defocus ranging from approximately −0.7 to −2.0 µm. A first dataset was collected with a dose rate of 9.5 Å/px/s (∼7.5 on the camera through the sample), exposure time 0.0712 s/frame (∼1 e^−^/Å^2^) and total exposure time of 4.31 s per movie (∼60 e^−^/Å^2^), resulting in a total of 10,246 movies with 60 frames each. Additional six datasets were collected in a similar manner and the data collection details can be found in Extended Data Table 3.

### Cryo-EM data processing

The overall workflow of image processing of *E. coli* MgtA is illustrated in Extended Data Fig. 8 and additional details can be found in Extended Data Tables 4 and 5 as well as Extended Data figures 9 and 17. All processing of MgtA without the addition of any nucleotides (dataset 1 and 2) was performed within cryoSPARC ^76^. Datasets 3-7 of MgtA in the presence of different nucleotides were processed using cisTEM ^75^ and cryoSPARC ^76^.

For dataset 1 and 2, movies were processed with full-frame motion correction and patch CTF estimation. CryoSPARC’s blob picker with a minimum and maximum particle diameter of 110 and 130 Å was used for automatic particle picking. Particles were extracted using a box size of 768 px (∼319 Å), fourier cropped to 384 px. For dataset 1 and 2, 1,578,214 and 1,775,726 particles were picked from 10,164 and 5,778 selected aligned micrographs. Particles were then subjected to 2D classification to remove junk particles. 1,068,828 and 725,244 particles selected from one round of 2D classification was subjected to a second round of 2D classification. At this point particles from 2D class averages showing clear side views for dimer and monomer species were separated while particles from less clear views including top and tilted views were added to both subsets. Ab-initio reconstruction followed by non-uniform refinement led to initial dimer and monomer reconstructions that were used for heterogenous refinement using all particles to include possible disgarded particles during earlier classifications. Additional ab-initio reconstructions followed by non-uniform refinement was used to cluster the best aligning particles and to improve reolution further, local motion correction and CTF refinement to correct for beam-tilt, spherical aberrations, and per-particle defocus parameters was applied before merging dimer or monomer particles from the two different datasets. Final rounds of 2D classification, ab-initio reconstruction, and non-uniform refinements were used to produce the final reconstructions which were used to estimate and filter to local resolution. All refinements of the dimer were run with (C2) and without (C1) symmetry, while all refinements of the monomer were performed without symmetry. The final dimer map was reconstructed from 160,139 particles to an average resolution of 2.93 Å (C2) or 3.03 Å (C1), while the final monomer map was reconstructed from 78,231 particles to an average resolution of 3.65 Å according to the gold-standard FSC = 0.143 criterion.

For dataset 3 to 7, movies were imported and binned to a pixel size of 0.83 (G1 Krios data, datesets 3-6) or 0.85 (G4 Krios, dataset 7), processed with full-frame motion correction and patch CTF estimation using CisTEM ^75^. After image selection, particles were picked using a maximum radius of 70 Å and a characteristic radius of 50 Å and a threshold of 3. Particles were extracted using a box size of 384 px (∼319 or ∼326 Å) and one round of 2D classification was performed before only getting rid of particles in junk 2D class averages and exporting the selected particle stacks for further processing in cryoSPARC ^76^. Additional rounds of 2D classification, ab-initio reconstructions, non-uniform and heterogenous refinements were performed before ending up with final maps at an average resolution of 3.72 Å, 3.87 Å and 3.75 Å for the MgtA dimer in the presence of 5 mM ATP, ATPγS, and ADP, respectively. The final unsharpened, sharpened and to local resolution filtered cryo-EM maps have been deposited in the Electron Microscopy Data Bank (EMDB) with the following accession codes: EMD-42794 (apo-dimer, C2), EMD-42795 (apo-dimer, C1), EMD-42796 (apo-monomer, C1), EMD-42797 (ATP-dimer, C2), EMD-42798 (ATPγS-dimer, C2), EMD-42799 (ADP-dimer, C2). Raw movies will be uploaded to the Electron Microscopy Public Image Archive (EMPIAR).

### Model building, refinement and validation

A preliminary structural model was generated using both an AlphaFold prediction (AF-P0ABB8) and the 1.6 Å crystal structure of the MgtA nucleotide binding subdomain spanning residues 382-545 of the total of 898 residues (PDB 3GWI, ^38^). Using UCSF Chimera ^71^, the AlphaFold model was separated into overlapping fragments (fragment 1, containing residues 1-86 and 149-281; fragment 2, containing residues 366-548; fragment 3, containing residues 540-700; fragment 4, containing residues 375-548; fragment 5, containing residues 86-144 for TM1-2; and fragment 6, containing residues 270-347 and 697-898 for TM3-4 and TM5-10) and rigid body docked into our 3D dimer map. Following docking, regions which did not fit including the N-terminal domain were removed and built manually followed by flexible fitting in Coot ^77^. In regions of the structure that were less clearly resolved the local resolution filtered as well as the non-sharpened map were used to trace the backbone and to connect segments. Further refinement of the structures were performed using real-space refinement in PHENIX (version 1.20.1-4487) ^78^. The geometry of the structural model was additionally validated using MolProbity ^79^. UCSF Chimera and ChimeraX ^80^ were used for visualization of the structures and to generate all figures in the manuscript. Movies, not including MD simulations, were created using UCSF Chimera. The statistics for the model refinements are in Extended Data Table 4. The fitted pdbs have been made available together with the EM maps with the following PDB IDs: 8UY7 (apo-dimer, C2), 8UY8 (apo-dimer, C1), 8UY9 (apo-monomer, C1), 8UYA (ATP-dimer, C2), 8UYB (ATPγS-dimer, C2), 8UYC (ADP-dimer, C2).

### MD simulations

Simulations were built with the CHARMM-GUI web interface ^81^ using an early cryo-EM resolved dimer structure of MgtA with only the transmembrane magnesium cation included. Symmetric bilayers were constructed with PVPG (20%), PPPE (75%), PVCL2 (5%), mimicking the headgroup chemical composition of the *E. coli* inner membrane, if not the potential (dynamic) asymmetry ^82^. The CHARMM C36m forcefield was used for proteins ^83^ and lipids ^84^. Following initial equilibration and minimization with NAMD ^85^, production simulations were run with the Amber software package (Amber20).Temperature was maintained at 310 K using a Langevin thermostat. Constant atmospheric pressure and zero surface tension was maintained using a Monte Carlo semi-isotropic barostat. Standard CHARMM forcefield parameters were applied (Particle Mesh Ewald for electrostatics, a 12 Å cutoff for non-bonded interactions with force switching between 10 and 12 Å). Simulations of the early cryo-EM resolved MgtA dimer were run for 2 total microseconds, while simulations of the ATP-bound configuration were run for 1.8 microseconds. Trajectories and AMBER input files for simulations with and without ATP are publicly available at Zenodo (http://doi.org/10.5281/zenodo.10017395).

Lipid solvation shell composition is resolved by a Voronoi decomposition as in ^86^. Five representative protein-bound ATP configurations were found via K-medoids ^87^ clustering on ∼1,800 configurations. Lipid shell analysis and ATP clustering were performed with public-domain software developed in-house (http://github.com/alexsodt/shells and https://github.com/alexsodt/clustering, respectively).

Static molecular images from simulations were created using UCSF Chimera ^71^. Simulation movies were created with Tachyon (written by John Stone) and assembled with ffmpeg.

### Sequence analysis

MgtA (National Center for Biotechnology Information (NCBI) Genbank database accession: NP_418663.1) was used as starting query for sequence similarity searches using PSI-BLAST program against the non-redundant (nr) clustered down to 50% sequence identity using the MMseqs program with a profile-inclusion threshold was set at an e-value of 10^−10^. Multiple sequence alignments (MSAs) were constructed using the MAFFT programs. Sequence logos were constructed using these alignments with the ggseqlogo library for the R language. Signal peptides and TM regions were predicted using hidden Markov model (HMM) specifying different sequence regions of a signal peptide and TM proteins in series of interconnected states as implemented in the Phobisu program.

### Structure analysis

The JPred program was used to predict secondary structures using MSAs (see above). Structural models were generated using the RoseTTAfold and AlphaFold2 programs. Multiple alignments of related sequences (>30% similarity) were used to initiate HHpred searches for the step of identifying templates to be used by the neural networks deployed by these programs.

### Comparative genomics and phylogenetic analysis

Clustering of proteins was done using the MMseqs program adjusting the length of aligned regions and bit-score density threshold empirically. Divergent sequences or small clusters were merged with larger clusters if other lines of evidence, including shared sequence motifs, reciprocal BLAST search results, and/or shared genome context associations, supported their inclusion. Phylogenetic analysis was performed using the maximum-likelihood method with the WAG or JTT models (determined empirically from data) with the IQTree. The FigTree program (http://tree.bio.ed.ac.uk/software/figtree/) was used to render phylogenetic trees.

## Supplementary Information

### Extended Data Figures

**Extended Data Fig 1.**
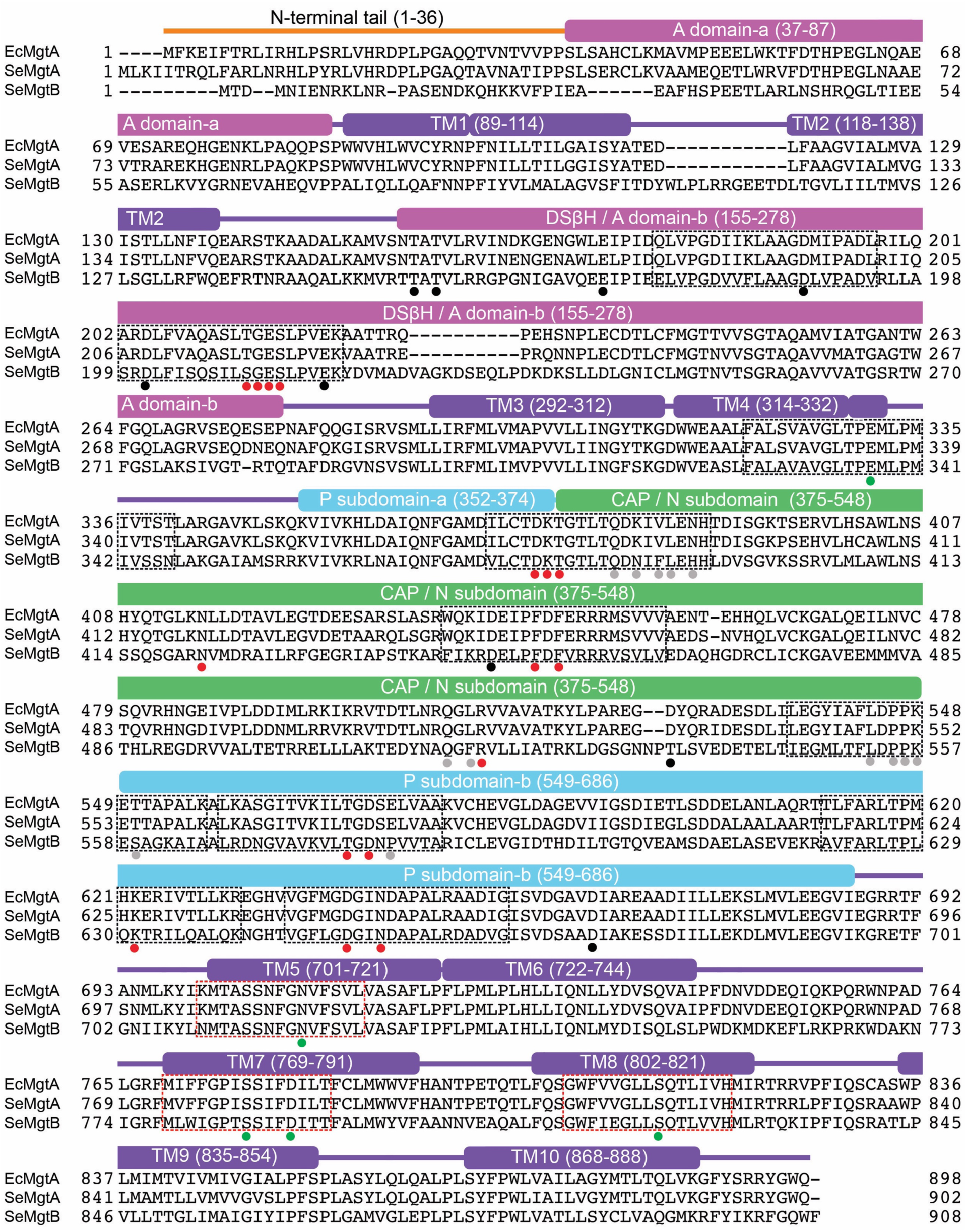
Multisequence alignment of MgtA and MgtB illustrates conserved structural features. Sequence alignment of *E. coli* MgtA (EcMgtA), *S. enterica* serovar Typhimurium MgtA (SeMgtA), and *S. enterica* serovar Typhimurium MgtB (SeMgtB) generated using Clustal Omega. Domains are colored and named according to Fig. 1. Both the canonical P-type ATPase nomenclature and more detailed description of the fold of the domain with boundaries are given. The soluble A domain is split into two regions a and b. The b segment of the A domain is comprised of a <u>D</u>ouble <u>S</u>tranded <u>beta</u>-<u>H</u>elix fold (DSβH). The soluble P subdomain is a noncontiguous segment comprised of two regions a and b that house the key catalytic residues required for phosphorylation. The N or CAP subdomain is a contiguous sequence that binds the nucleotide and aids in catalysis. The P and N subdomains comprise the <u>H</u>alo<u>a</u>cid <u>d</u>ehalogenase (HAD) domain. The TM-spanning alpha-helical regions, as determined by a residue’s alpha-carbon position residing, on average, within the hydrocarbon bilayer interior of the molecular dynamics simulations (+/− 15 Angstroms from the bilayer midplane) are denoted by TM1-10. Gray circles denote residues present at the dimer interface, red circles denote residues near ATP, green circles denote residues coordinating Mg^2+^ in the transmembrane domain, and black circles denote residues coordinating Mg^2+^ in the cytoplasmic domain in our dimeric cryo-EM structures, as annotated in Fig. 2, 3 and 4. Black dashed boxes indicate sequences conserved across all P-type ATPases as shown in the logos in Extended Data Fig. 3. Red dashed boxes indicate sequences specific to Mg^2+^ importers as shown in the logos in Extended Data Fig. 22.

**Extended Data Fig 2.**
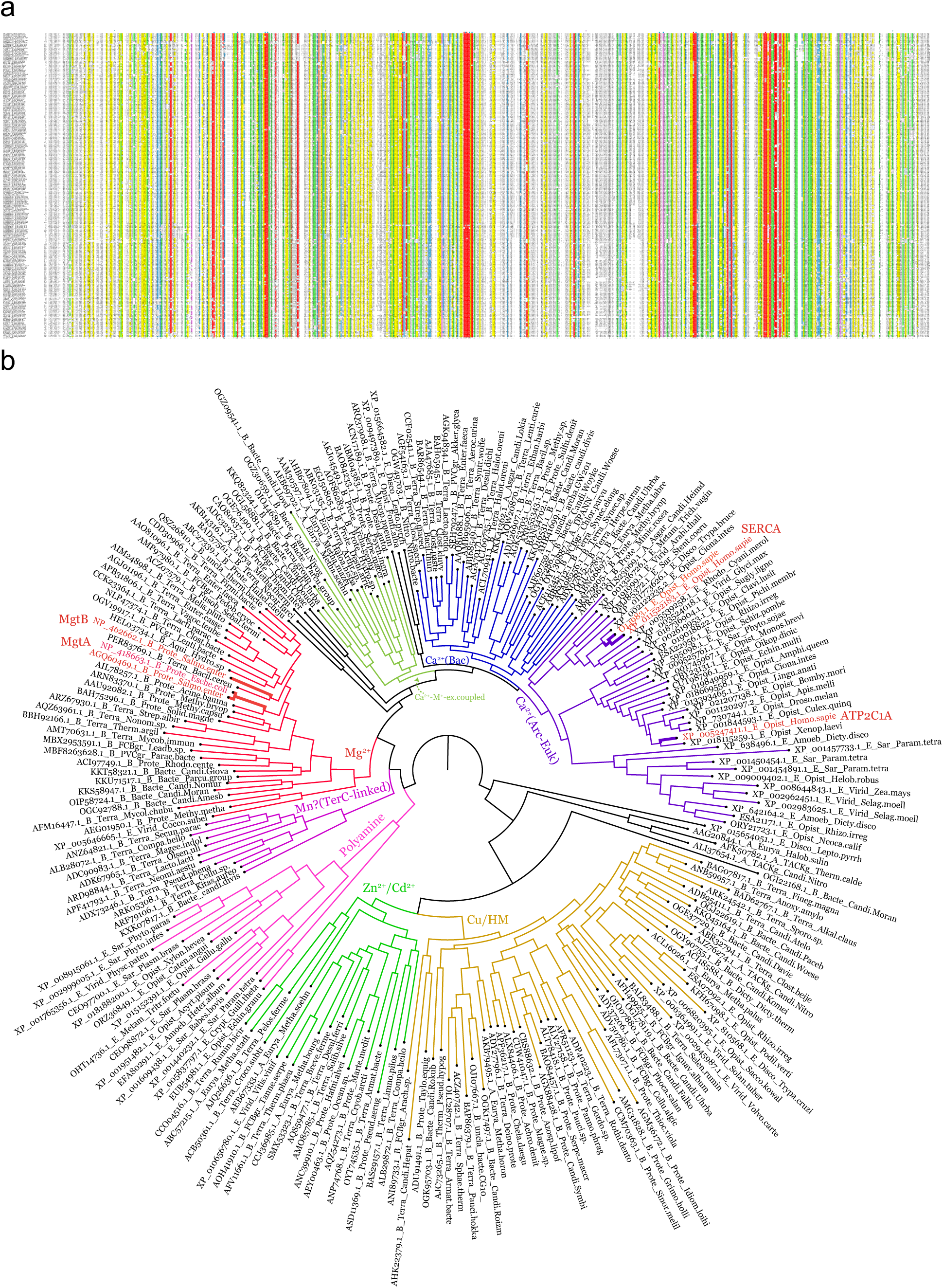
Multisequence alignment of all P-type ATPases. **a,** A multiple sequence alignment of representatives of the eight major clades of P-type ATPases that are shown in different colors in the tree in **b** (see Supplementary Data 1 for an extended sequence alignment). The first six sequences are *E. coli* MgtA, *S. enterica* MgtA, *S. enterica* MgtB; *H. sapiens* ATP2C1 mutated in Hailey-Hailey disease (the HHD mutations are marked with H above the alignment), *H. sapiens* SERCA1 mutated in Brody’s myopathy (the Brody’s myopathy mutations are marked with B above the alignment) and *H. sapiens* SERCA3. The alignment is colored according to the 85% consensus shown in the final line and the sidechain type of the consensus position: l, aliphatic; a, aromatic; h, hydrophobic; +, positive; −, negative; c, charged; p, polar; t, tiny; s, small; b, big. The sequences are labeled by their NCBI Genbank accession, followed by taxonomic marker where B is bacteria, A is Archaea, and E is Eukaryota. This is followed by abbreviations of higher-order taxonomic lineage and species. The positions of the catalytic residues are indicated by red dots, while other conserved positions are shown with blue dots. **b,** In the tree, the clades are labeled as per their known or predicted transport substrates. All the distinctly colored clades are supported with IQtree bootstrap sport of 90% or higher.

**Extended Data Fig 3.**
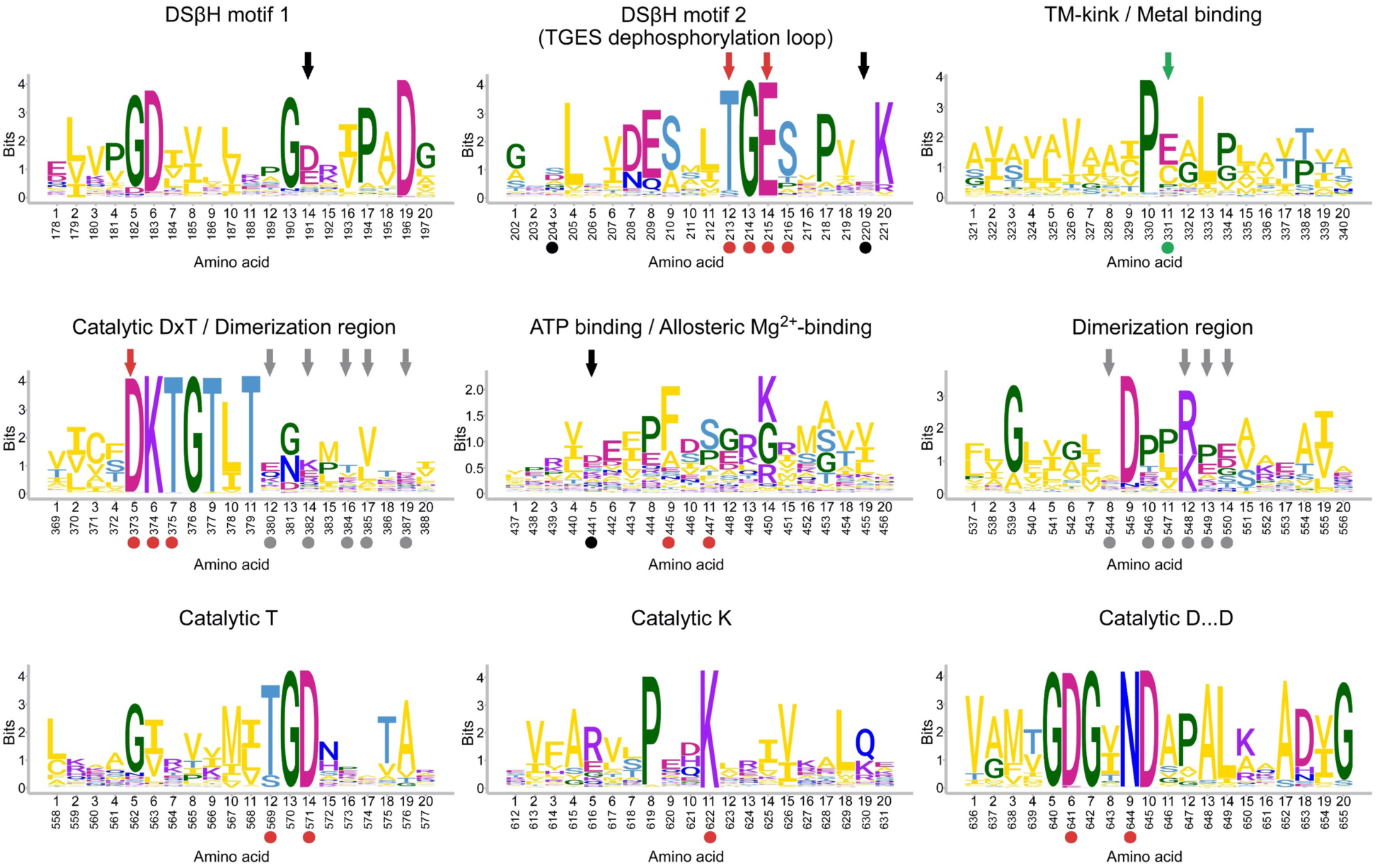
Amino acids related to catalysis and structural architecture are highly conserved across the P-type ATPase family. Sequence logos showing conservation of amino acid residues involved in ATP hydrolysis and structural architecture conserved among entire family of P-type ATPases (indicated by black dashed boxes in Extended Data Fig. 1). Letters represent amino acid abbreviations and height represents the probability of conservation in the P-type ATPase family. As in Extended Data Fig. 1, gray circles denote residues present at the dimer interface, red circles denote residues involved in ATP hydrolysis, green circles denote residues coordinating Mg^2+^ in the transmembrane domain, and black circles denote residues coordinating Mg^2+^ in the cytoplasmic domain in our dimeric structures, as annotated in Fig. 2, 3 and 4. Gray, red, green and black arrows, respectively, indicate residues located at the dimer interface, involved in ATP hydrolysis, coordinating the transmembrane or cytoplasmic Mg^2+^, which were mutated in subsequent experiments.

**Extended Data Fig 4.**
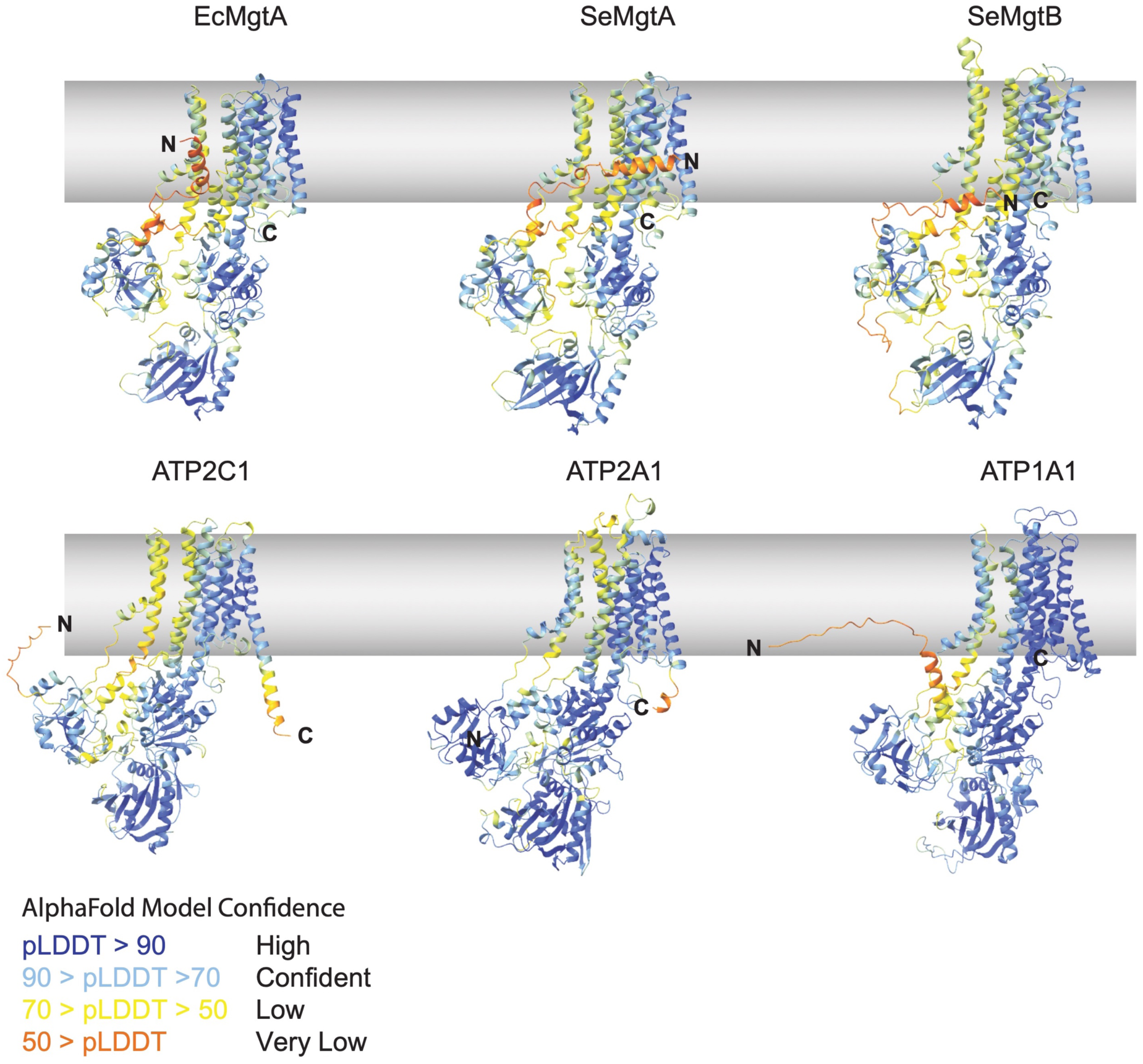
EcMgtA is predicted to be structurally similar to SeMgtA/B and the closest mammalian homologs ATP2C1 and SERCA. AlphaFold2 model for MgtA and related transporters predict a similar overall structural architecture. Models are displayed based on AlphaFold2 predictions for *E. coli* MgtA (EcMgtA), *S. enterica* MgtA (SeMgtA) and *S. enterica* MgtB (SeMgtB). The highest confidence *Homo sapiens* ATP2C1 (a Ca^2+^/Mn^2+^ transporter) and SERCA (ATP2A1) models are shown along with the highest confidence model for *H. sapiens* Na^+^/K^+^ transporter (ATP1A1), which also has an N-terminal tail. The per-residue confidence metric (pLDDT) was used to color the models. Regions expected to be modelled with higher confidence are blue while regions with lower confidence are orange as indicated in the legend. The structures of all N-termini are of low confidence.

**Extended Data Fig 5.**
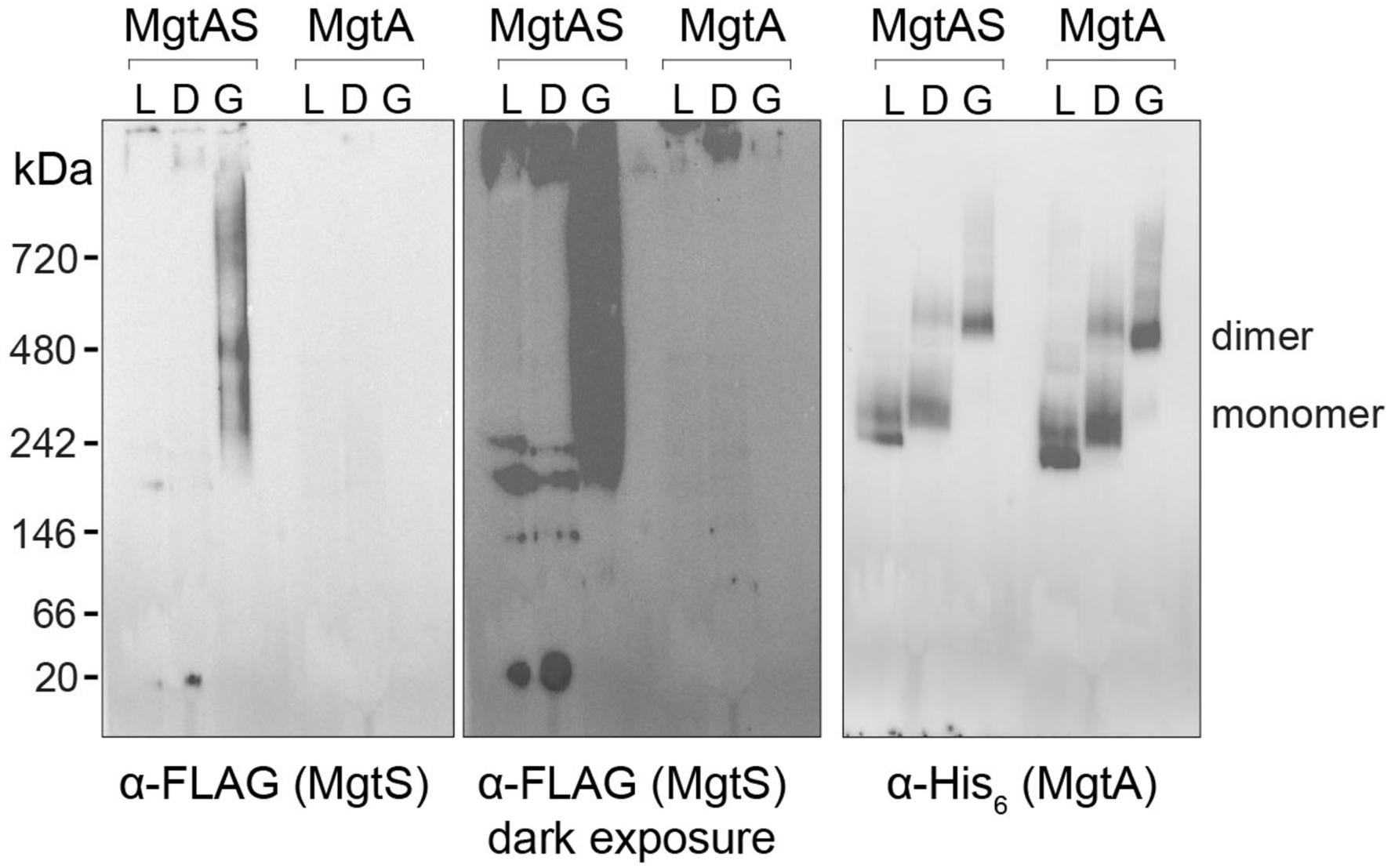
Weaker detergents preserve MgtA native protein-protein interactions. Solubilization of membranes expressing MgtA or MgtAS with detergents of varying strengths show differences in native protein interactions when analyzed by G250 Blue-Native gel and Western blot analysis. Membranes from cells overexpressing MgtA or MgtAS were solubilized with the detergents LMNG (L), DDM (D), or GDN (G) prior to Blue-Native PAGE and Western blot analysis using α-FLAG antibodies against tagged MgtS (left two panels; middle panel is a longer exposure of the left panel) or α-His_6_ antibodies against tagged MgtA (right panel).

**Extended Data Fig 6.**
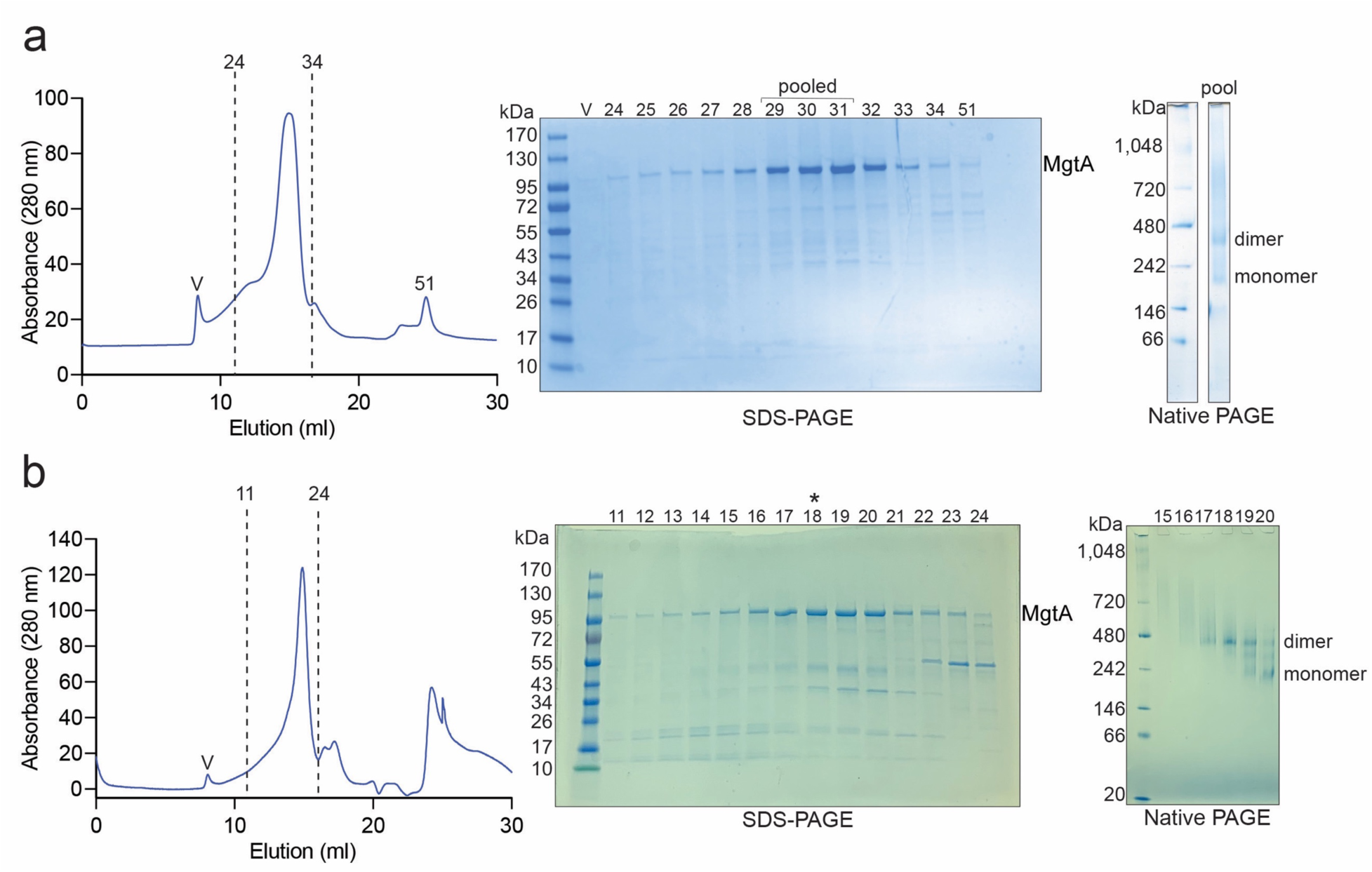
MgtA forms two distinguishable MW species when purified from *E. coli.* **a**, Purification of MgtA results in two distinct MW protein complexes. SEC profile (left) of MgtA used to solve the dimer and monomer structures of MgtA in Fig. 1. Fractions were analyzed by SDS-PAGE (middle) and Blue-Native PAGE gels (right). V indicates the void volume of the SEC column. Fractions that were used to solve the structure are referred to as pooled. **b**, Dimeric MgtA can be separated from monomeric MgtA. SEC profile (left) of MgtA used to solve the nucleotide bound dimer structures of MgtA in Fig. 3 and Extended Data Fig. 16. Fractions were analyzed by SDS-PAGE (middle) and Blue-Native PAGE gels (right). Fraction 18 which possessed predominantly dimer species was used for structural analysis.

**Extended Data Fig 7.**
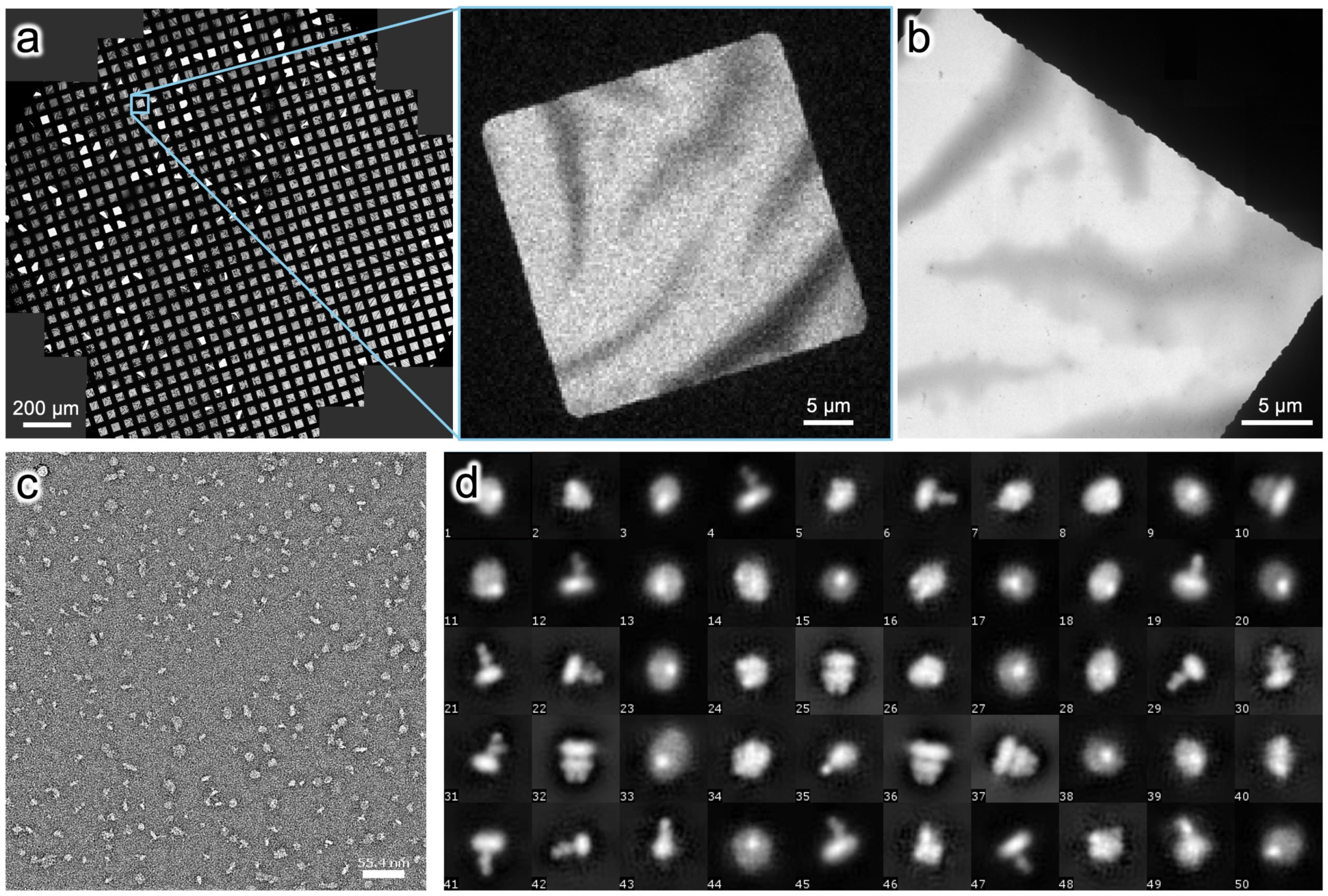
Negative staining EM analysis of purified MgtA. **a**, Low magnification atlas of a negative staining grid of MgtA diluted to 0.01 mg/ml of purified MgtA and zoom in into a suitable grid square for data collection with a decent thickness of staining in the lighter regions. **b**, Medium magnification image of part of the grid square highlighted in a. **c**, Representative micrograph of negatively stained single particles shows no protein aggregation and sufficient particle density. **d**, 2D class averages with a box size of 84 px (∼255 Å) indicating larger particles as well as monomeric looking particles in different orientations.

**Extended Data Fig 8.**
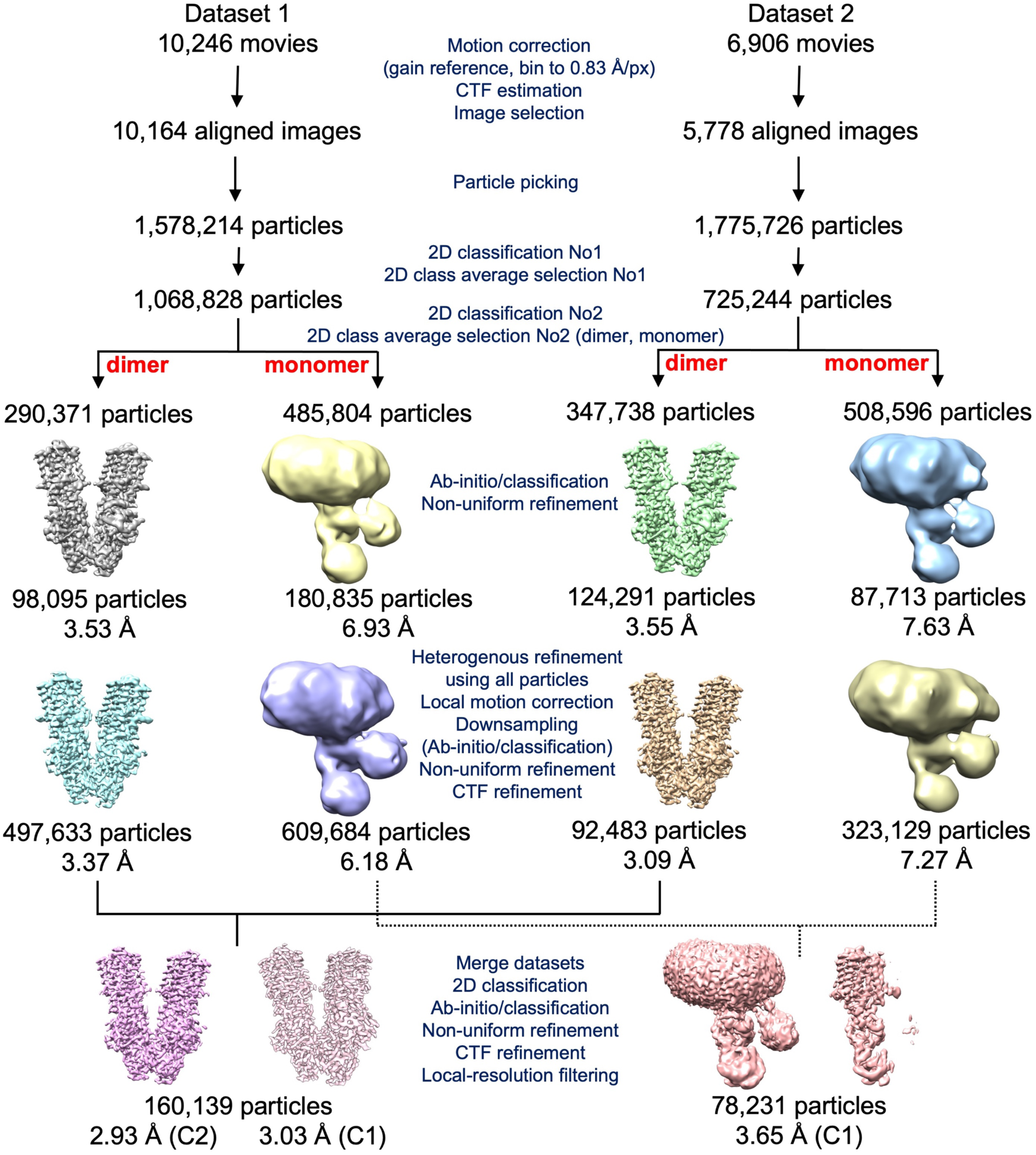
Schematic showing cryo-EM data processing workflow for dimeric and monomeric MgtA. Two datasets for EcMgtA in the presence of 5 mM MgCl_2_ were initially processed separately before merging dimeric and monomeric particles from each dataset followed by additional classification and refinement steps to obtain the final reconstructions used for model building. Number of movies and particles as well as the by cryoSPARC estimated resolution is indicated (also see Extended Data Fig. 9 and Extended Data Tables 3 and 4).

**Extended Data Fig 9.**
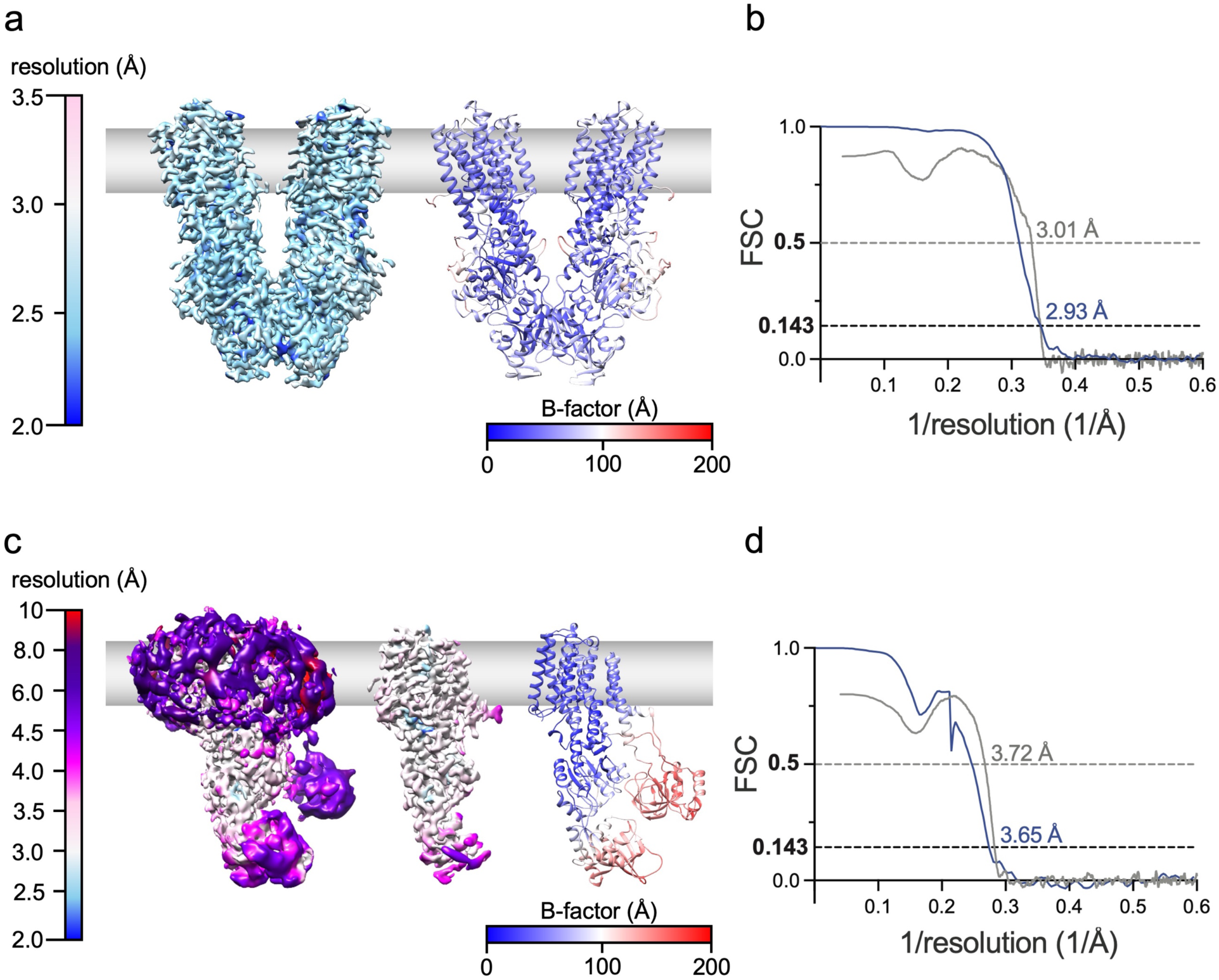
Local and average resolution estimation of the dimeric and monomeric MgtA cryo-EM maps and B-factor distribution of models. **a**, Final dimer reconstruction filtered and colored to local resolution (left) and fitted model colored according to B-factor distribution (right) indicating rigid and more flexible regions of the complex. **b**, Fourier Shell Correlation (FSC) curve of the final dimer reconstruction of MgtA in blue indicating an average resolution of 2.93 Å according to the FSC=0.143 criterion. FSC between the final dimer map and fitted model is shown in gray. **c**, Final monomer reconstruction filtered and colored to local resolution at different thresholds (left and middle) and fitted model colored according to B-factor distribution (right) indicating rigid and more flexible regions. **d**, FSC curve of the final monomer reconstruction of MgtA in blue indicating an average resolution of 3.65 Å according to the FSC=0.143 criterion. FSC between the final monomer map and fitted model is shown in gray.

**Extended Data Fig 10.**
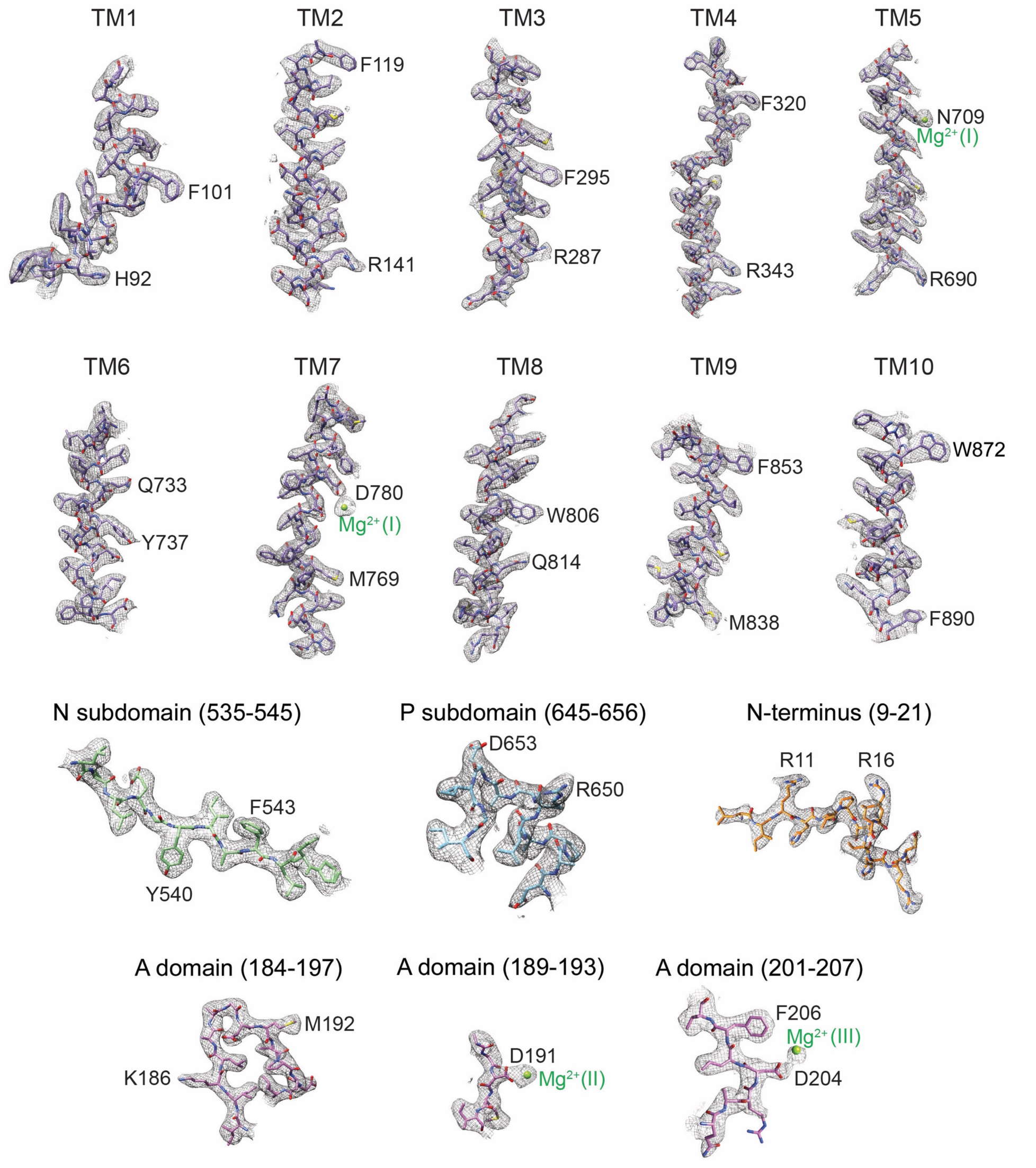
Example regions documenting quality of cryo-EM dimer map for key structural features. Cryo-EM density and atomic model of the TM segments (1-10), soluble domains (A, P, N), N-terminus, and Mg^2+^ ions colored as in Fig. 1. Atomic model of each structural element is shown in stick representation, the atoms are colored by heteroatom within the cryo-EM density and the corresponding density is represented in gray mesh.

**Extended Data Fig 11.**
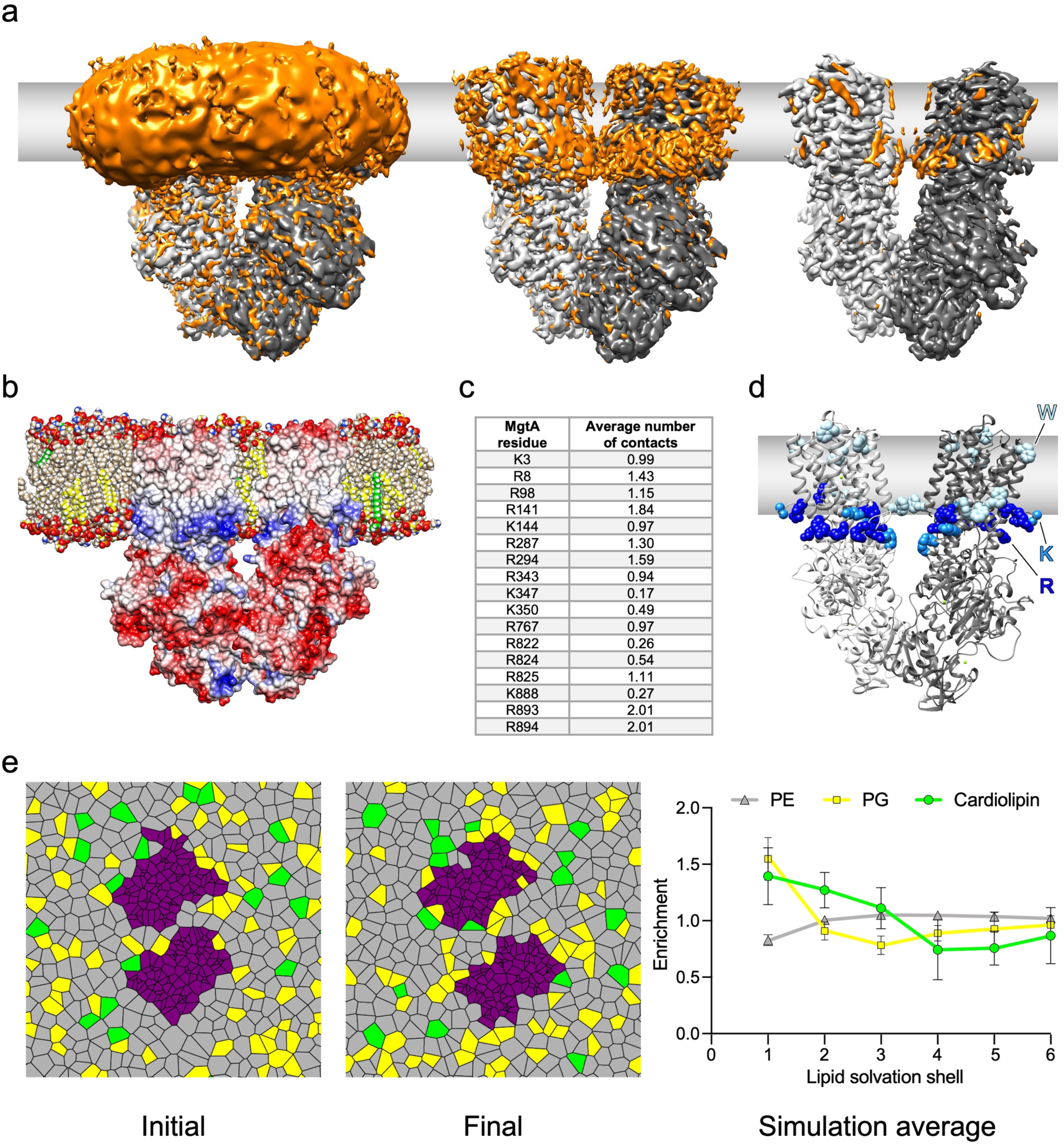
Experimental support of transmembrane borders and lipid distribution. **a**, Side views of cryo-EM maps at different thresholds showing extra densities in orange corresponding to the detergent micelle, detergent molecules, or potential co-purified lipids near the transmembrane region. **b**, Side view of MgtA simulated in a native lipid environment displayed in surface representation colored by electrostatic potential (UCSF Chimera coloring varies from red [−10 kcal/mol/e] to blue [+10 kcal/mol/e] with distance-dependent dielectric constant 4, distance from surface 1.4). Phospholipids corresponding to phosphatidylethanolamine (PE), phosphatidylglycerol (PG) and cardiolipin are colored tan, yellow, and green, respectively. **c**, Arginine and lysine residues that interact with anionic lipids during at least 20% of the simulation, with cutoff enclosing the first peak of the radial distribution function. **d**, Side view of MgtA with arginine (R), lysine (K), and tryptophan (W) residues near the lipid membrane borders highlighted in blue spheres. **e**, Voronoi decomposition of lipid centers-of-geometry in the cytoplasmic leaflet initially (initial) and at the end of the simulation (final). Yellow and green are anionic PG and cardiolipin lipids, respectively. At right, average enrichment or depletion of lipids based on solvation shell as assigned by Voronoi decomposition averaged over the trajectory.

**Extended Data Fig 12.**
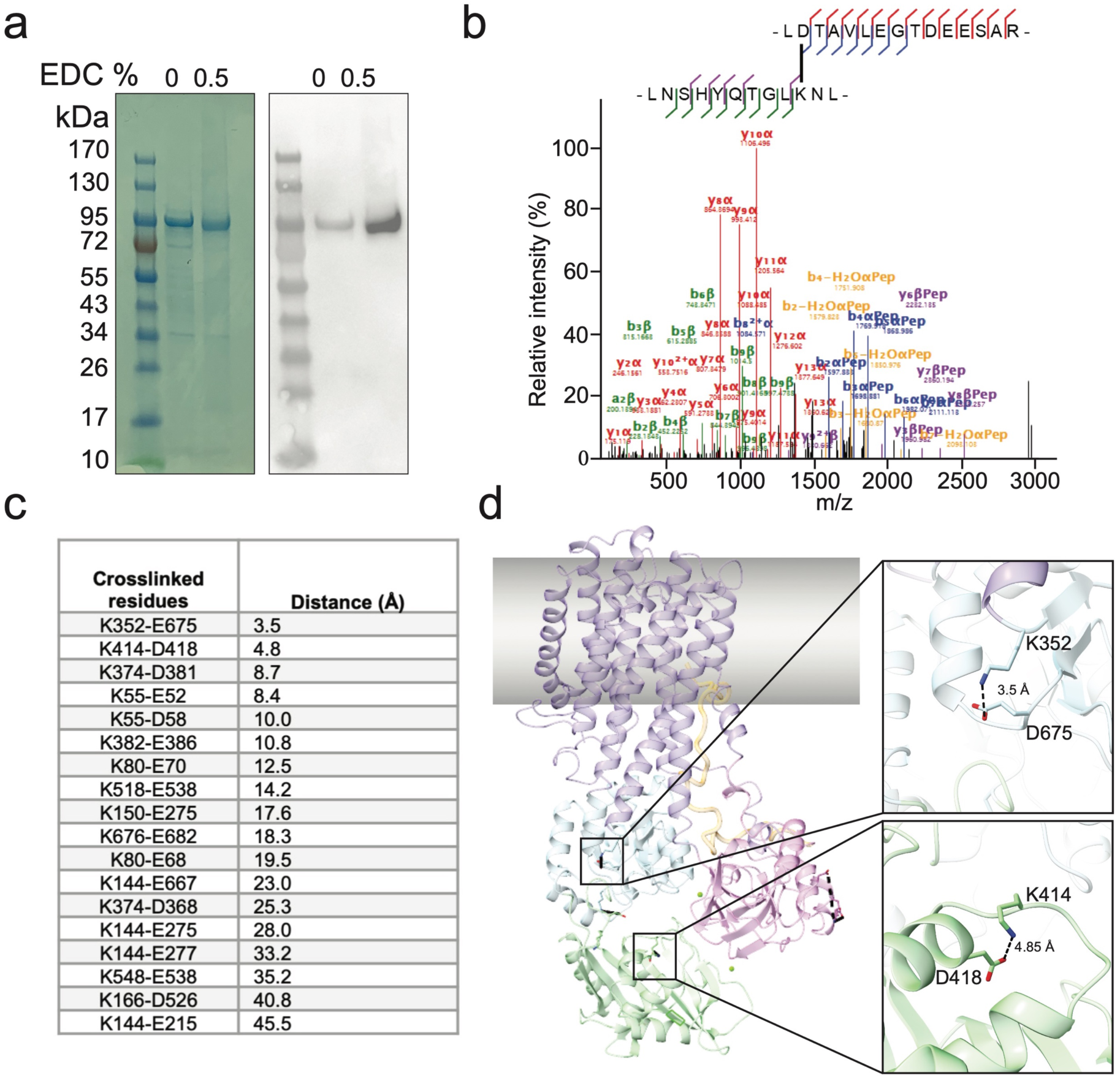
Structural model is supported by crosslinking and mass spectrometry. **a**, MgtAS purified with GDN and crosslinked with EDC. Crosslinked samples were analyzed by SDS-PAGE followed by Coomassie staining (left) or Western blot analysis using α-His_6_ antibodies against MgtA (right). **b**, High-quality MS/MS spectrum for a representative crosslinked peptide listed in panel **c**. Matched b- and y-ions are highlighted for the respective peptides, according to their sequences illustrated above. **c**, Panel of high confidence crosslinks identified by mass spectrometry and their distances measured within a monomeric subunit of the dimeric MgtA structure. **d**, Two representative crosslinks of <10 Å mapped onto a single subunit of the atomic model of the MgtA dimer for viewing purposes. Side chains are displayed in stick representation and colored by heteroatom. The color scheme of the model is based on Fig. 1 with transparency applied for visualization of the crosslinks.

**Extended Data Fig 13.**
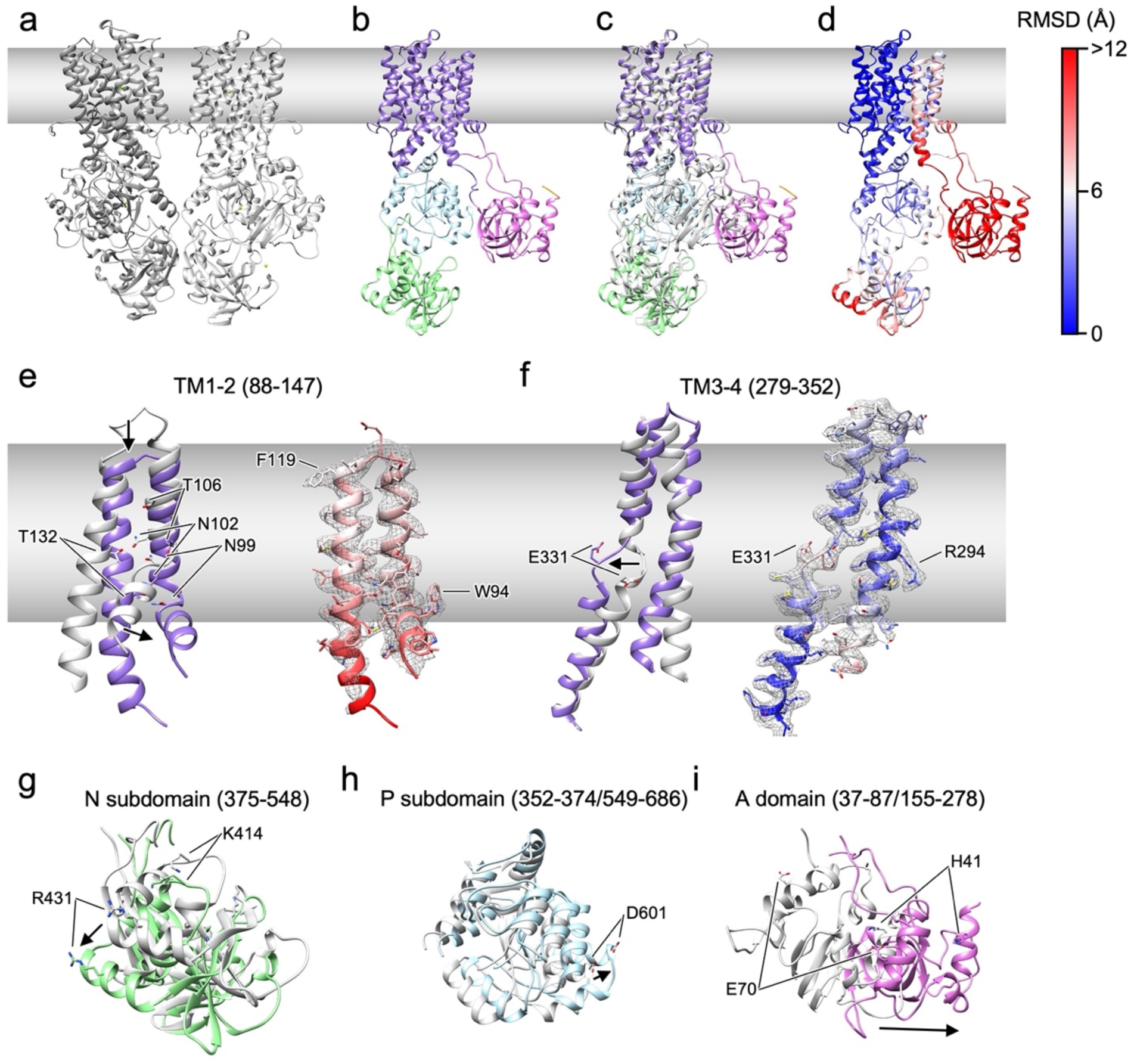
Monomeric MgtA structure adopts a more open conformation relative to the dimer with major changes in TM1-4, N and A. **a**, Dimeric structure with two subunits in two different gray tones and Mg^2+^ ions as green spheres. **b**, Monomeric structure colored as in Fig. 1. **c-d**, To visualize structural changes the colored monomer and a single dimer subunit in gray were superimposed (**c**) and RMSD was calculated. The monomer structure is shown colored by RMSD indicating the largest differences when comparing the two structures by high RMSD values in red. **e-f**, Structural differences between the dimer (gray) and monomer (purple) for TM1-2 (**e**) and TM3-4 (**f**). The monomer colored according to RMSD is on the right with densities in mesh indicating the fit. **g-i**, Structural differences between the soluble domains of the dimer (gray) and monomer (colored).

**Extended Data Fig 14.**
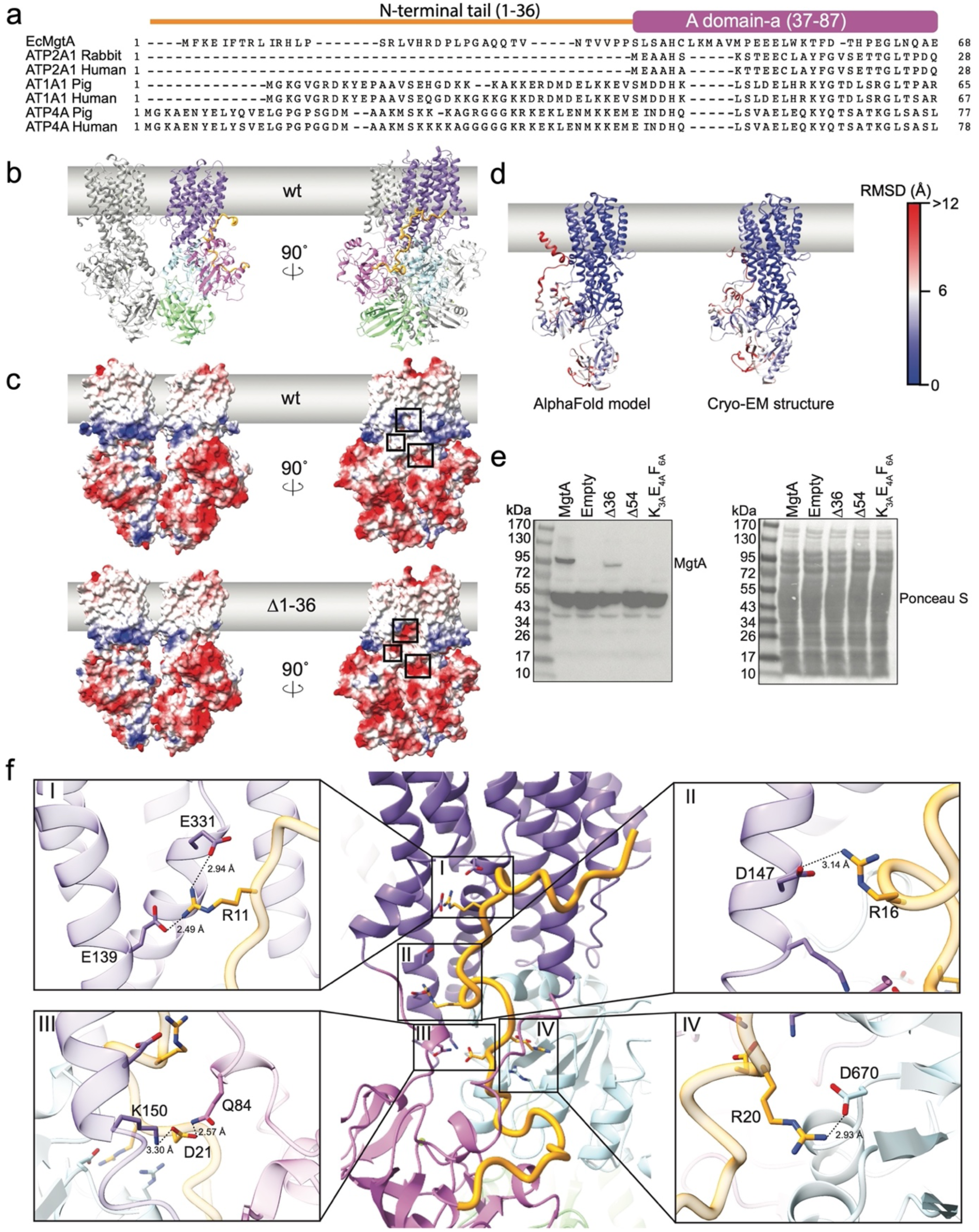
The extended N-terminus forms multi-domain electrostatic interactions between the A, P and TM domains. **a**, Reduced alignment showing an extended N-terminus is present in MgtA, Na^+^/K^+^ ATPases, and H^+^/K^+^ ATPases, but not the Ca^2+^ pump SERCA. The sequence alignment of *E. coli* MgtA (EcMgtA), ATP2A1 (SERCA) from rabbit and human, AT1A1 (Na^+^/K^+^) from pig and human, and ATP4A (H^+^/K^+^) from pig and human was generated using Clustal Omega. Domains are colored and named according to Fig. 1. **b**, Front and side view of the EcMgtA dimer structure colored as in Fig. 1 with the N-terminus highlighted in a thicker loop. The N-terminus of MgtA forms electrostatic interactions with the A, P and TM domains. **c**, Front and side view of the surface representation of the electrostatic potential is displayed for wt and the deletion of the N-terminus (Δ1-36). The structure is rotated 90° to highlight the negatively charged patch that is interacting with the N-terminus. **d**, AlphaFold is unable to predict the N-terminus of MgtA. To visualize structural differences between the AlphaFold model and dimeric MgtA cryo-EM structure the RMSD was calculated between a single subunit. Both the AlphaFold model and a single subunit of the dimeric MgtA structure are shown colored by RMSD. **e**, Mutations of the N-terminus reduce levels of MgtA. Cells were grown uninduced (- IPTG) overnight at 37°C in LB supplemented with 100 mM MgSO_4_ and normalized in lysis buffer prior to western blot analysis with polyclonal anti-MgtA antibodies. Ponceau S-stained membrane serves as a loading control. **f**, Close-up views of sites I-IV which mark salt bridges formed between the N-terminus and the various domains of MgtA.

**Extended Data Fig 15.**
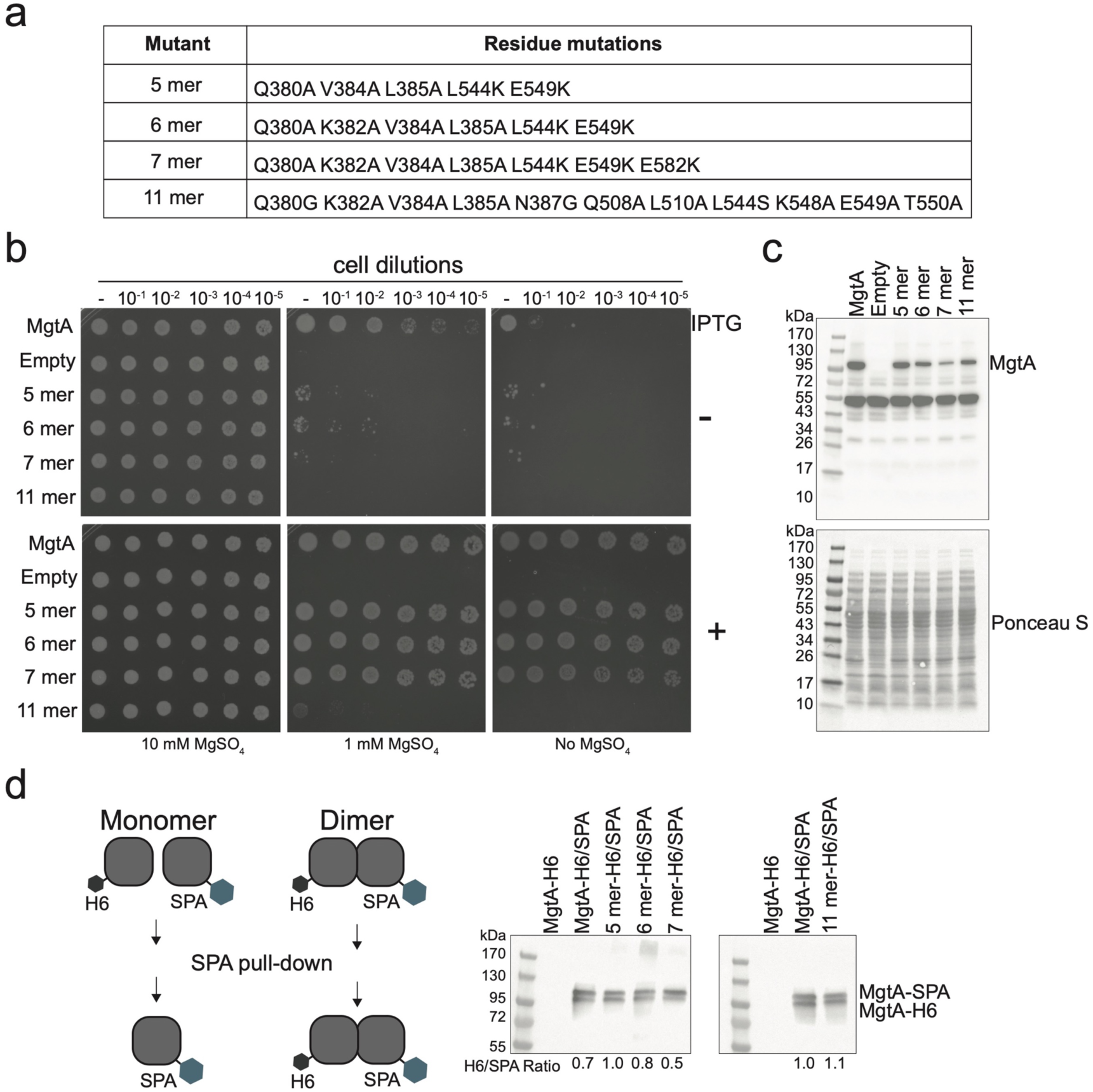
Mutations at the dimer interface impair Mg^2+^ transport. **a**, Summary table of residues mutated at the dimer interface. **b**, Mutations at the dimer interface impair Mg^2+^ ion translocation as visualized by complementation using a Mg^2+^-auxotrophic *E. coli* strain. Overnight cultures were serial diluted and spotted onto LB agar plates supplemented with the indicated concentrations of MgSO_4_ with (+) and without (−) 0.1 mM IPTG for induction and grown at 37°C prior to imaging. **c**, Mutations at the dimer interface reduce levels of MgtA. Cells from the indicated strains were grown uninduced (- IPTG) overnight at 37°C in LB supplemented with 100 mM MgSO_4_ and normalized in lysis buffer prior to Western blot analysis with polyclonal anti-MgtA antibodies. Ponceau S-stained membrane serves as a loading control. **d**, The dimer is resistant to directed mutations. Dimer mutants were copurified using differentially tagged MgtA derivatives (as described in Fig. 2b). Proteins were visualized by Western blot analysis using polyclonal anti-MgtA antibodies.

**Extended Data Fig 16.**
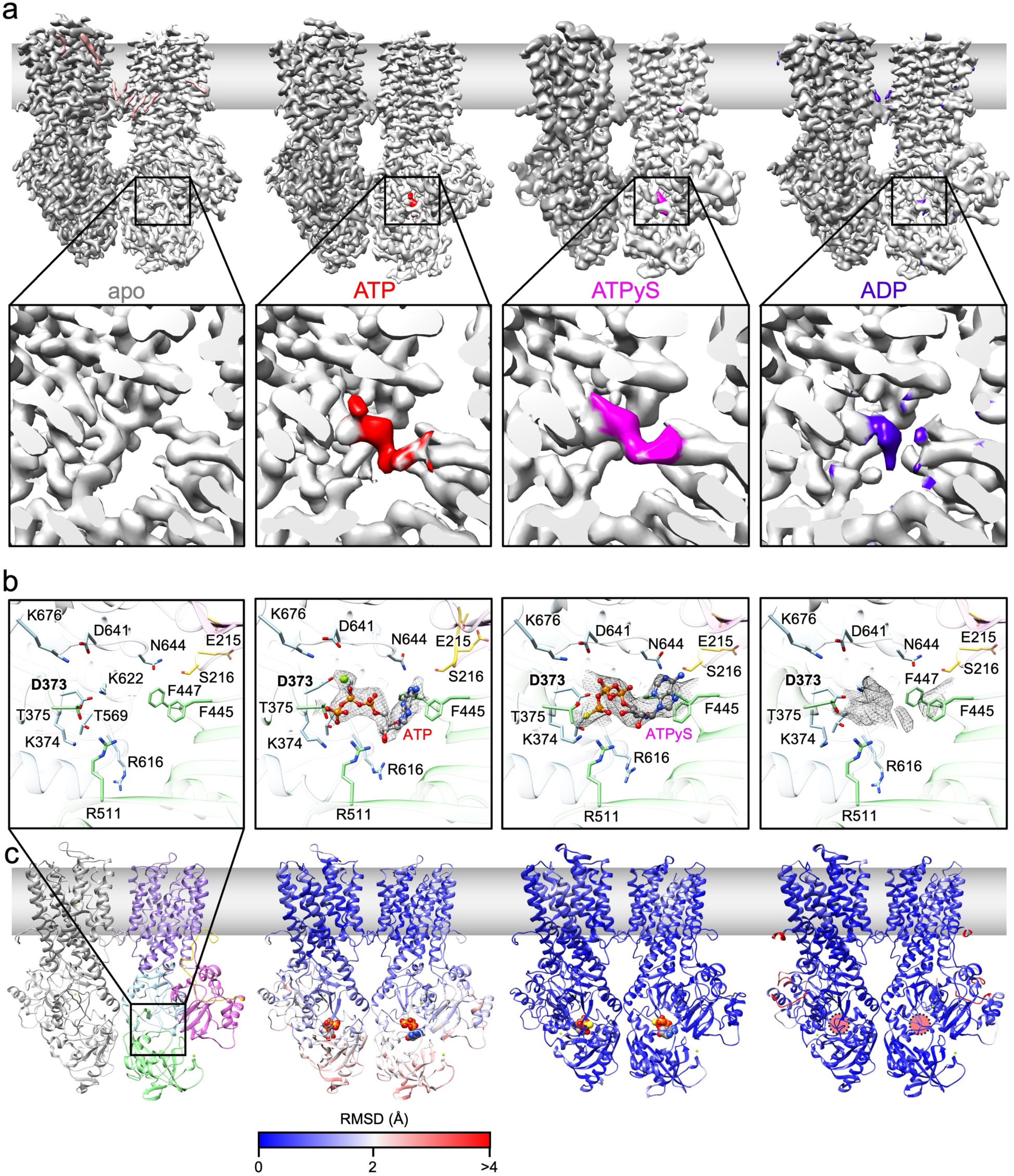
Cryo-EM of dimeric MgtA bound to nucleotides reveals extra density in the nucleotide binding pocket. **a**, Side view of cryo-EM maps of the EcMgtA dimer in the presence of 5 mM MgCl_2_, and nucleotides. The map of the apo structure is the same as in Fig. 1. Maps are shown in gray and the extra densities are highlighted in color corresponding to the respective nucleotides for MgtA in the presence of 5 mM ATP (red), 5 mM ATPγS (pink), and 5 mM ADP (purple). The average resolution of the respective maps are: apo 2.93 Å, ATP 3.72 Å, ATPγS 3.87 Å, and ADP 3.75 Å. **b**, A zoom in into the nucleotide binding site with selected residues shown in stick representation while the nucleotides are in ball-and-stick. Cryo-EM density near the nucleotide is shown in gray mesh. **c**, Dimeric structural models colored according to RMSD when compared to the apo dimeric structure and nucleotides in spheres. For the structure with ATP, residues F447, F445 and N415 from the nucleotide binding subdomain interact with the adenine group of ATP, while F445 appears to form a pi-pi interaction with the adenine and N415 a hydrogen bond with nitrogen N7 of the ATP adenine component. An additional small, isolated density 3 Å from the N6 nitrogen might be a water molecule or ion. Nucleotide binding subdomain residue R511 interacts with the ATP ribose component through a water molecule. Several residues, including D373, K374, and T375 in the phosphorylation subdomain, which form the conserved DxT catalytic motif required for ATP hydrolysis, as well as T569, D571, and K622, all of which interact with the β- and γ-phosphate groups of the ATP molecule. A continuous density to T375 and T569 indicates a close interaction between the γ-phosphate and these two residues. A Mg^2+^ was assigned to a strong round density observed near the β-phosphate. This is near D373, which is known to be the phosphorylated catalytic residue.

**Extended Data Fig 17.**
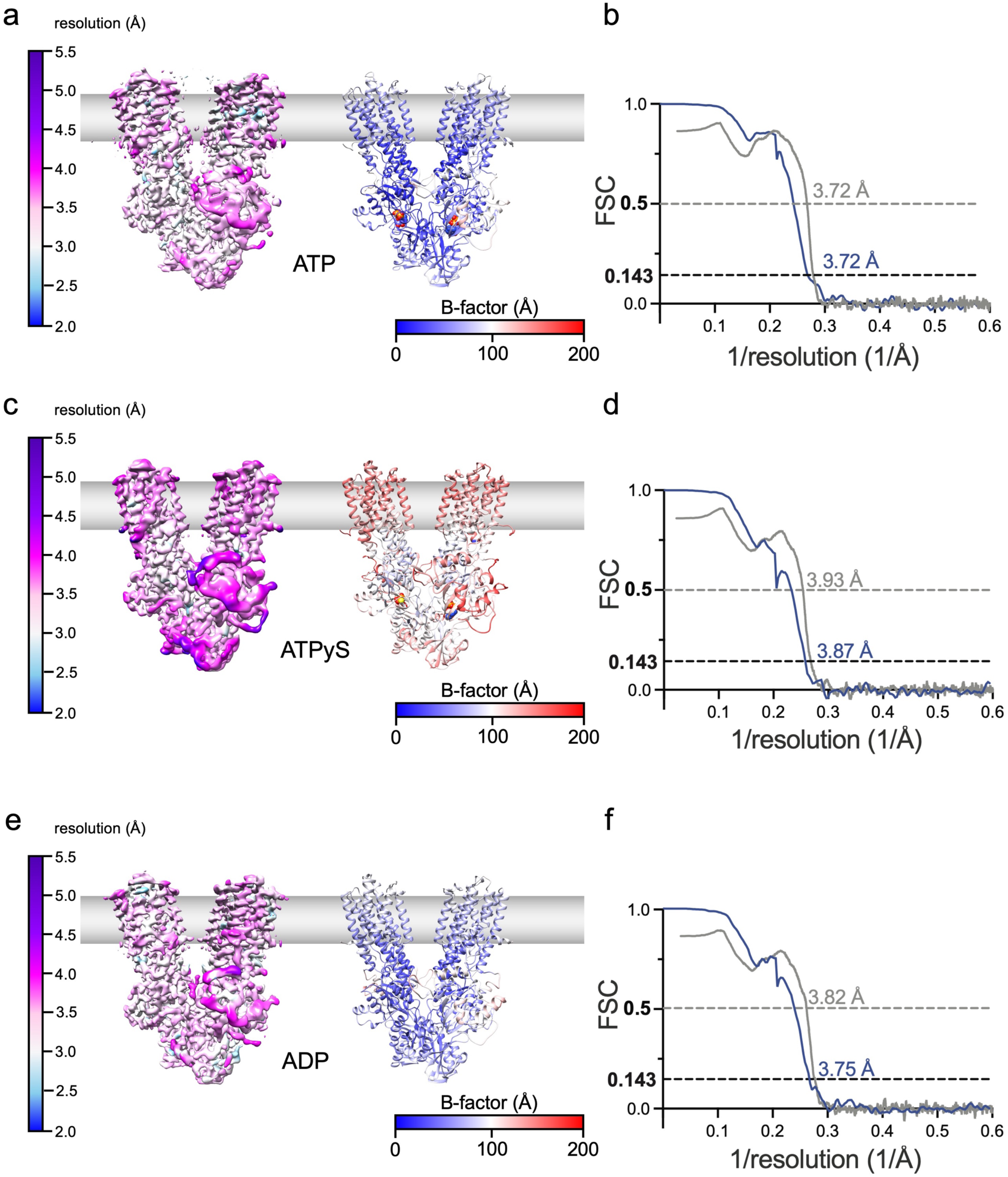
Local resolution and average resolution of the dimeric MgtA with nucleotides and B-factor distribution of models. **a**, Final dimer reconstruction in the presence of 5 mM ATP filtered and colored to local resolution (left) and fitted model colored according to B-factor distribution (right) indicating rigid and more flexible regions of the complex. **b**, Fourier Shell Correlation (FSC) curve of the final dimer reconstruction of MgtA with ATP in blue indicating an average resolution of 3.72 Å according to the FSC=0.143 criterion. FSC between the final ATP dimer map and fitted model is shown in gray. **c**, Final dimer reconstruction in the presence of 5 mM ATPγS filtered and colored to local resolution (left) and fitted model colored according to B-factor distribution (right) indicating rigid and more flexible regions of the complex. **d**, FSC curve of the final dimer reconstruction of MgtA with ATPγS in blue indicating an average resolution of 3.87 Å according to the FSC=0.143 criterion. FSC between the final ATPγS dimer map and fitted model is shown in gray. **e**, Final dimer reconstruction in the presence of 5 mM ADP filtered and colored to local resolution (left) and fitted model colored according to B-factor distribution (right) indicating rigid and more flexible regions of the complex. **f**, FSC curve of the final dimer reconstruction of MgtA with ADP in blue indicating an average resolution of 3.75 Å according to the FSC=0.143 criterion. FSC between the final ADP dimer map and fitted model is shown in gray.

**Extended Data Fig 18.**
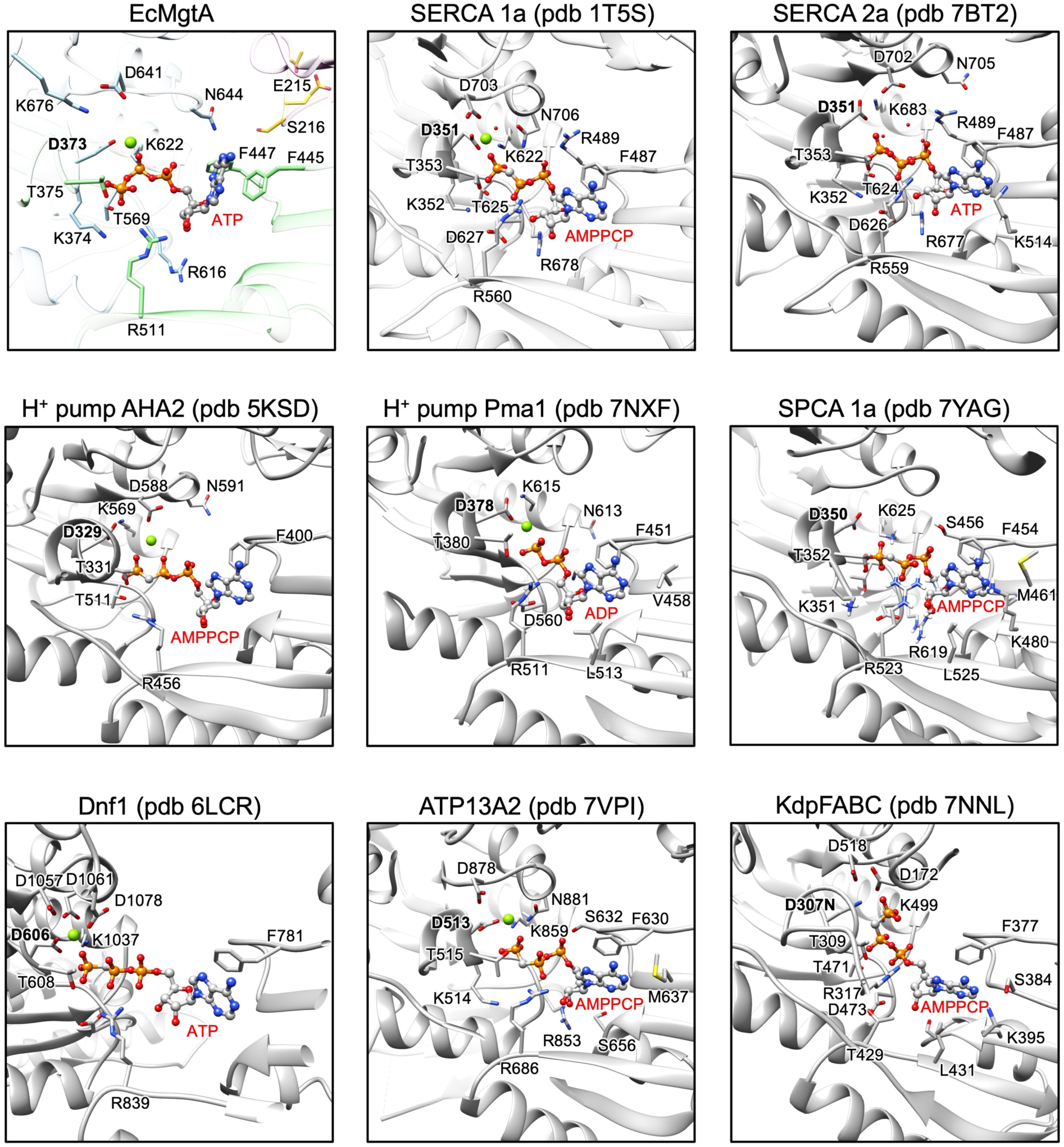
The nucleotide binding pocket of MgtA and other P-type ATPases are structurally similar. Zoom in into the nucleotide binding pocket of EcMgtA colored as in Fig 1 and 3. Superimposed nucleotide binding pockets of eight other P-type ATPases are shown in gray. X-ray structure of Ca^2+^ pump SERCA 1a with AMPPCP (PDB 1T5S), X-ray structure SERCA 2a with ATP (PDB 7BT2), X-ray structure of proton pump AHA2 with AMPPCP (PDB 5KSD), X-ray structure of proton pump Pma1 with ADP (PDB 7NXF), Cryo-EM structure of Mn^2+^ and Ca^2+^ pump SPCA with AMPPCP (PDB 7YAG), X-ray structure of phospholipid transporter Dnf1 with ATP (PDB 6LCR), cryo-EM structure of inorganic ion transporter ATP13A2 with AMPPCP (PDB 7VPI), and cryo-EM structure of bacterial potassium pump KdpFABC complex with AMPPCP (PDB 7NNL). Residues near the nucleotide binding pocket are displayed as stick, nucleotides as ball-and-stick, Mg^2+^ ions as green spheres.

**Extended Data Fig 19.**
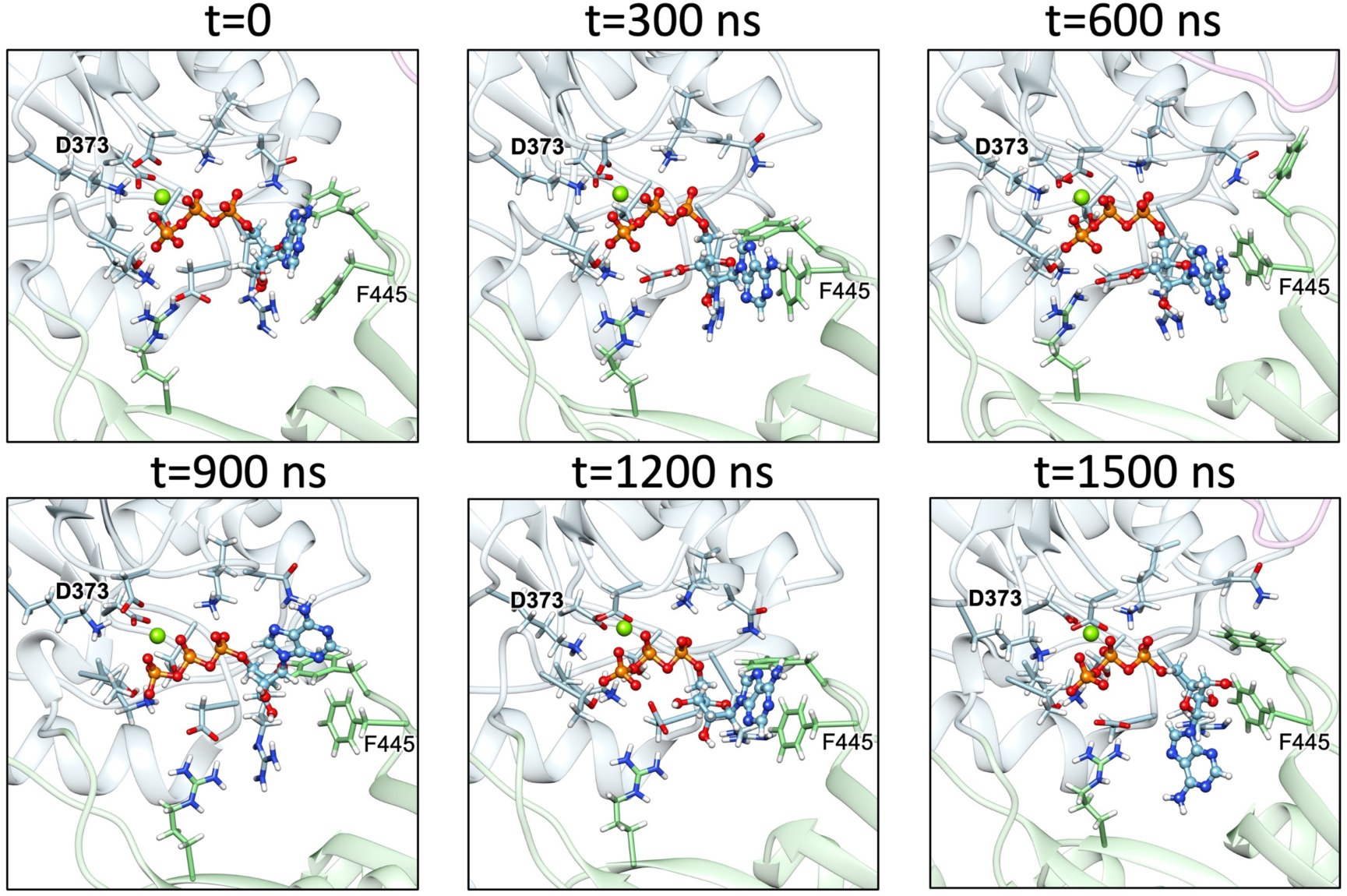
Time sequence of MD simulations of the MgtA dimer with ATP. Six representative conformations of the ATP binding pocket during time points in the molecular simulation. The ATP and coordinating residues are displayed according to Extended Data Fig 18.

**Extended Data Fig 20.**
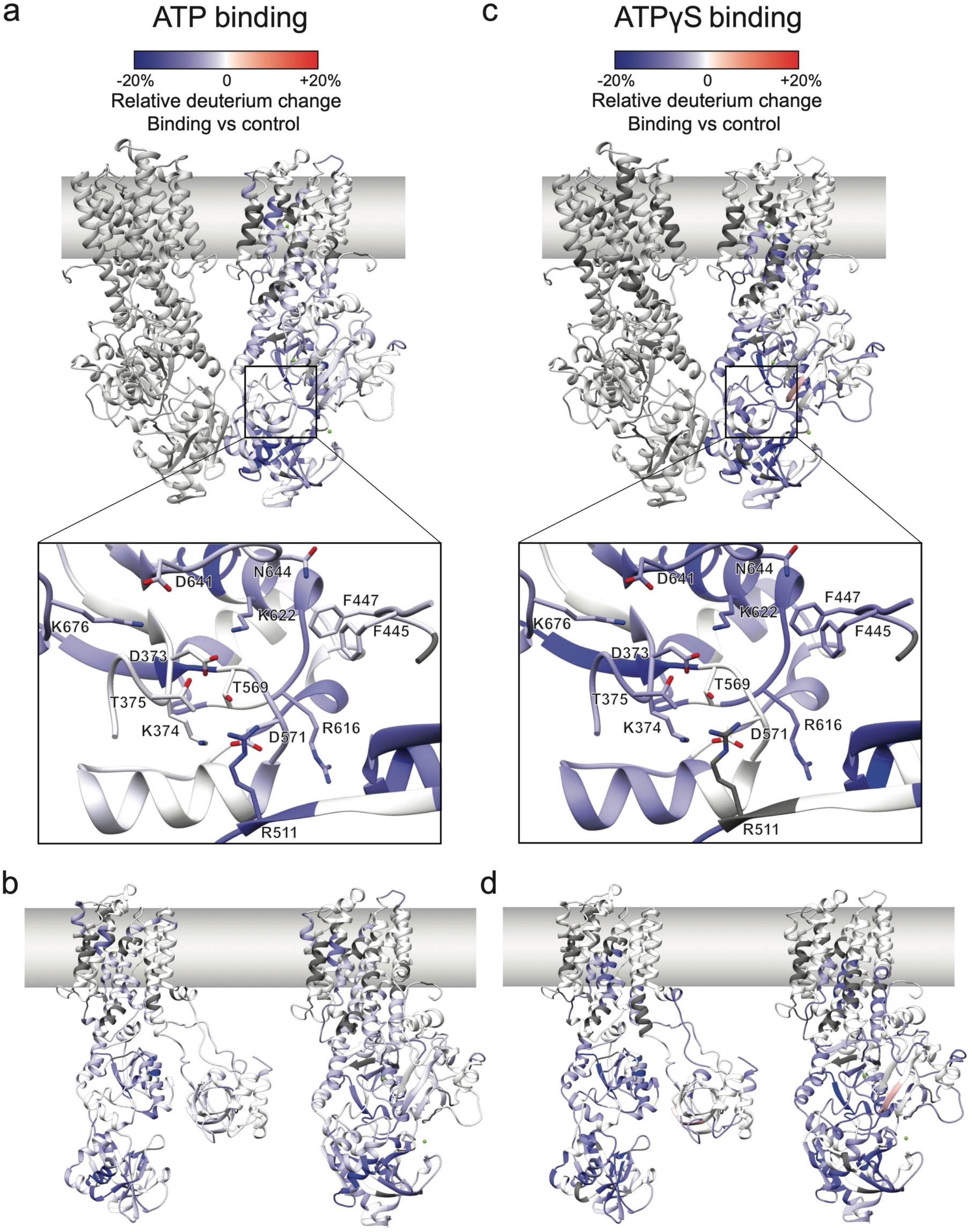
HDX-MS analysis of MgtA reveals structural changes upon ATP and ATPγS binding. **a**, HDX-MS results of 5 mM ATP binding to MgtA after 2 h of deuterium labeling. MgtA is shown as a homodimer in parallel to the membrane in ribbon representation. The ribbon representation has one subunit gray and one subunit of the dimer colored based on uptake differentials. Zoom in for the indicated region and display of residues involved in ATP binding. Atoms are colored by heteroatoms and dark gray indicates no peptides were detected. **b**, Comparison of HDX-MS uptake differentials for ATP binding to MgtA mapped onto the monomer and dimer structures. Side view of monomer (left) and one subunit of the dimer (right) MgtA structure colored by HDX-MS uptake differentials between −/+ ATP. **c**, HDX-MS results of 5 mM ATPγS binding to MgtA after 2 h of deuterium labeling. MgtA is shown as a homodimer in parallel to the membrane in ribbon representation. The ribbon representation has one subunit gray and one subunit of the dimer colored based on uptake differentials. Zoom in for the indicated region and display of residues that coordinate ATPγS. Atoms are colored by heteroatoms. **d**, Comparison of HDX-MS uptake differentials for ATPγS binding to MgtA mapped onto the monomer and dimer structures. Side view of monomer (left) and one subunit of the dimer MgtA structure (right) colored by HDX-MS uptake differentials between −/+ ATPγS. For all panels, decreased deuterium uptake is shown in blue, and increased deuterium uptake is shown in red.

**Extended Data Fig. 21.**
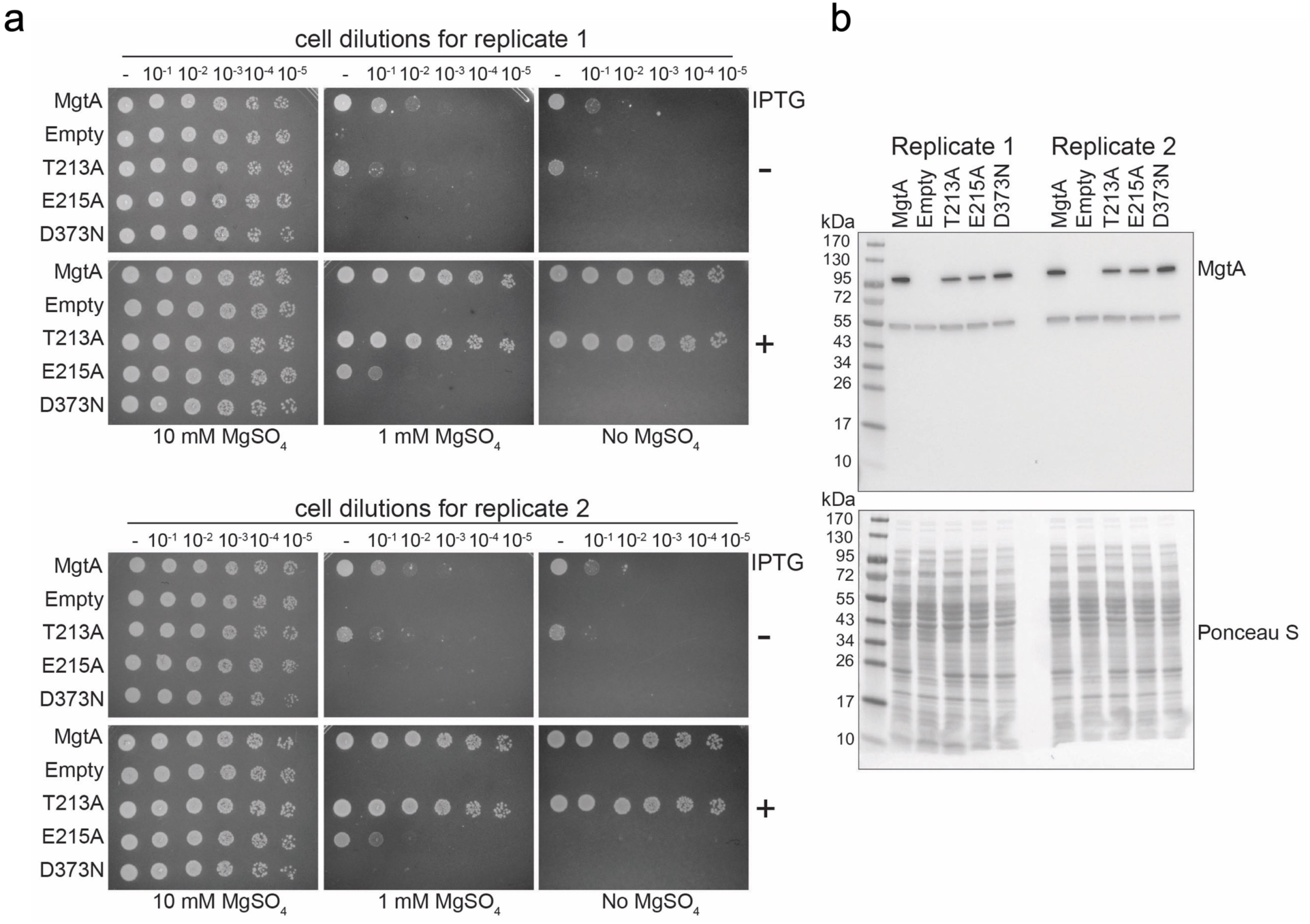
Residues D373 and E215 are required for Mg^2+^ transport. **a**, MgtA_D373N_ does not complement and MgtA_E215A_ only slightly complements a Mg^2+^-auxotrophic *E. coli* strain. Overnight cultures were serial diluted and spotted onto LB agar plates supplemented with the indicated concentrations of MgSO_4_ with (+) and without (−) 0.1 mM IPTG for induction and grown at 37°C prior to imaging. **b**, MgtA proteins with mutated residues involved in ATP hydrolysis are expressed at levels comparable to the wild-type protein. Cells from the indicated strains were grown uninduced (- IPTG) overnight at 37°C in LB supplemented with 100 mM MgSO_4_ and normalized in lysis buffer prior to Western blot analysis with polyclonal anti-MgtA antibodies. Ponceau S-stained membrane serves as a loading control. Results from biological replicates (2n) are shown.

**Extended Data Fig. 22.**
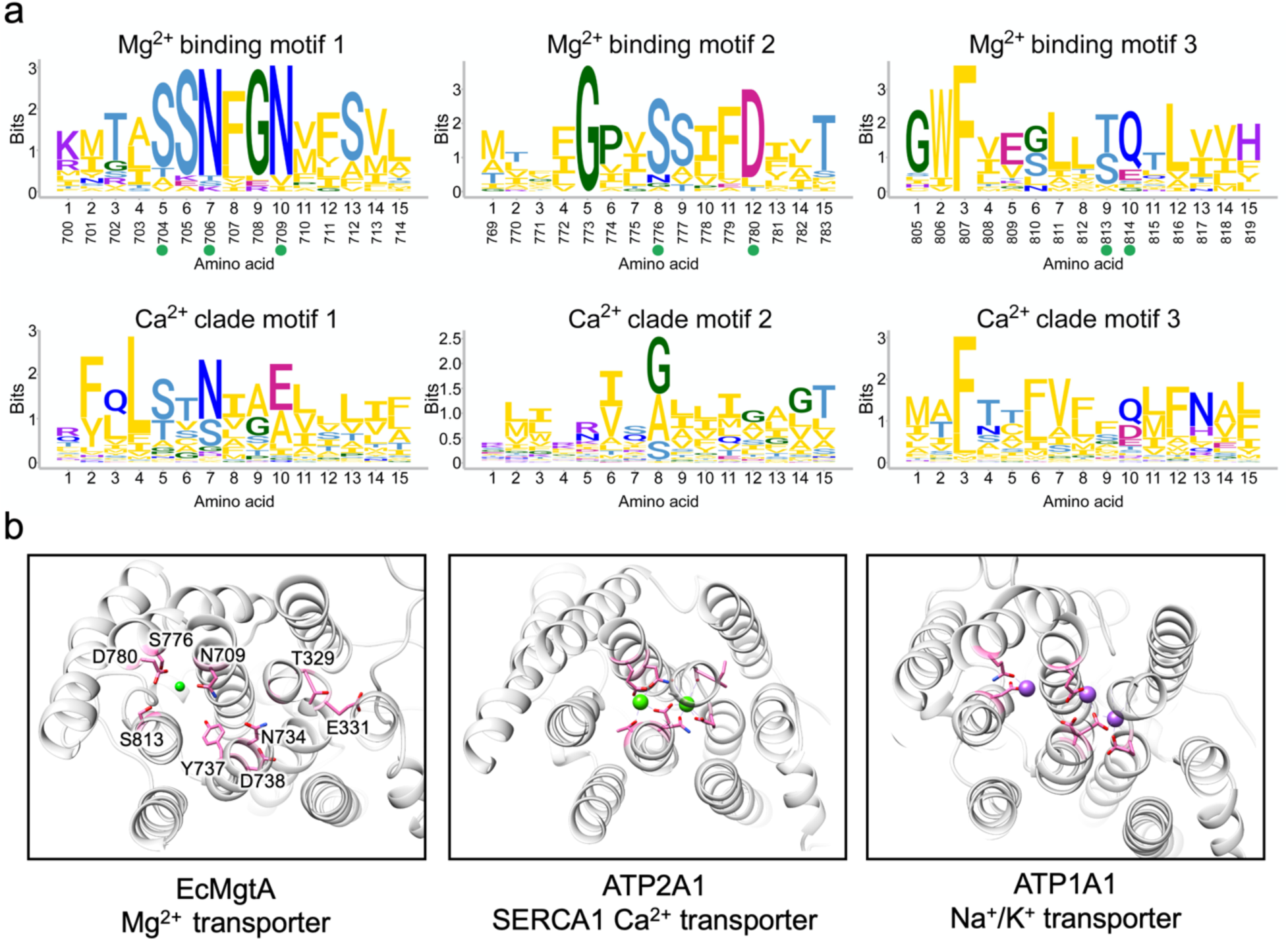
Conservation of key Mg^2+^ binding residues. **a**, Sequence logos displaying conservation of amino acids in different members of the P-type ATPase transporters. Letters represent amino acid abbreviations; the height of each letter represents the relative probability of conservation among members of the P-type ATPase family. Logos correspond to the Mg^2+^ TM binding sites to illustrate residues conserved in the MgtA clade compared to the Ca^2+^ clade. The sequence logos are also highlighted in the reduced multisequence alignment in Extended Data Fig. 1. **b**, Structural comparison of ion-bound P-type ATPases. One subunit of the dimeric EcMgtA transporter is shown with corresponding regions of ATP2A1 (SERCA1 Ca^2+^ transporter PDB 2ZBD) and ATP1A1 (Na^+^/K^+^ transporter PDB 4HQJ). Residues of EcMgtA predicted to be involved in ion binding are colored in pink and the Mg^2+^ ion is colored green. Residues of ATP2A1 and ATP1A1 involved in ion binding are colored in pink and the Ca^2+^ and Na^+^ ions are colored green and purple, respectively.

**Extended Data Fig. 23.**
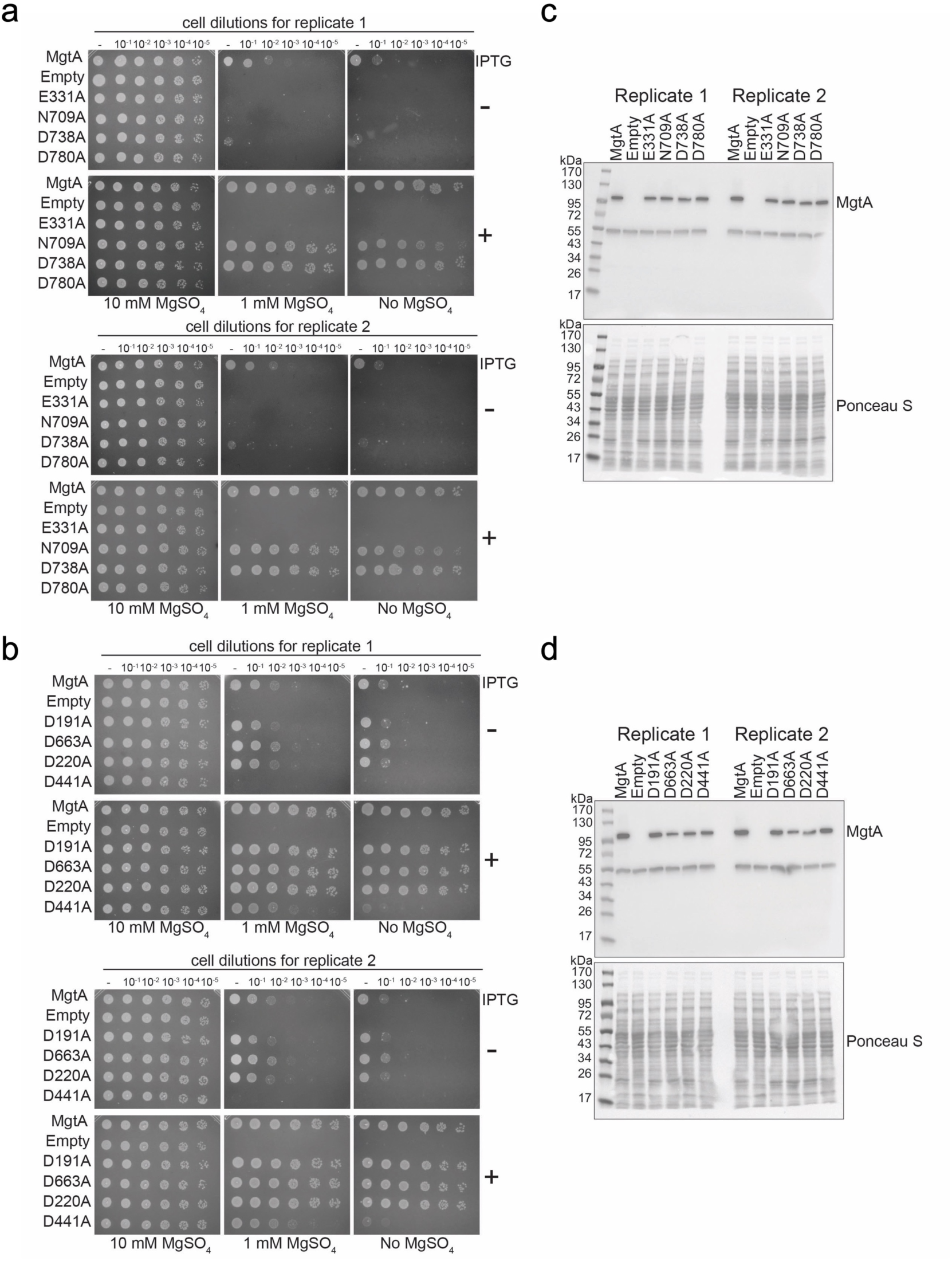
Functional analysis of key Mg^2+^ binding residues. **a-b**, Complementation assay using a Mg^2+^-auxotrophic *E. coli* strain and EcMgtA with mutations of Mg^2+^ binding residues contained within TM segments (N709, D780) and based on sequence conservation and structural comparison predicted residues (E331, D738) (**a**) or soluble domain (D191, D663, D220, D441) (**b**). Overnight cultures were serial diluted and spotted onto LB agar plates supplemented with the indicated concentrations of MgSO_4_ with (+) and without (−) 0.1 mM IPTG for induction and grown at 37°C prior to imaging. **c-d**, Levels of MgtA with mutations of Mg^2+^ binding residues contained within TM segments (E331, N709, D738, D780) (**c**) or soluble domain (D191, D663, D220, D441) (**d**). Cells from the indicated strains were grown uninduced (- IPTG) overnight at 37°C in LB supplemented with 100 mM MgSO_4_ and normalized in lysis buffer prior to western blot analysis with polyclonal anti-MgtA antibodies. Ponceau S-stained membrane serves as a loading control. Results from biological replicates (2n) are shown.

**Extended Data Fig. 24.**
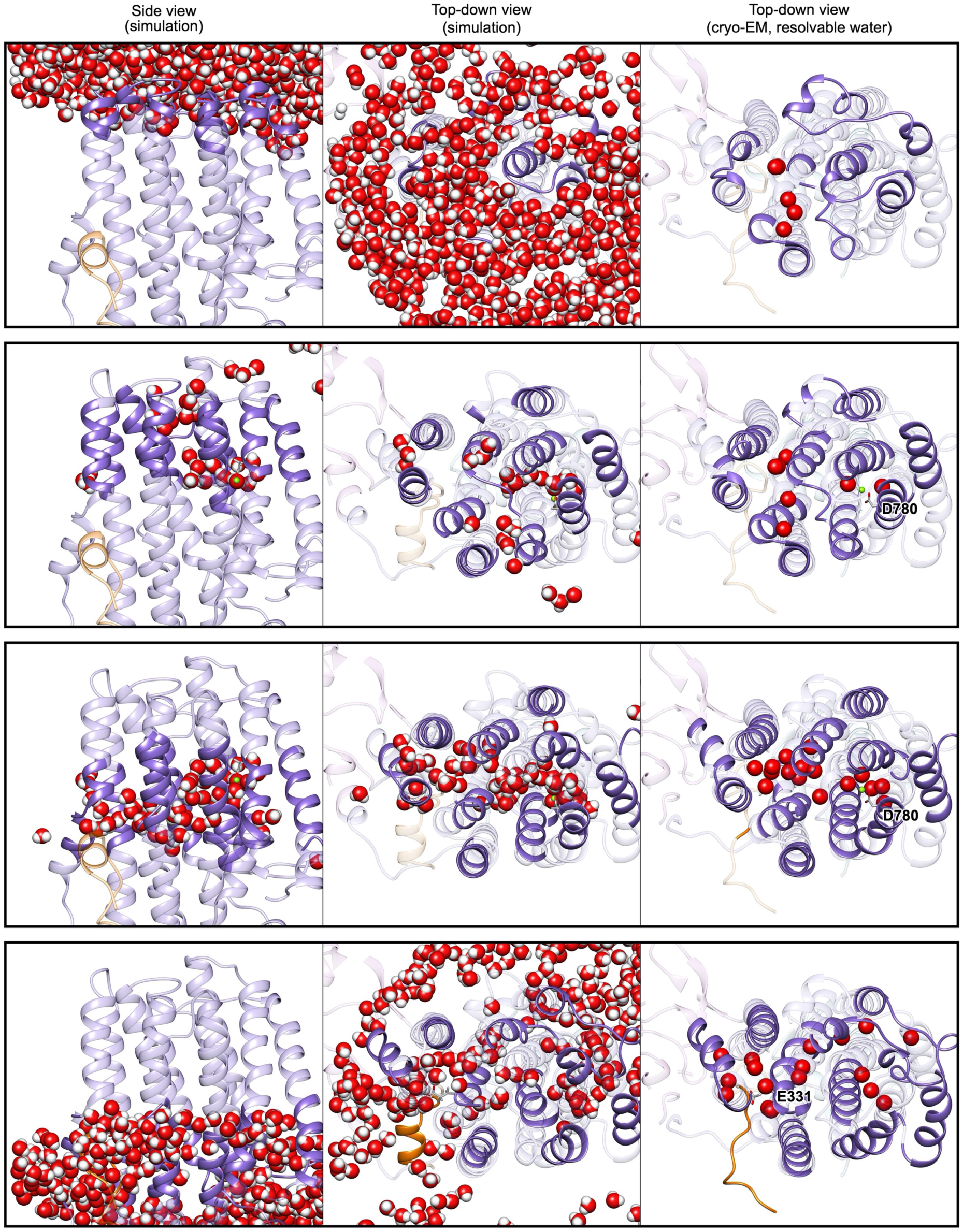
Water accessibility in the TM domain of MgtA. Figures are arranged such that rows correspond with overlapping 1.5 nanometer thick cuts spaced every 1 nanometer. The left column is the side view from the simulation, while the middle column is from the top-down starting on the periplasmic side of the transporter. For the simulation, waters (including hydrogens) are shown in sphere representation. At right is the corresponding top-down view of the MgtA dimer from cryo-EM, with resolved water molecules shown as red spheres. The protein ribbon model is shown opaque through the cut of the simulation, while outside of the cut waters are not shown, and the protein is transparent. The transmembrane Mg^2+^ as well as glutamic and aspartic acid residues 331 and 780 (respectively) are shown in stick representation.

**Extended Data Fig. 25.**
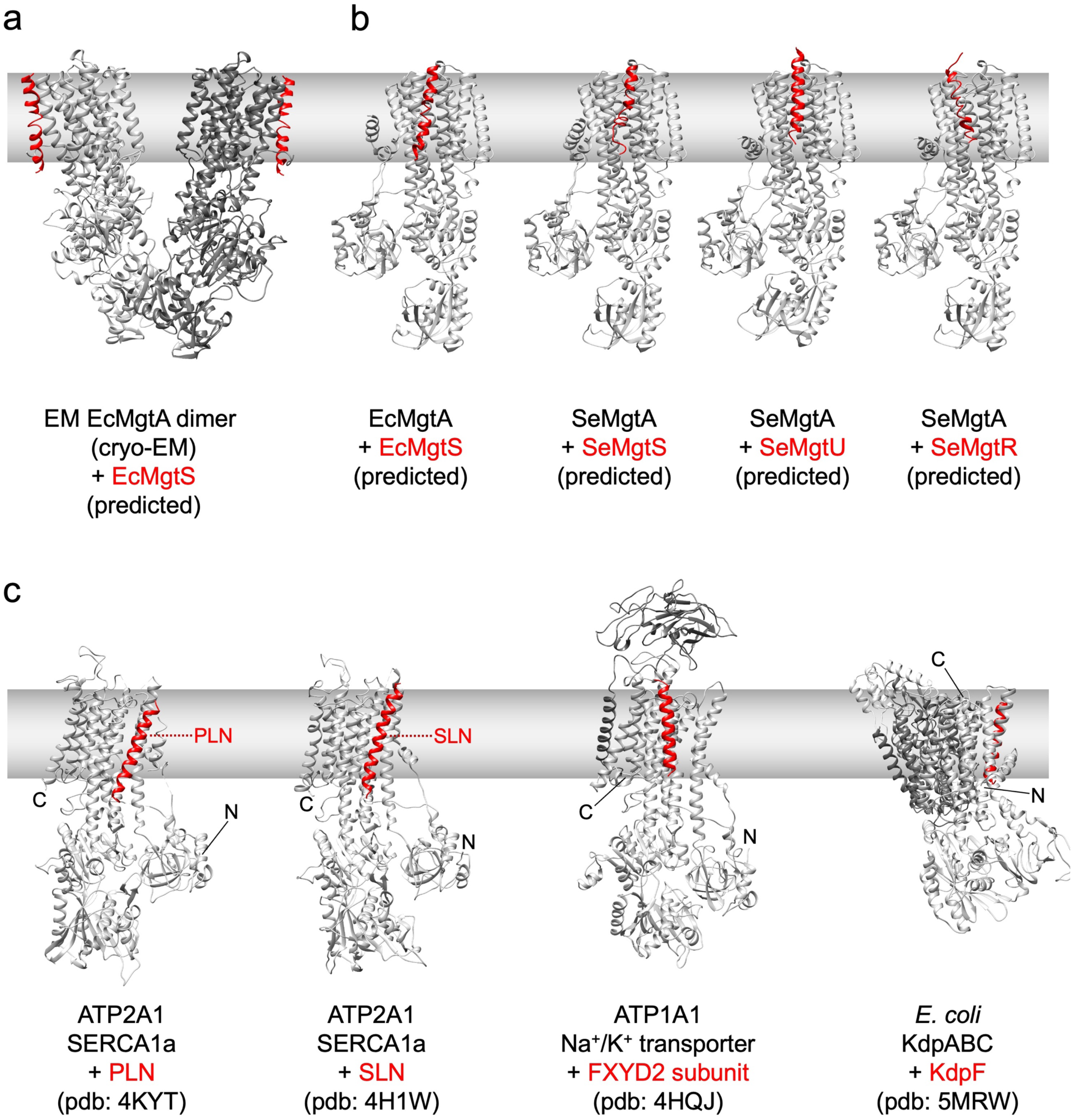
Predicted and documented interactions of small proteins with P-type ATPase proteins. **a**, Dimer cryo-EM structure of *E. coli* MgtA from this study with *E. coli* MgtS binding predicted by AlphaFold Multimer beta. **b**, Models of *E. coli* MgtA monomer and *E. coli* MgtS and *S. enterica* MgtA and *S. enterica* MgtS, MgtU and MgtR predicted by AlphaFold Multimer beta. **c**, Selected structures of indicated P-type ATPases solved with small α-helical proteins. P-type ATPases are in gray with small protein in red.

### Extended Data Tables

**Extended Data Table 1.** Residues involved in dimer interface.

| **Residue chain A** | **Residue chain B** | **Interaction Type** |
| --- | --- | --- |
| Q380 | Q380 | Electrostatic/h-bond |
| K382 | E582 | Electrostatic |
| N387 | Q508 | Electrostatic/h-bond |
| K548 | E549 | Electrostatic |
| E386 | R578 | Electrostatic<br>(in simulation) |
| V384 | L544 | Hydrophobic |
| V384 | P546 | Hydrophobic |
| V384 | P547 | Hydrophobic |
| L385 | L385 | Hydrophobic |
| L385 | L510 | Hydrophobic |
| L385 | L544 | Hydrophobic |
| L544 | L544 | Hydrophobic |
| P546 | P546 | Hydrophobic |

**Extended Data Table 2.** Summary of all mutants generated and their outcome.

| **Mutation** | **Functional Significance** | **Outcome** |
| --- | --- | --- |
| D373N | DxT motif, ATP hydrolysis | No Activity |
| T213A | TGES loop, ATP hydrolysis | Reduced activity mild |
| E215A | TGES loop, ATP hydrolysis | Reduced activity severe |
| E331A | TM Mg<sup>2+</sup> binding site | No activity |
| D738A | TM Mg<sup>2+</sup> binding site | Reduced activity mild |
| D780A | TM Mg<sup>2+</sup> binding site | No activity |
| N709A | TM Mg<sup>2+</sup> binding site | Reduced activity mild |
| D191A | Site II cytoplasmic Mg<sup>2+</sup> binding | Active |
| D663A | Site II cytoplasmic Mg<sup>2+</sup> binding | Active |
| E220A | Site III cytoplasmic Mg<sup>2+</sup> binding | Active |
| D441A | Site III cytoplasmic Mg<sup>2+</sup> binding | Reduced activity severe |
| Q380A, V384A, L385A, L544K, E549K | Dimer interface | Reduced activity mild |
| Q380A, K382A, V384A, L385A, L544K, E549K | Dimer interface | Reduced activity mild |
| Q380A, K382A, V384A, L385A, L544K, E549K, E582K | Dimer interface | Reduced activity mild |
| Q380A, K382A, V384A, L385A, N387G, Q508A, L510A, L544S, K548A, E549A, T550A | Dimer interface | No activity |
| Δ36 | N-terminus | Unstable |
| Δ54 | N-terminus | Unstable |
| K3A, E4A, F6A | N-terminus | Unstable |

**Extended Data Table 3.** Cryo-EM data collection parameters and analysis.

|  | **Dataset 1** | **Dataset 2** | **Dataset 3** | **Dataset 4** | **Dataset 5** | **Dataset 6** | **Dataset 7** |
| --- | --- | --- | --- | --- | --- | --- | --- |
| **Date** | 2021/09/16-19 | 2021/10/29-31 | 2022/08/01-02 | 2022/08/23-25 | 2022/08/03-04 | 2022/10/28-30 | 2022/09/26-29 |
| **Protein concentration** | 3.2 mg/mL | 3.2 mg/mL | 2.3 mg/mL | 2.3 mg/mL | 2.3 mg/mL | 2.3 mg/mL | 2.3 mg/mL |
| **Ligands** | (Mg) | (Mg) | Mg-ATPγS | Mg-ATPγS | Mg-ADP | Mg-ADP | Mg-ATP |
| **Sample volume** | 3 μl | 3 μl | 3 μl | 3 μl | 3 μl | 3 μl | 3 μl |
| **Grid type** | QF R1.2/1.3<br>400 Cu mesh | QF R1.2/1.3<br>400 Cu mesh | QF R1.2/1.3<br>400 Cu mesh | QF R1.2/1.3<br>400 Cu mesh | QF R1.2/1.3<br>400 Cu mesh | QF R1.2/1.3<br>400 Cu mesh | QF R1.2/1.3<br>400 Cu mesh |
| **Plunge freezer** | Leica EM<br>GP2 | Leica EM<br>GP2 | Leica EM<br>GP2 | Leica EM<br>GP2 | Leica EM<br>GP2 | Leica EM<br>GP2 | Leica EM<br>GP2 |
| **Blotting time (s)** | 5 | 4 | 6 | 6 | 6 | 6 | 6 |
| **Temperature (°C)** | 5 | 5 | 4 | 4 | 4 | 4 | 4 |
| **humidity set (measured)** | 95%<br>(77-88%) | 95%<br>(77-88%) | 95%<br>(77-88%) | 95%<br>(77-88%) | 95%<br>(77-88%) | 95%<br>(77-88%) | 95%<br>(77-88%) |
| **Microscope** | Titan Krios<br>G1 | Titan Krios<br>G1 | Titan Krios<br>G1 | Titan Krios<br>G1 | Titan Krios<br>G1 | Titan Krios<br>G1 | Titan Krios<br>G4 |
| **Voltage (kV)** | 300 | 300 | 300 | 300 | 300 | 300 | 300 |
| **Camera** | K3 (CDS<br>mode) | K3 (CDS<br>mode) | K3 (CDS<br>mode) | K3 (CDS<br>mode) | K3 (CDS<br>mode) | K3 (CDS<br>mode) | K3 (CDS<br>mode) |
| **Energy filter (slit)** | Yes (20 eV) | Yes (20 eV) | Yes (20 eV) | Yes (20 eV) | Yes (20 eV) | Yes (20 eV) | Yes (20 eV) |
| **Cs corrector** | No | No | No | No | No | No | No |
| **Objective aperture** | C2 100 μm | C2 70 μm | C2 100 μm | C2 100 μm | C2 100 μm | C2 100 μm | No |
| **Magnification** | 105,000 x | 105,000 x | 105,000 x | 105,000 x | 105,000 x | 105,000 x | 105,000 x |
| **Physical pixel size (Å/px)** | 0.83 | 0.83 | 0.83 | 0.83 | 0.83 | 0.83 | 0.85 |
| **Super-resolution pixel size (Å/px)** | 0.415 | 0.415 | 0.415 | 0.415 | 0.415 | 0.415 | 0.425 |
| **Electron exposure (e<sup>-</sup>/Å<sup>2</sup>)** | 60 | 50 | 50 | 50 | 50 | 50 | 50 |
| **Number of movie frames** | 60 | 50 | 50 | 50 | 50 | 50 | 60 |
| **Dose rate (e<sup>-</sup>/px/s)** | 9.5 (~7.5 on camera) | 10 (~7.5 on camera) | 9 (~7.5 on camera) | 9 (~7.5 on camera) | 9 (~7.5 on camera) | 9 (~7.5 on camera) | 9 (~7.5 on camera) |
| **Defocus (μm)** | -1 to -2 | -1 to -2 | -0.7 to -1.8 | -0.7 to -1.8 | -0.8 to -1.5 | -0.8 to -1.5 | -0.8 to -1.8 |
| **Number of total micrographs** | 10,246 | 6,906 | 3,396 | 6,472 | 2,519 | 7,107 | 14,585 |
| **Number of selected micrographs** | 10,164 | 5,778 | 2,849 | 6,337 | 1,897 | 7,082 | 11,446 |
| **Number of particles picked** | 1,578,214 | 1,775,726 | 454,311 | 1,329,014 | 302,981 | 1,153,466 | 2,339,956 |
| **Number of MgtA dimer particles** | 497,633 | 149,351 | 22,495 | 43,993 | 16,962 | 48,407 | 94,029 |
| **Number of MgtA monomer particles** | 609,684 | 323,129 | N/A | N/A | N/A | N/A | N/A |

**Extended Data Table 4.** Cryo-EM map and model analysis.

|  | MgtA<br>dimer<br>(C2)<br>EMD-42794<br>PDB 8UY7 | MgtA<br>dimer<br>(C1)<br>EMD-42795<br>PDB 8UY8 | MgtA<br>monomer<br>(C1)<br>EMD-42796<br>PDB 8UY9 | MgtA<br>dimer<br>with ATPyS<br>(C2)<br>EMD-42798<br>PDB 8UYB | MgtA<br>dimer<br>with ADP<br>(C2)<br>EMD-42799<br>PDB 8UYC | MgtA<br>dimer<br>with ATP<br>(C2)<br>EMD-42797<br>PDB 8UYA |
| --- | --- | --- | --- | --- | --- | --- |
| **Number of particles in map** | 160,139 | 160,139 | 78,231 | 28,120 | 28,666 | 48,080 |
| **Final pixel size used for**<br>**final maps (Å/px)** | 0.83 | 0.83 | 0.83 | 0.83 | 0.83 | 0.85 |
| **Resolution of map (Å)** | 2.93 | 3.03 | 3.65 | 3.87 | 3.75 | 3.72 |
| **Local resolution range (Å)** | 1.9 -3.3 | 2.0 - 3.7 | 2.3 - 8.0 | 2.5 – 5.0 | 2.5 – 4.5 | 2.5 – 4.0 |
| **B-factor for map (Å<sup>2</sup>)** | 104 | 92 | 106 | 123 | 110 | 133 |
| **Non-hydrogen atoms** | 14,038 | 13,930 | 6,587 | 13,882 | 13,828 | 13,904 |
| **Residues** | 1,796 | 1794 | 859 | 1,796 | 1,796 | 1,796 |
| **Ligands** | 6 | 6 | 1 | 8 | 6 | 10 |
| **Bond lengths (Å)** | 0.003 | 0.003 | 0.003 | 0.002 | 0.002 | 0.002 |
| **Bond angle (°)** | 0.612 | 0.551 | 0.652 | 0.511 | 0.517 | 0.54 |
| **MolProbity score** | 1.62 | 1.64 | 1.64 | 1.75 | 1.64 | 1.82 |
| **Clashscore** | 7.49 | 6.73 | 9.69 | 8.53 | 7.64 | 6.44 |
| **Rotamer outliers (%)** | 0.06 | 0.00 | 0.00 | 0.00 | 0.00 | 0.00 |
| **Ramachandran favored (%)** | 96.65 | 96.20 | 97.32 | 95.81 | 96.54 | 92.63 |
| **Ramachandran allowed (%)** | 3.29 | 3.74 | 2.68 | 4.19 | 3.46 | 7.37 |
| **Ramachandran outliers (%)** | 0.06 | 0.06 | 0.00 | 0.00 | 0.00 | 0.00 |
| **Number of water molecules** | 200 | 150 | 0 | 0 | 0 | 14 |
| **Number of Mg<sup>2+</sup> ions** | 6 | 6 | 1 | 6 | 6 | 8 |
| **Number of nucleotides** | 0 | 0 | 0 | 2 | 0 | 2 |

### Additional Data

**Supplementary Data 1 Extended multisequence alignment of all P-type ATPases (see separate .aln file).**

**Supplementary Data 2 Primers, plasmids, and strains used in this study (see separate .xlsx file).**

### Extended Data Movies

**Extended Data Movie 1. 360 degree view of the dimeric cryo-EM map of *E. coli* Mg^2+^ transporter MgtA.**

**Extended Data Movie 2. 360 degree view of the monomeric cryo-EM map of *E. coli* Mg^2+^ transporter MgtA.**

**Extended Data Movie 3. Morph between the structural model of a single subunit of the dimeric and monomeric *E. coli* Mg^2+^ transporter MgtA.**

**Extended Data Movie 4. MD simulation movie of the N-terminal tail of the *E. coli* Mg^2+^ transporter MgtA.**

**Extended Data Movie 5. MD simulation movie of the *E. coli* Mg^2+^ transporter MgtA showing the full dimer (in color) aligned to the cryo-EM structure (grey).**

**Extended Data Movie 6. Zoomed in MD simulation movie of the *E. coli* Mg^2+^ transporter MgtA dimer interface.**

**Extended Data Movie 7. The *E. coli* Mg^2+^ transporter MgtA dimer with ATP bound.**

**Extended Data Movie 8. MD simulation movie of the Mg^2+^ ion in the middle of the transmembrane domains of the *E. coli* Mg^2+^ transporter MgtA.**

